# Deciphering the oxidative modifications via disulfide mapping

**DOI:** 10.1101/2025.07.30.666989

**Authors:** Yeonjoo Lee, Tae-Kyung Kim, Seungjin Na, Kong-Joo Lee, Jaeho Jeong, Eun Joo Song

**Affiliations:** College of Pharmacy and Graduate School of Pharmaceutical Sciences, Ewha Womans University, Republic of Korea; CJ BIO Manufacturing, CJ CheilJedang, Suwon, Republic of Korea; Digital Omics Research Center, Korea Basic Science Institute, Cheongju, Republic of Korea; Graduate Program in Innovative Biomaterials Convergence, Ewha Womans University, Seoul, Republic of Korea

## Abstract

Oxidative stress triggers redox-sensitive post-translational modifications, notably disulfide bond formation involving cysteine residues. However, these bonds are often overlooked in proteomics due to the routine use of reducing agents. Here, we employed LC-MS-based metabolomics and non-reducing tandem mass tag (TMT) proteomics to investigate the effects of H₂O₂ on MDA-MB-231 cells. Metabolomic analysis revealed pathway-specific inhibition of major metabolic pathways including glycolysis, the TCA cycle, and nucleotide biosynthesis. Proteomic analysis using the DBond algorithm revealed extensive and isoform-specific disulfide crosslinks across more than 1,000 proteins. These linkages were enriched at redox-sensitive cysteines near basic residues and displayed high isoform specificity. Our findings demonstrate that disulfide bond formation serves as a selective mechanism of redox regulation. This study highlights the utility of non-reducing proteomics in elucidating redox-controlled protein networks and structural dynamics under oxidative stress.

**Teaser:** Non-reducing proteomics uncovers hidden disulfide-linked networks that rewire protein responses to oxidative stress.

## Introduction

Hydrogen peroxide (H₂O₂), a reactive oxygen species (ROS), is produced by non-phagocytic cells in response to various stimuli and functions as a second messenger regulating diverse cellular processes, including proliferation, differentiation, inflammation, stress responses, aging, and cell death(*1*). These effects are primarily mediated by redox modifications of cysteine (Cys) residues in regulatory proteins such as PTEN, protein tyrosine phosphatases, peroxiredoxins (PRDXs), Hsp33, and EGFR(*1-4*). Cys oxidation serves as a molecular switch, altering protein structure and function, with redox sensitivity influenced by the local microenvironment and pKa(*5-7*).

Cysteine thiols (Cys–SH) can undergo reversible or irreversible oxidation, forming sulfenic acid (Cys–SOH), disulfide bonds (Cys–S–S–Cys), sulfinic acid (Cys–SO₂H), and sulfonic acid (Cys– SO₃H)(*8-10*). These oxidative forms participate in signaling and metabolic control in a concentration-dependent manner. While low levels of H₂O₂ (<100 μM) support physiological signaling, higher levels (>200 μM) can induce oxidative stress or apoptosis. Among these modifications, disulfide bonds are particularly important due to their reversible nature and structural specificity. However, they are often overlooked in proteomics workflows that use reducing agents like DTT or β-mercaptoethanol(*11, 12*).

Many redox-regulated proteins—including GAPDH, PRDX1, DJ-1, and Nm23-H1—undergo disulfide bonding that alters their activity, interactions, or structural conformation (*5-7, 13-16*). For example, Nm23-H1 forms an intramolecular disulfide bond upon ROS exposure, while its isoform Nm23-H2 lacks key Cys residues and is unresponsive to oxidation (*5, 6*). These examples highlight how isoform-specific Cys content contributes to redox selectivity. Selective thiol labeling using NPSB-1 has demonstrated that cysteine reactivity varies depending on local structure and isoform context(*12*).

Despite their functional relevance, disulfide bonds are difficult to detect with conventional proteomic methods due to reduction during sample preparation. To address this, we implemented a non-reducing tandem mass tag (TMT)-based proteomics workflow combined with LC-MS-based metabolomics in H₂O₂-treated MDA-MB-231 breast cancer cells(*7*). Non-reducing SDS-PAGE preserved endogenous disulfide bonds and oxidation states, enabling isoform-level analysis of redox-sensitive proteins.

A key component of this strategy is the DBond algorithm, which enables precise identification of disulfide-linked peptides directly from tandem mass spectra(*11*). DBond incorporates scoring rules based on cleavage patterns, mass shifts, and fragmentation features unique to disulfide-linked peptides. This allows confident detection of intra- and interprotein disulfide bonds, including those that are isoform- or site-specific, which would be otherwise undetectable using conventional peptide search engines.

Metabolomics revealed H₂O₂-induced inhibition of glycolysis, the tricarboxylic acid (TCA) cycle, and nucleotide biosynthesis, while proteomics uncovered widespread disulfide crosslinking. These disulfide linkages were enriched at basic residue regions and showed high isoform specificity. Collectively, these results demonstrate that disulfide bonding is a selective and structurally guided mechanism of redox regulation, and that DBond-enabled non-reducing proteomics is a powerful tool for mapping oxidative stress–responsive protein networks.

## Results

To assess cellular responses to varying oxidative stress levels, we performed metabolomic and proteomic analyses on cells treated with 0.1–1 mM H_2_O_2_ (Fig. 1), using standard workflows for each omics platform.

**Fig. 1.**
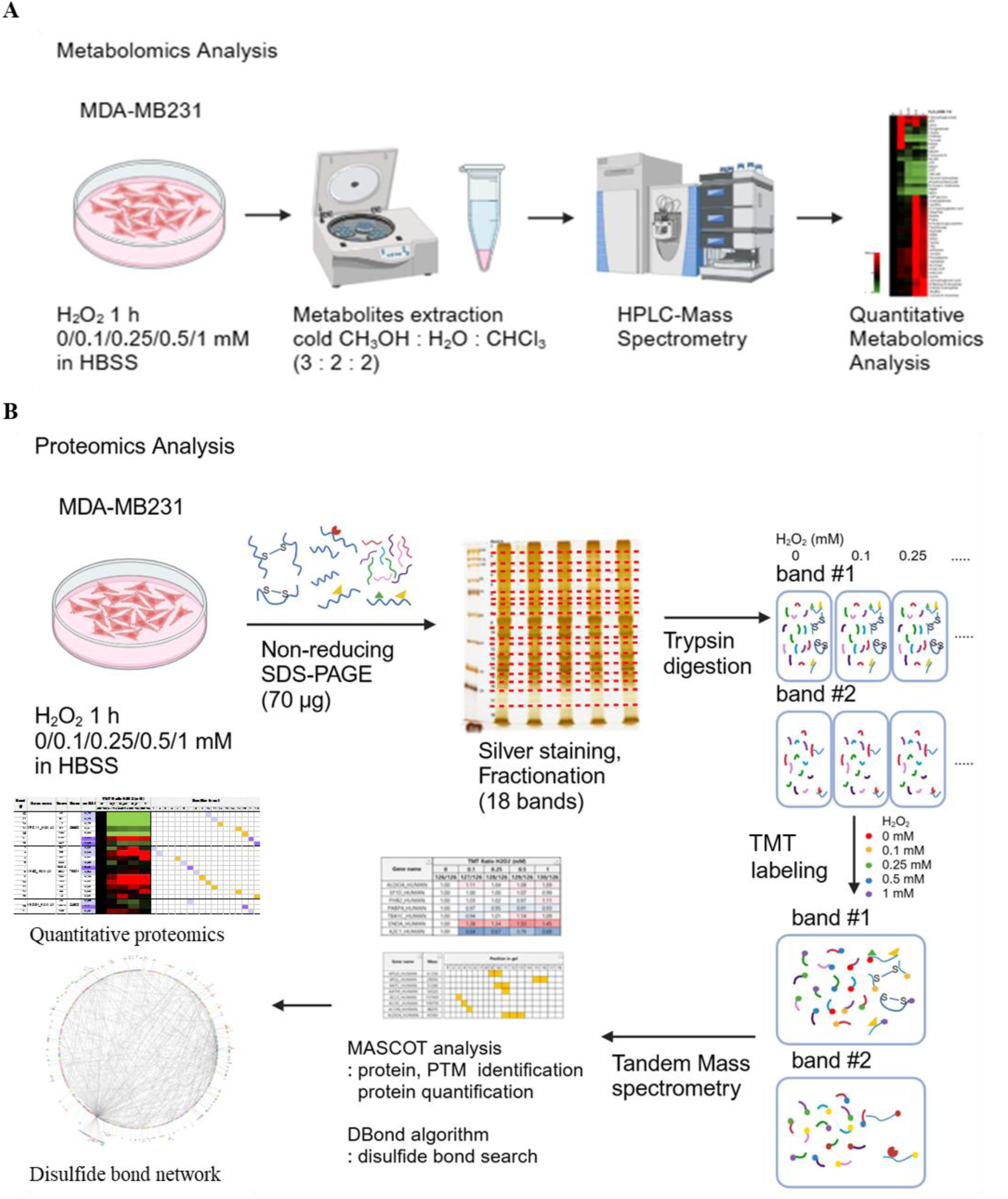
Schematic overview of the metabolomics and proteomics workflow. MDA-MB-231 cells were treated with various concentrations of H_2_O_2_ before processing for proteomics and metabolomics analyses. *n = 3* for proteomics, *n = 5* for metabolomics. **(A)** Metabolomics workflow. Following H_2_O_2_ treatment, hydrophilic metabolites were extracted, before being analyzed using HPLC-MS. **(B)** Proteomics workflow. Cell lysates were prepared in a non-reducing sample buffer following H_2_O_2_ exposure. Different TMT reagents were used to label samples corresponding to each H_2_O_2_ concentration.

### Redox-sensitive pathways identified by metabolomics in oxidative stress

To assess concentration-dependent metabolic responses to oxidative stress, we performed LC-MS–based metabolomics on MDA-MB-231 cells treated with 0–1 mM H_2_O_2_ for 1 h. Metabolites were extracted using cold methanol–water–chloroform (3:2:2) and analyzed by HPLC-MS, with five replicates per condition. A total of 79 intracellular metabolites were quantified. Heatmap and PCA analysis confirmed clear separation of samples based on H_2_O_2_ concentration (Fig. 2A–B).

**Fig. 2.**
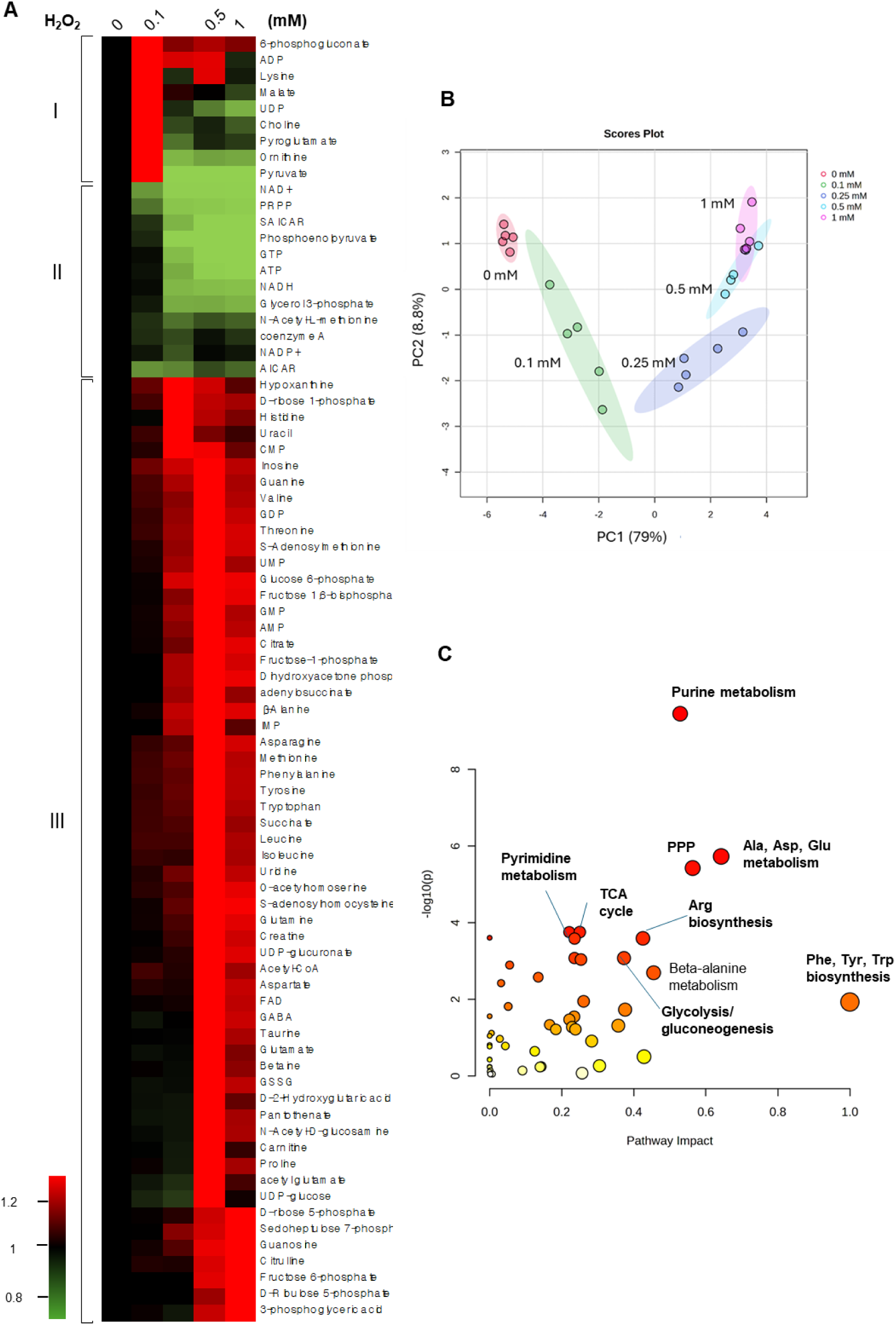
Metabolite changes under oxidative stress conditions. **(A)** Heatmap showing metabolite changes across H_2_O_2_ concentrations. Metabolites were categorized into three groups based on their response patterns. **(B)** PCA of metabolite profiles for H_2_O_2_ concentration. **(C)** Pathway analysis of redox-sensitive metabolic pathways using MetaboAnalyst 6.0. Only metabolites exhibiting >20% change at 1 mM H_2_O_2_ were included. fig. S1B illustrates detailed values and the full list of affected pathways.

Metabolites were classified into three groups. Group I (e.g., 6-phosphogluconate, ADP, UDP, pyruvate) showed a transient increase at 0.1 mM, followed by decline. Group II (e.g., AICAR (5-Aminoimidazole-4-carboxyamide ribonucleoside), SAICAR (succinyl-AICAR), ATP, GTP, PRPP (phosphoribosyl pyrophosphate)) decreased progressively. Group III (58 metabolites) increased dose-dependently. Pathway analysis of >20% changing metabolites at 1 mM H_2_O_2_ revealed oxidative sensitivity in purine/pyrimidine metabolism, the pentose phosphate pathway (PPP), the tricarboxylic acid (TCA) cycle, glycolysis/gluconeogenesis, and amino acid metabolism (Fig. 2C, fig. S1B).

Glycolysis showed early metabolite accumulation (G6P, DHAP) with downstream depletion, suggesting redox-sensitive blockades (fig. S2A). In the TCA cycle, citrate and succinate increased, while malate decreased, implying disrupted flux due to altered NAD⁺/FAD (fig. S2B). The PPP showed enhanced NADPH production but GSSG accumulation, suggesting NADPH depletion (fig. S2C-D). Nucleotide pathways showed precursor depletion and imbalance (fig. S2E-F), and many amino acids peaked at moderate ROS, reflecting inhibited translation (fig. S2G). These changes reflect targeted metabolic rewiring under oxidative stress.

### Proteomic analysis of protein diversity under non-reducing conditions

Since cellular metabolites and pathways responded differently to oxidative stress, we examined redox-sensitive proteomic changes under non-reducing conditions across varying H_2_O_2_ concentrations (0–1 mM). MDA-MB-231 cells were treated for 1 h, and proteins were separated by non-reducing PAGE, followed by silver staining (Fig. 1, fig. S3). Gels were divided into 18 bands, digested, labeled with 6 TMT tags, and quantified via TMT mass spectrometry. Proteins were filtered based on peptide scores (≥20), ≥2 unique peptides, and standard deviations (≤1.3). Post-translational modifications (PTMs) and disulfide crosslinks were identified using MASCOT and DBond, respectively(*11*). A total of 1,669 protein bands corresponding to 1,062 proteins were quantified, including data on band positions, exponentially modified Protein Abundance Index (emPAI), TMT ratios, disulfide-linked cysteines, and PTMs.

Among the identified proteins, 14 appeared across ≥9 bands and 83 across 3–8 bands (fig. S4). These proteins included peroxiredoxin 1 (PRDX1), which formed high-molecular-weight oligomers under oxidative stress(*17*), and proteins with distributed emPAI values such as keratins, and glyceraldehyde-3-phosphate dehydrogenase (GAPDH) with one dominating emPAI values (*18*).

To validate the MS findings, Western blotting was conducted for six proteins (Fig. 3). Each showed multiple bands under non-reducing conditions, with primary bands being relatively unresponsive and secondary bands highly sensitive to H_2_O_2_. These secondary bands were often detected only in MS due to low abundance (emPAI <1% of the primary) and required higher lysate input for Western detection. PRDX1 upper bands decreased ∼70% at 0.5 mM H_2_O_2_, while the main band increased 1.5-fold at 0.25 mM. Western confirmed PRDX1 dimers, which diminished as H_2_O_2_ increased, supporting disulfide-dependent regulation.

**Figure 3.**
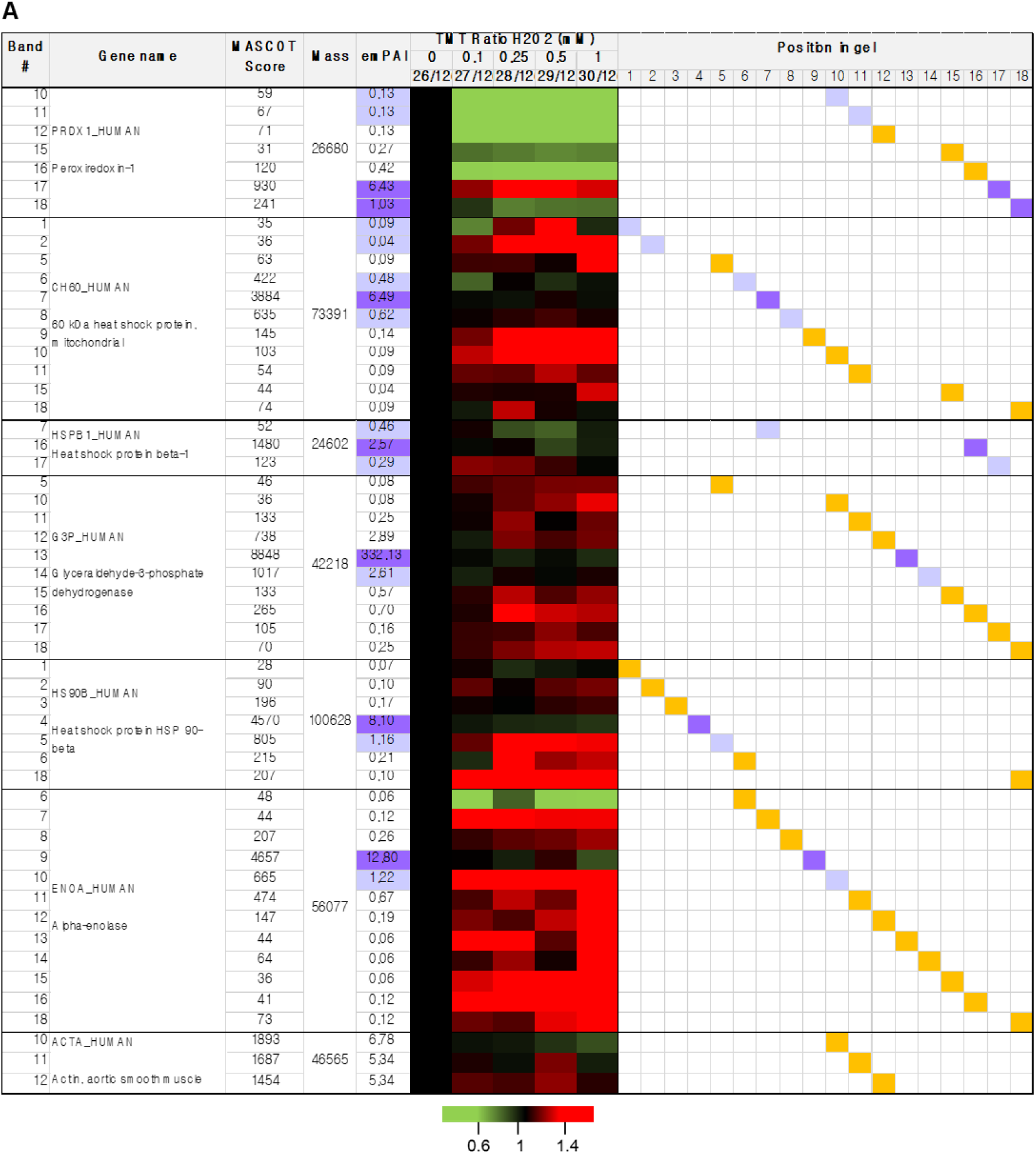
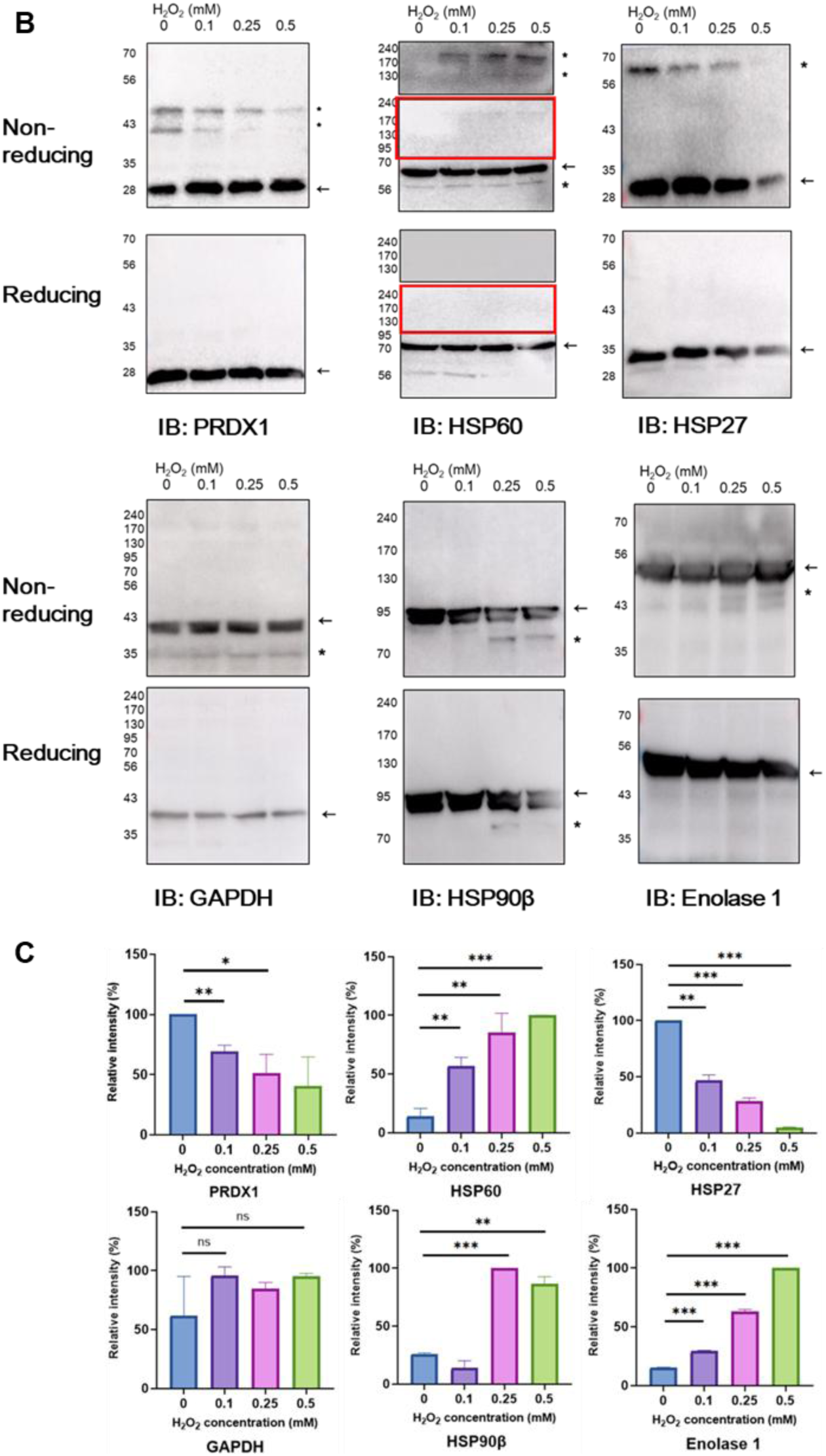
Western blot validation of mass spectrometry results. **(A)** Selection of proteins with significant abundance changes for validation. TMT ratios ≤0.6 are shown in green, and ratios ≥1.4 are shown in red. Main Western blot bands are marked in purple and secondary bands in pale purple. ACTA_HUMAN serves as the negative control. **(B)** Western blot analysis of proteins in panel A under non-reducing and reducing conditions. Equal amounts of lysate and identical exposure times were maintained. Arrows indicate main bands (purple), and asterisks indicate secondary bands (pale purple) corresponding to each protein in pane A. **(C)** Quantification of Western blot bands shown in panel B. Band intensities were normalized to the 0 mM H₂O₂ condition for each protein and plotted as relative intensity (%). Data represent mean ± standard deviation (n = 3). *p < 0.05; **p < 0.01; ***p < 0.001; ns = not significant.

Similarly, heat shock protein 60 (HSP60/CH60) showed ≥1.4-fold increases in upper bands at ≥0.25 mM H_2_O_2_, with minimal change in the dominant band. Detection of secondary bands required increased lysate loading. Heat shock protein beta-1 (HSP27/HSPB1) upper bands decreased (0.81–0.87-fold) at 0.5 mM, while primary bands were largely unchanged. GAPDH upper bands increased up to 1.4-fold at 1 mM H_2_O_2_ but were undetectable by Western due to the large emPAI value of primary band (332.13). Heat shock protein 90 beta (HS90B) lower bands increased 1.7-fold at 0.25 mM but were detected regardless of redox conditions, suggesting a disulfide-independent mechanism. Enolase-1 (ENOA) showed up to 1.5-fold increases in secondary bands at 0.5 mM H_2_O_2_, consistent with disulfide-dependent Western results.

In conclusion, MS provided higher sensitivity than Western for detecting redox-induced changes in low-abundance protein subpopulations. The redox-responsive secondary bands, often invisible in Western blotting, suggest that minor protein populations undergo disulfide-mediated regulation under oxidative stress.

### Secondary protein bands reflect oxidative stress–induced post-translational modifications

Multiple secondary bands and their redox-responsive changes were commonly observed across proteins. To explore the origin of these bands, we analyzed PTMs using MASCOT-based MS/MS spectra identification. A detailed proteomic comparison between primary and secondary bands was conducted to assess PTM patterns and redox-induced abundance shifts (Fig. 3).

In total, 13,107 peptides from 1,511 proteins were identified, with 6,604 peptides (from 858 proteins) showing MASCOT scores >30 (table S2). PTMs included ubiquitination (GlyGly) and acetylation on Lys, phosphorylation on Ser/Thr/Tyr, deamidation on Asn/Gln, oxidation on Met, and several Cys modifications such as propionamide adducts, dehydroalanine (DHA), and trioxidation— indicative of irreversible oxidation(*10, 11*). Among proteins with >10 bands, eight were cytoskeletal (e.g., K1C10, K1C9, K2C1, ACTG, TBA1B, TBB5), while others included HSP60, ENOA, GAPDH, EF1A1, and ubiquitin (fig. S4; table S1, 2).

These secondary bands often had emPAI values <1.0—up to 100-fold lower than the primary band—indicating lower abundance despite rich PTM profiles(*18*). Nonetheless, their TMT ratio changes under oxidative stress mirrored those of the main bands (table S1, 2). For instance, GAPDH showed a primary emPAI of 332.13 at ∼36 kDa. Its peptide ^146^IISNASCTTNCLAPLAK^162^ appeared in multiple bands, exhibiting DHA and trioxidation at Cys152/156 (suggesting intramolecular disulfide bonding), phosphorylation at Ser148/151, and Lys162 acetylation (table S2).

Cytoskeletal proteins also displayed varied PTMs across multiple bands. Unlike GAPDH, which concentrated in a single main band, cytoskeletal proteins (e.g., K2C1, K1C9) were evenly distributed with maximum emPAI (∼1.1), showing broad band patterns with distinct PTM combinations and TMT shifts (fig. S5, 6). These results suggest keratins exist in multiple stable forms, each characterized by unique PTM signatures.

Together, these findings indicate that secondary bands represent distinct protein populations generated by diverse PTMs. Further investigation is needed to determine their biological significance.

### TMT-based classification of redox-sensitive protein isoforms

To classify redox-sensitive proteins, we analyzed TMT ratio changes across 1,669 bands from 1,062 proteins using nanoLC-MS/MS, with 0 mM H_2_O_2_ as the control. Proteins were grouped based on the band showing the most significant TMT ratio change (≤0.875 or ≥1.25) at the lowest H_2_O_2_ concentration (Fig. 4A). Group I (180 proteins), including PRDX1 and HSP60, responded at 0.1 mM H_2_O_2_; Group II (80), including GAPDH and PPIA, at 0.25 mM; Group III (88), including ROA2, at 0.5 mM; Group IV (86), including 14-3-3 beta/alpha, at 1 mM. Group V (628 proteins), including PDIA6 and ARPC3, showed no change across all conditions (table S3).

**Fig. 4.**
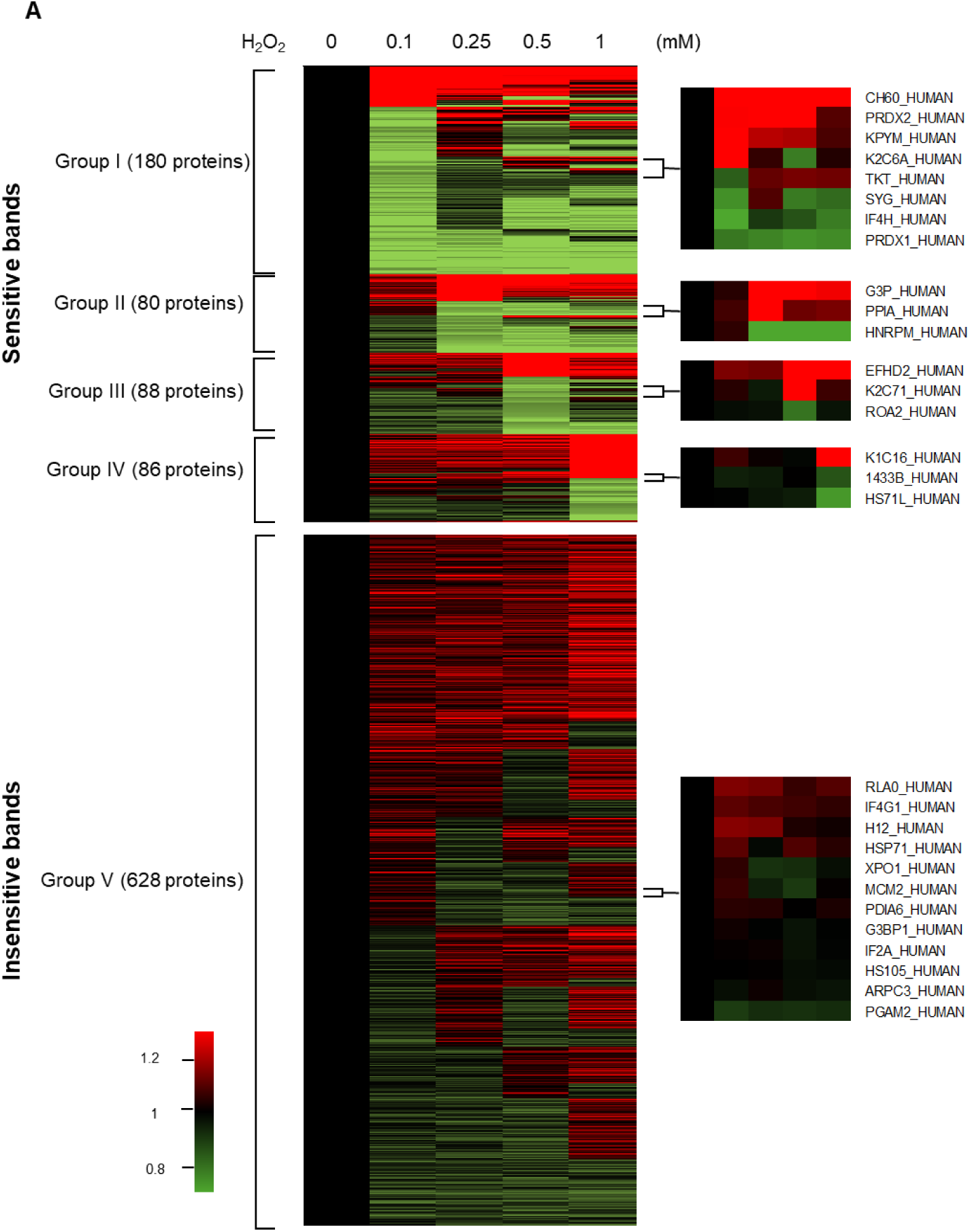
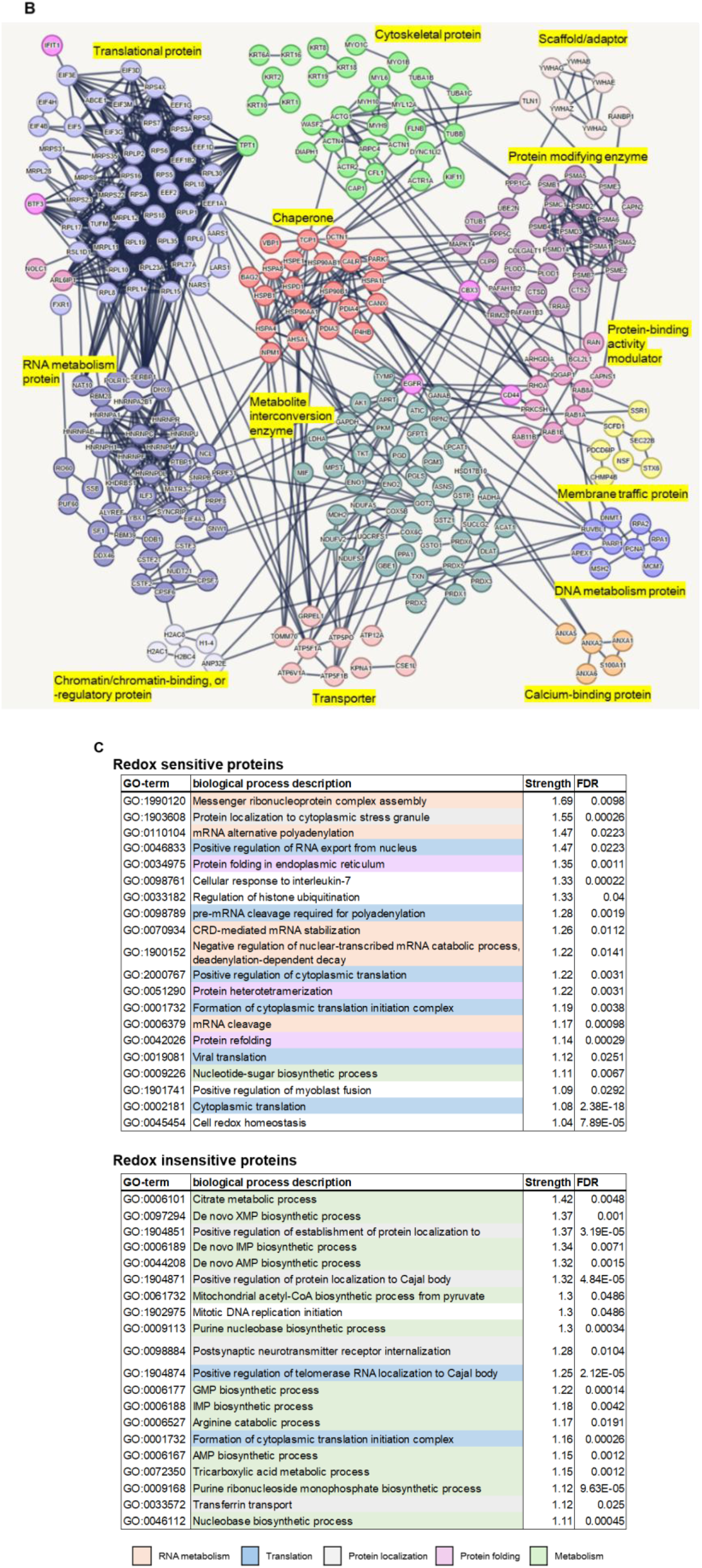
Classification and network analysis of redox-sensitive proteins. **(A)** Heatmap of redox-sensitive and redox-insensitive proteins. Proteins were classified into five groups based on the concentration at which the band intensity changed by ≤0.875 or ≥1.25. TMT ratios ≤0.8 are shown in green and ≥1.2 in red. table S3 presents detailed TMT ratio values and group assignments. **(B)** STRING network of 434 proteins with TMT ratios ≤0.875 or ≥1.25 at any H_2_O_2_ concentration. Proteins within the same PANTHER protein class were clustered and color-coded accordingly. Edge thickness indicates the confidence level of predicted protein–protein interactions; only interactions with the highest confidence score (≥0.900) are shown. **(C)** Top 20 biological processes enriched in redox-sensitive and redox-insensitive proteins; based on STRING analysis. Related processes are color-coded: RNA metabolism (orange), protein localization (gray), translation (blue), protein folding (purple), and metabolism (green).

We further analyzed the 434 redox-sensitive proteins (Groups I–IV) using STRING (v12.0; http://string-db.org/) and PANTHER classifications (Fig. 4B). These proteins spanned diverse classes, including chaperones, cytoskeletal proteins, chromatin regulators, RNA/DNA metabolism proteins, translation-related proteins, modifying enzymes, transporters, and adaptors. In contrast, redox-insensitive proteins in Group V showed no significant class distribution differences (data not shown). Notably, even within the same complexes, such as ribosomes, individual proteins exhibited distinct TMT ratio changes (fig. S7), reflecting heterogeneous redox responses.

Functional enrichment analysis using STRING revealed that redox-sensitive proteins were predominantly involved in RNA metabolism, translation, protein folding, localization, and metabolism (Fig. 4C). These processes likely reflect rapid responses to oxidative damage, facilitating turnover and repair. In contrast, redox-insensitive proteins were primarily associated with general metabolic functions. This discrepancy suggests that oxidative stress triggers coordinated changes in redox-sensitive pathways, while metabolic responses are limited to specific enzymes, as also indicated by our metabolomics data. Although metabolism-related pathways were enriched among the insensitive group, key enzymes like PRDX1, GAPDH, and enolase exhibited diverse TMT ratio shifts and PTMs across secondary bands (Fig. 3A), indicating selective regulation by redox state.

### Redox-sensitive enzyme modulation disrupts metabolic pathways under oxidative stress

To validate the effects of oxidative stress on cellular metabolism, we integrated proteomic data on redox-sensitive enzymes with metabolic pathway maps (fig. S8, table S4). Instead of relying on raw data, we highlighted key enzymes with altered abundance or PTMs under stress.

Previously, we observed a reversed ratio of glyceraldehyde 3-phosphate (G3P) and 3-phosphoglycerate (3-PG) under oxidative stress (fig. S2A), implicating GAPDH and phosphoglycerate kinase 1 (PGK1). Although both act sequentially in glycolysis, their proteomic responses differed. PGK1 showed minimal changes, while GAPDH appeared in 10 bands, with secondary bands increasing up to 1.5-fold. GAPDH displayed diverse PTMs—Cys oxidation, phosphorylation, acetylation, and ubiquitination—suggesting redox-mediated structural diversity. Other glycolytic enzymes (e.g., PGAM, enolase, pyruvate kinase) also showed dynamic changes in TMT ratios and PTMs (fig. S8).

TCA cycle enzymes responded variably. Succinate dehydrogenase (SDH) increased up to 1.3-fold and showed Cys oxidation. Malate dehydrogenase (MDH) and citrate synthase (CISY) also changed, corresponding to altered metabolite levels, such as citrate and succinate accumulation and malate and ATP depletion (fig. S2B). These changes suggest that oxidative stress disrupts TCA cycle flux via quantitative and PTM-based enzyme regulation.

The oxidative branch of the pentose phosphate pathway (PPP), crucial for NADPH production, was disrupted, with enzymes like 6-phosphogluconolactonase (6PGL) and 6-phosphogluconate dehydrogenase (6PGD) showing significant TMT ratio changes. Despite stable NADPH levels, increased GSSG indicated insufficient reducing power (fig. S2D). The non-oxidative PPP enzymes, including transketolase (TKT) and transaldolase (TAL), remained unchanged.

In nucleotide biosynthesis, PRPS1, involved in phosphoribosyl pyrophosphate (PRPP) production, increased by >20% at 1 mM H_2_O_2_ and exhibited PTMs such as Cys oxidation and phosphorylation. ATIC showed a >20% reduction in upper band intensity under stress, indicating redox-modulated activity. Nucleoside diphosphate kinase-B (NDPK-B) remained stable, while NDPK-A is known to lose activity through oxidative dissociation(*5, 6*).

Most amino acids accumulated under oxidative stress, peaking at 0.5 mM H_2_O_2_. Histidine peaked earlier at 0.25 mM before declining; lysine remained unchanged. These patterns likely reflect translational inhibition, confirmed by puromycin incorporation assays showing dose-dependent repression (fig. S9), contributing to amino acid accumulation (fig. S2G).

Together, our findings highlight that oxidative stress alters metabolism through both quantitative changes and functionally relevant PTMs in enzymes. GAPDH, SDH, and oxidative PPP enzymes emerge as central redox-sensitive regulators. For detailed data, see table S4 and fig. S8.

### Identification of intra- and inter-disulfide crosslinked proteins

To identify disulfide-bonded proteins and associated Cys residues, we analyzed MS/MS data from proteins prepared under non-reducing conditions (Fig. 1). Disulfide-linked peptides were detected using the DBond algorithm, which identifies crosslinked peptides based on fragmentation patterns and PTMs(*11*). A total of 68,038 disulfide-linked peptides were identified, corresponding to 1,287 unique protein pairs (table S5).

To ensure reliability, peptides with low DBond scores (<3.5), short sequences (<6 residues), or ambiguous origins were filtered using the neXtProt Peptide Uniqueness Checker (http://nextprot.cn/tools/peptide-uniqueness-checker), yielding 21,059 high-confidence disulfide bonds from 1,185 proteins and 16,666 unique Cys site pairs. Representative MS/MS spectra of disulfide-linked peptides are shown in fig. S10. Actin, for example, formed high-confidence disulfide crosslinks with multiple proteins such as pyruvate kinase and vimentin, which were validated by immunoprecipitation under non-reducing conditions (fig. S11A–B).

Among these, 3,483 disulfide crosslinks (involving 2,856 site pairs across 1,006 proteins) were associated with shifts to higher molecular weight bands than their canonical counterparts, suggesting that inter-disulfide bonding alters electrophoretic mobility under non-reducing conditions. These results provide a large-scale, previously underappreciated dataset of disulfide-linked proteins and their reactive Cys residues.

To explore the redox-regulated interaction network, a disulfide bond network was constructed using Cytoscape (v3.10.3; https://cytoscape.org)(19) (Fig. 5A). Only protein-specific peptides and bonds supported by at least three linkages per protein pair were included. The resulting network included 5,275 disulfide bonds connecting 1,064 protein pairs among 637 proteins. Louvain clustering via ClusterMaker2 revealed modular subclusters based on connection density. Some subclusters (Fig. 5B) showed Gene Ontology (GO) enrichment, suggesting functional relatedness, while others (Fig. 5C) lacked significant GO terms, indicating novel or uncharacterized redox-regulated protein groups.

**Fig. 5.**
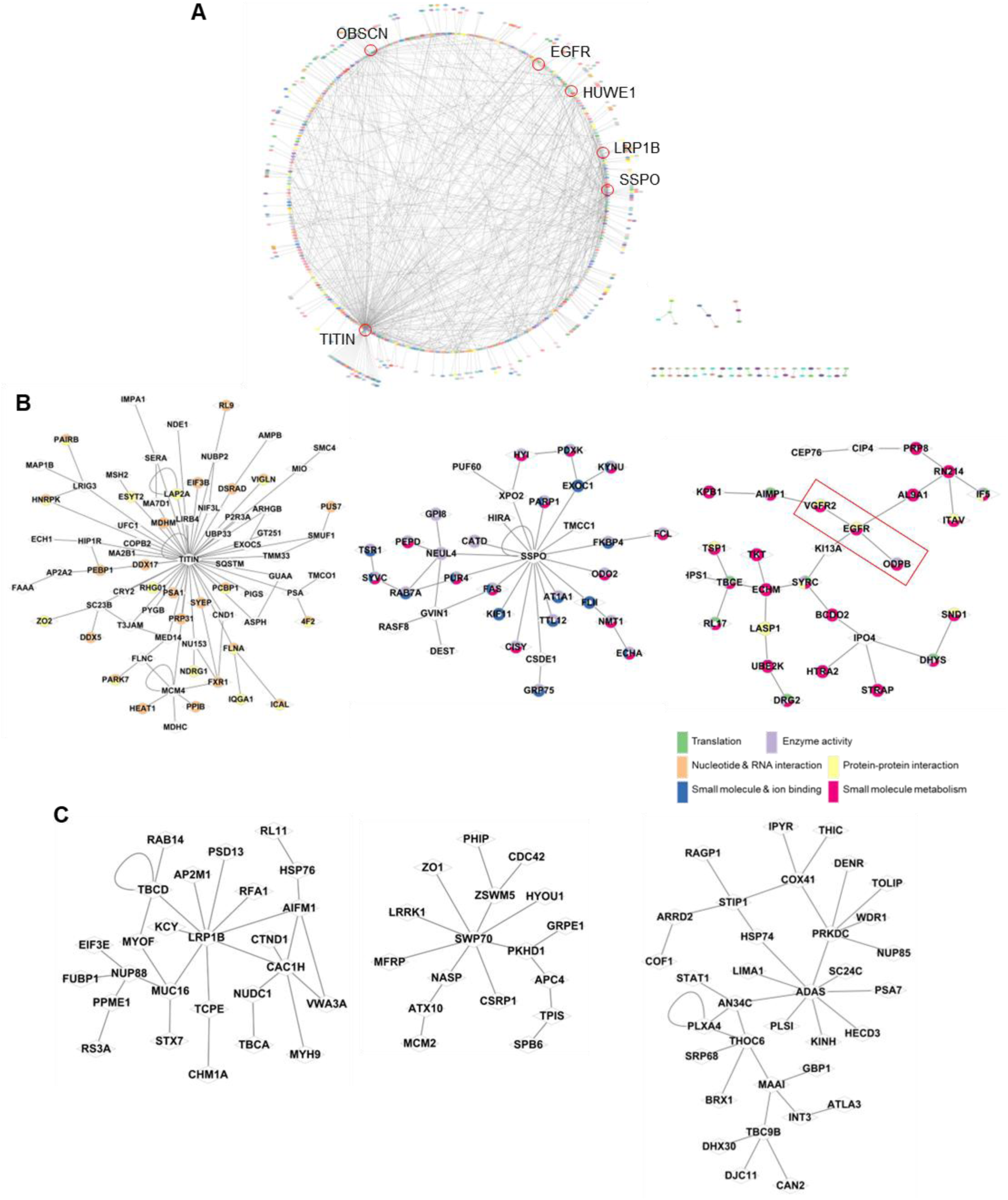
Disulfide bond network. Global disulfide bond network among proteins. Disulfide linkages were identified using the DBond algorithm and visualized in Cytoscape. Only disulfide bonds observed in at least three independent peptides and not derived from isoform-conserved sequences were included. Each node represents a distinct protein, and each node indicates recurring disulfide linkages observed three or more times. **(B–C)** Representative subclusters derived from multilevel clustering analysis. Subclusters containing proteins with known STRING interactions are shown in **(B)**, while those without shared STRING interactions are shown in **(C)**.

Overall, this analysis identified over 21,000 high-confidence disulfide bonds in more than 1,100 proteins. The disulfide network illustrates widespread and functionally diverse redox-driven interactions that have been largely missed under conventional reducing conditions, underscoring the structural and regulatory roles of disulfide bonding under oxidative stress.

### Specificity of disulfide bindings between proteins

To evaluate whether disulfide bond formation is specific and regulated, we analyzed structural and sequence features surrounding Cys residues using our high-confidence disulfide bond dataset.

Among 15,752 Cys residues in 1,163 proteins with DBond scores ≥3.5, only 32.7% (5,137) participated in disulfide bonding (Fig. 6A, table S6). This selective pattern was consistent across protein sizes. For example, proteins with ≤5 Cys residues often showed >50% disulfide usage, whereas in proteins with >50 Cys residues, only 3.0% reached this threshold, indicating that disulfide bond formation is not proportional to Cys content or protein size (Fig. 6B). Even large proteins like titin (TITIN), SCO-spondin (SSPO), and stabilin-2 (STAB2) contained many Cys residues, yet only a small subset formed disulfide bonds. Similarly, large proteins with few disulfide-linked Cys residues, such as UBR4 and DNHD1, supported this selectivity.

**Fig. 6.**
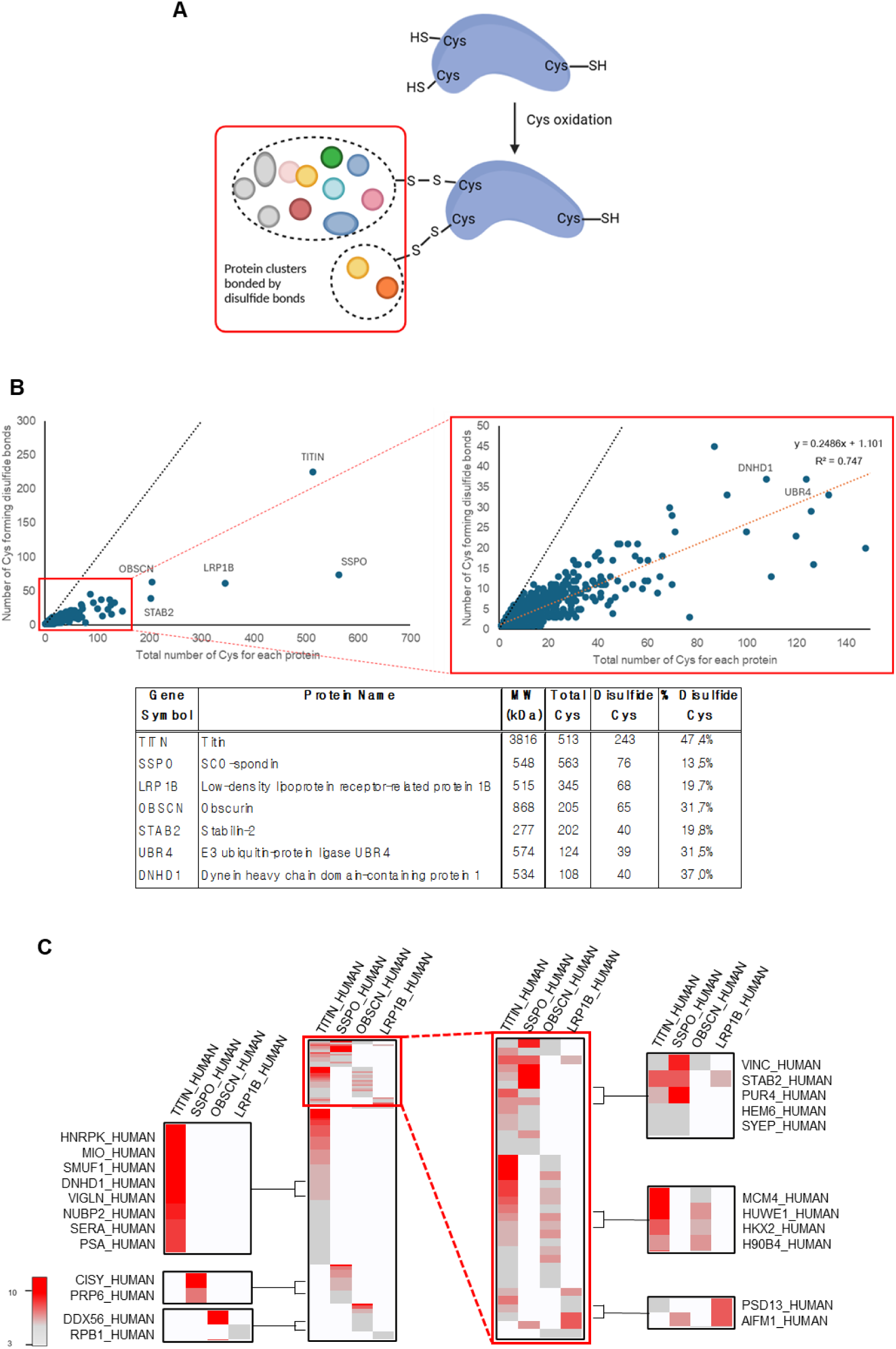
Features of Cys residues in disulfide-bonded proteins. **(A)** Schematic diagram showing subsets of Cys residues within individual proteins. Protein clusters sharing the same Cys binding sites were grouped using dotted lines. **(B)** Number of Cys residues forming disulfide bonds relative to the total number of Cys residues per protein. Protein names are indicated for those >200 total Cys residues. table S5 summarizes detailed values and protein names. **(C)** Heatmap showing proteins that interact with four large proteins (e.g., TITIN, SSPO, OBSCN, and LRP1B), each containing > 50 disulfide-bonded Cys residues. To reduce non-specific interactions, only proteins with at least three recurrent binding events were included.

To assess functional specificity, we focused on four large proteins (TITIN, SSPO, OBSCN, LRP1B), each with >50 disulfide-linked Cys residues. Among 726 interacting proteins with ≥3 DBond-supported links, distinct disulfide bonding partners were observed for each, indicating selective network integration (Fig. 6C).

Notably, disulfide crosslinks also varied across closely related isoforms. Using only high-confidence interactions (DBond score ≥3.5), we found that isoforms with highly conserved sequences often exhibited unique disulfide-linked partners (Fig. 7, fig. S13). For instance, the 14-3-3 isoforms (ε, σ, θ, ζ) shared overall structure and sequence, yet formed isoform-specific disulfide bonds. While both 14-3-3θ and ζ shared Cys25, 14-3-3θ crosslinked with DIAP1, whereas 14-3-3ζ interacted with α-aldolase, integrin β2, and TNFα-induced protein 8. The σ isoform used its unique Cys38 to link with Rab-5A and ESYT2, while Cys94 was primarily reactive in 14-3-3ζ (Fig. 7A), highlighting distinct redox networks despite structural similarity.

**Fig. 7.**
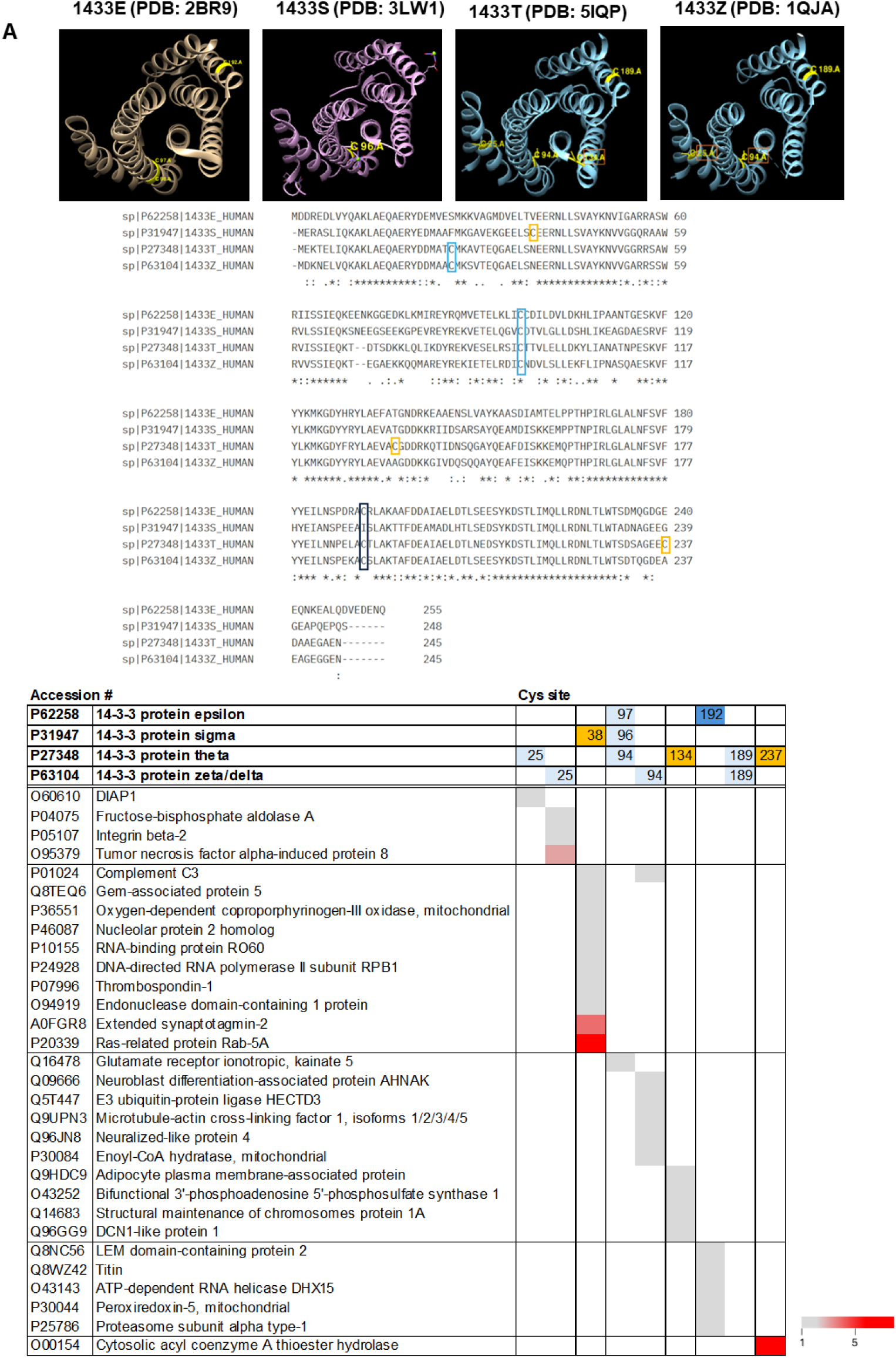
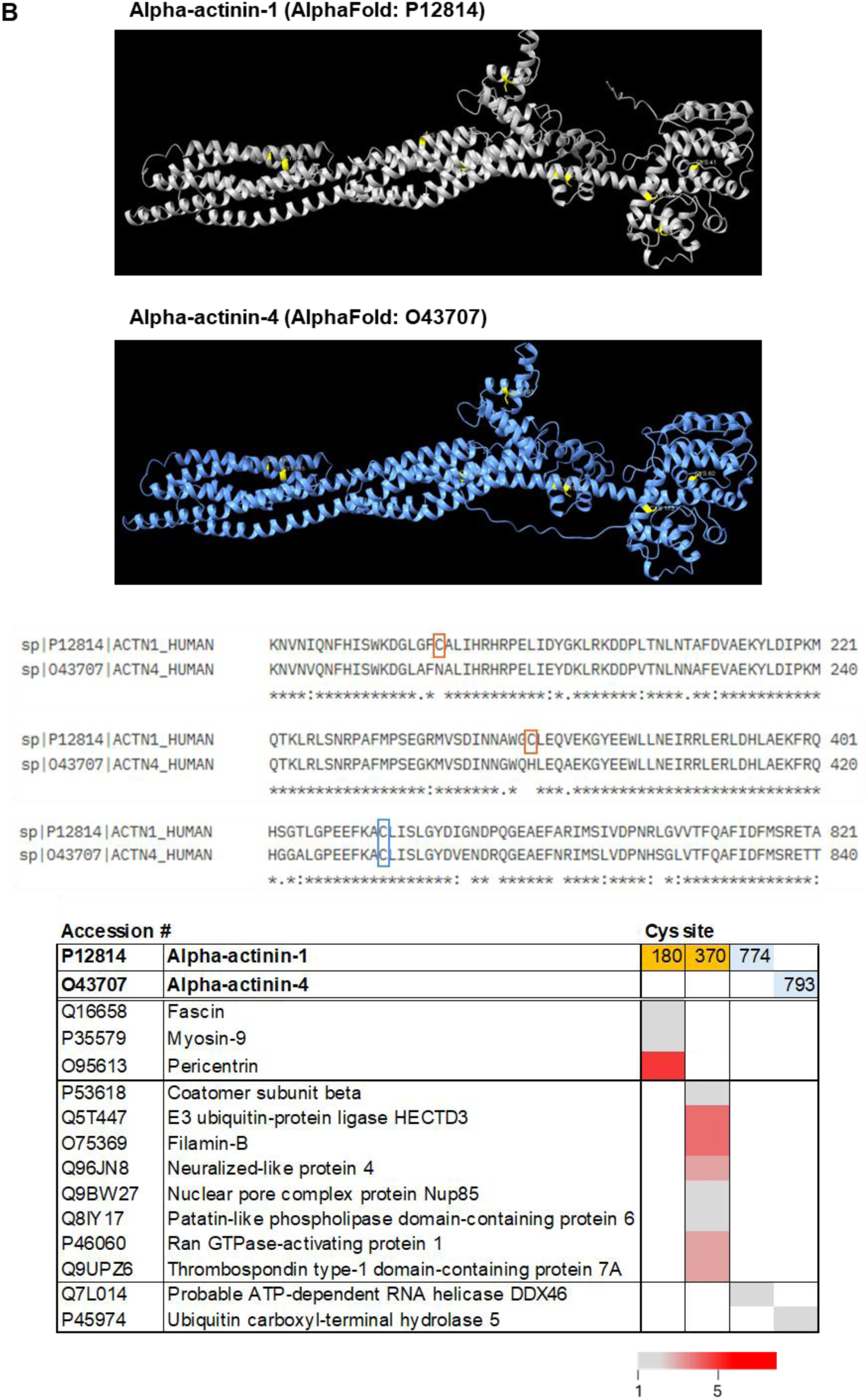
Homologous proteins with distinct disulfide bond clusters. Disulfide-bonded protein groups were compared between isoforms with high-sequence homology and similar structural features. **(A)** 14-3-3 proteins. **(B)** Alpha-actinin. Cys residues are shown in yellow on PDB or AlphaFold-predicted structures, with local sequence alignments provided. Binding partners for each Cys are listed in tables, with binding frequency color-coded: gray (1 detection) and red (≥5 detections). Only isoform-specific interactions with DBond scores ≥3.5 were included. In the alignments and tables, Cys residues were color-coded to reflect their distribution: isoform-specific Cys residues (orange); Cys residues shared by some isoforms (light blue); conserved Cys residues with biased disulfide linkage toward one isoform (dark blue). fig. S13 and table S5 depict additional examples.

Another example is alpha-actinin (ACTN), an actin crosslinking protein. Although ACTN1 and ACTN4 share 86% sequence identity and eight conserved Cys residues, only ACTN1-specific residues (Cys180 and Cys370) formed disulfide bonds with 11 unique proteins, including pericentrin (Fig. 7B, fig. S13B). This suggests disulfide bonds contribute to isoform-specific interaction profiles.

Enolase isoforms showed similar divergence. Enolase-γ, with a lower emPAI value (0.71), formed five or more disulfide bonds with choline-phosphate cytidylyltransferase A, a pattern not seen in the more abundant α (emPAI 12.80) or β (1.24) isoforms (fig. S13D). Similarly, Rab GDP dissociation inhibitor isoforms formed Cys302-dependent crosslinks with distinct RNA-binding proteins, including heterogeneous nuclear ribonucleoprotein L (fig. S13G).

Importantly, these isoform-specific disulfide networks did not correlate with protein abundance. For example, ACTN1/4 (1.51/1.90), ADT2/3 (0.39/0.28), and enolase γ (0.71) formed distinct disulfide bonds despite abundance similarity, suggesting structural context rather than expression level drives bonding specificity.

Together, these results indicate that disulfide bond formation is a selective, regulated process. Specific Cys residues act as redox-sensitive interaction hubs, enabling isoform-specific and functionally distinct protein networks. These findings suggest that disulfide bonds extend beyond structural roles to actively modulate protein interaction specificity and contribute to the functional diversification of isoforms.

### Characteristics of Cys residues forming disulfide crosslinks

To characterize the sequence and structural features of Cys residues involved in disulfide bond formation, we compared the local amino acid environments surrounding disulfide-forming and non-disulfide-forming Cys. To minimize artifacts from low-confidence interactions, we analyzed only those Cys involved in ≥3 disulfide bonds, identifying 2,930 Cys residues across 976 proteins—18.6% of the 15,752 total Cys sites (table S6). The remaining 12,822 Cys residues, either uninvolved or engaged in only 1–2 disulfide linkages, served as the comparison group.

Two key patterns emerged. First, positively charged amino acids—Arg and Lys—were enriched within ±10 residues of disulfide-forming Cys (Fig. 8A–C; fig. S14), particularly at the +1 and +2 positions. Arg appeared at 12.6% and 10.8%, and Lys at ∼8% in these positions—nearly double the frequencies near non-disulfide Cys (Arg: 4.1%; Lys: 4.6%). In contrast, histidine, glutamate, and aspartate showed minimal differences between the groups. A pattern search in >16,000 proteins (https://prix.hanyang.ac.kr/download/PatternMatchFrm.jsp) revealed 16,507 CK/R or CxK/R motifs across 1,547 proteins. However, only 22% were associated with disulfide formation, while 3,653 motifs from 937 proteins were identified in our dataset, suggesting selective involvement despite motif abundance.

**Fig. 8.**
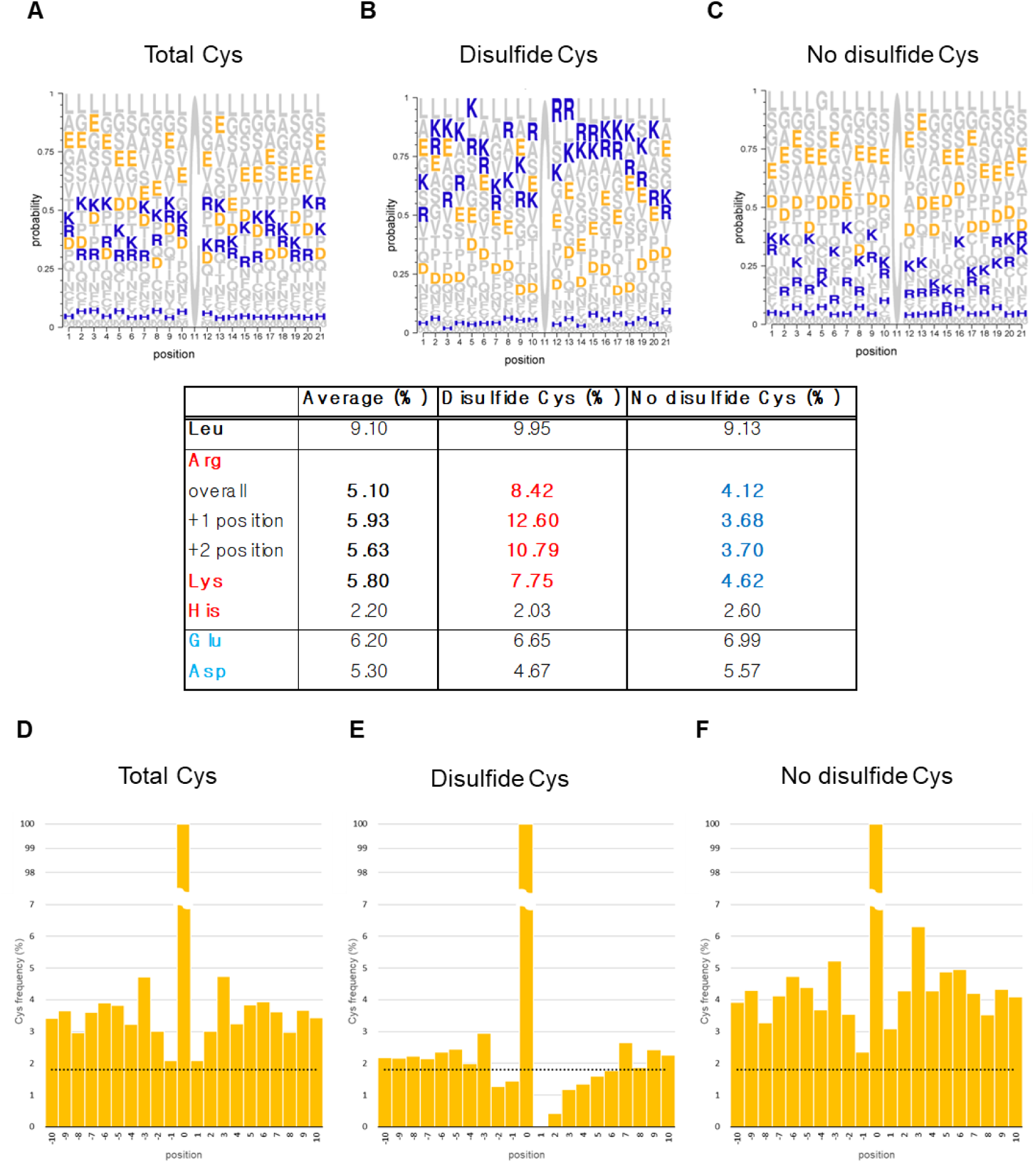
Amino acid frequencies near Cys residues. (A–C) Amino acid frequencies at positions ±10 around Cys residues, categorized as **(A)** all Cys residues, **(B)** disulfide-forming Cys residues, and **(C)** non-disulfide Cys residues. Positively charged amino acids (Lys, Arg) are shown in blue, while negatively charged amino acids (Asp, Glu) are shown in yellow. **(D–F)** Frequencies of Cys residues in each category. The dotted lines indicate the average proportion of Cys residues.

This enrichment of positively charged residues likely facilitates electrostatic stabilization of the thiolate anion, shifting the thiol–thiolate equilibrium toward the more reactive thiolate state and enhancing redox sensitivity.

Second, neighboring Cys residues were underrepresented around disulfide-forming sites, especially at the +1 and +2 positions, where Cys frequency ranged from 0% to 2.95%, compared to 2.35–6.30% near non-disulfide Cys (Fig. 8E–G). Pattern searches revealed 6,863 CC and 6,765 CxC motifs in the PRIX database, yet none of the CC and only 12 CxC motifs (from 10 proteins) were associated with disulfide bonds in our dataset. This suggests that adjacent Cys are spatially excluded from disulfide-forming regions, potentially to reduce steric hindrance or oxidative competition.

Together, these results identify two key features promoting disulfide bond formation: (1) electrostatic enrichment of Arg and Lys, especially at +1/+2 positions, and (2) exclusion of adjacent downstream Cys. These findings support a model in which disulfide formation is regulated by local sequence context and constrained by structural and functional considerations. Rather than occurring randomly, disulfide bond formation appears guided by sequence-driven stabilization and spatial selectivity, contributing to the efficiency and specificity of redox-based protein regulation.

## Discussion

In this study, we used non-reducing proteomics to identify redox-sensitive proteins and disulfide-linked networks, revealing how oxidative stress selectively modifies protein subpopulations and shapes their functions. Even minor protein subpopulations exhibited significant oxidative modifications under stress, highlighting the selectivity and compartmentalization of redox regulation. The identification of intra- and inter-protein disulfide bonds across a broad range of proteins provides a comprehensive map of disulfide networks across thousands of proteins. As demonstrated by reactive thiol probes such as NPSB-1, only a subset of cysteine residues exhibits reactivity under oxidative conditions(*12*). Isoform-specific disulfide formation, despite high sequence similarity, illustrates that disulfide bonding is a highly selective process shaped by local structural and contextual factors. Sequence analysis showed a strong enrichment of positively charged Lys/Arg residues near disulfide-prone cysteines, particularly at the +1/+2 positions, suggesting local charge influences thiolate formation and disulfide bonding(*20*).

These proteomic results align with metabolic findings showing that oxidative stress disrupts central metabolism such as glycolysis and the TCA cycle(*21*). Increased H_2_O_2_ concentrations led to accumulation of upstream glycolytic intermediates and reduced downstream products, redirecting flux toward the oxidative PPP to generate NADPH (fig. S2)(*22, 23*). Corresponding PTMs were found in PPP enzymes like G6PD and 6PGD(*24*), while GAPDH and SDH were inactivated via Cys oxidation(*25, 26*). Additionally, NDPK-A, another redox-sensitive metabolic enzyme, shifted from an active hexamer to an inactive dimer in an oxidation-dependent manner(*27, 28*). These redox induced modifications reinforce the connection between redox-sensitive PTMs and metabolic rewiring(*23*).

Though not directly detected, inverse abundance trends between uridine and uracil and between UMP and UDP suggest redox-related modulation of uridine phosphorylase and UMP kinase. The sharp decline in His may reflect reduced PRPP availability due to ATP depletion under oxidative stress(*29*). In contrast, Lys remained stable, likely due to redundancy in its biosynthetic pathways—via oxaloacetate/aspartate or α-ketoglutarate(*29*)—allowing compensation. Other amino acids with single biosynthetic routes were more vulnerable to redox stress. Arg, which is synthesized through the complex urea cycle, was undetected. Disulfide bond formation was a predominant Cys-based PTM enriched under oxidative conditions. GAPDH, PRDX1, and HSP60 showed multiple non-reducing SDS-PAGE bands, confirmed by MS/MS to represent redox-dependent PTM variants. These species were often absent under reducing conditions, underscoring the need to maintain disulfide integrity for accurate redox proteome profiling. Notably, low emPAI values and rich PTM diversity in secondary bands suggest that even minor redox-modified populations may exert distinct functional roles. Several “moonlighting” proteins such as GAPDH, PRDX1, and enolase exhibited extensive PTM remodeling and redox-driven functional shifts. GAPDH not only underwent disulfide bonding but also participated in RNA splicing complexes like p54nrb–PSF(*15, 25*). PRDX1 alternated between peroxidase and chaperone activities depending on redox state(*7, 30, 31*), while enolase 1 was implicated in mRNA stability regulation(*32, 33*). HSP60/HSP10 chaperonin complexes protected mitochondria from oxidative injury(*34*), and HSP27 formed Cys137-linked dimers associated with chemoresistance(*35*). Each of these proteins presented with multiple bands and PTMs, reinforcing the idea that low-abundance redox variants could hold specialized roles (Fig. 3; fig. S8). In contrast, most metabolic enzymes showed minimal banding and TMT changes, suggesting restricted redox responsiveness.

Our disulfide mapping also revealed novel disulfide-mediated inter-protein interactions. Examples include HSP70–*β*4-spectrin complexes and disulfide-linked apolipoproteins(*36*). STRING analysis revealed that while some subclusters shared known GO terms, others lacked annotated connections, implying previously uncharacterized interaction networks (Fig. 5B–C). Novel disulfide bridges between EGFR and vascular endothelial growth factor receptor 2 (VEGFR2) or pyruvate dehydrogenase E1 subunit beta (ODPB/PDHB) suggest redox-dependent crosstalk between signaling and metabolic pathways (Fig. 5B)(*37, 38*). Similarly, TITIN and SSPO—large, disulfide-rich proteins—formed networks enriched in cell cycle or structural assembly functions, indicating that disulfide bonding can organize coordinated functional modules (data not shown).

Isoform-specific disulfide formation, as seen in 14-3-3 proteins, reinforces the idea that protein-protein interactions and redox modifications are context-dependent, and not solely dictated by primary sequence (Fig. 7). While ACTN1 and ACTN4 colocalized at stress fibers and invadopodia, only ACTN4 regulated invadopodia formation and ER*α* activity(*39-42*). Disulfide-linked interactomes of 14-3-3 isoforms also differed, with unique Cys mediating distinct bonds (Fig. 7A)(*43-45*). Despite nearly identical local sequences, disulfide patterns diverged—suggesting cellular environment and PTMs, not just sequence, determine isoform specificity. This principle is demonstrated by Nm23-H1 and Nm23-H2, isoforms of NDPK. Nm23-H1 formed a stress-induced Cys4–Cys145 disulfide that triggered hexamer dissociation and inactivation, whereas Nm23-H2, lacking Cys4, remained unaffected(*5, 6, 27, 46*).

Although AlphaFold and crystal structures provide static insights, they fail to capture these redox-specific differences(*47*). This highlights that dynamic redox environments and PTMs are crucial determinants of disulfide formation and function. Our sequence analysis further supports this, showing selective enrichment of basic residues near redox-prone Cys residues (Fig. 8A)(*20*). However, not all CxK/R motifs formed disulfides, indicating additional layers of regulation.

Despite the scope of this work, several limitations should be acknowledged. The peptide-level ambiguity in disulfide assignment―particularly for sequences shared among homologs―posed challenges in protein-specific interpretation. For example, ERACYLSINPQK was shared by *α*-/*β*-centractin, and CLSSLK appeared in three unrelated proteins—6-phosphogluconate dehydrogenase, the transcription factor HIVEP2, and the E3 ligase Nedd4—complicating protein assignment. These unspecific peptides were excluded from network visualizations using the neXtProt Peptide Uniqueness Checker to maintain data accuracy. Furthermore, while AlphaFold predictions and static crystal structures provided useful reference points, they failed to account for redox-dependent conformational changes, underscoring the need for dynamic structural analyses under oxidative conditions.

To translate these findings into biological or clinical insights, future work should investigate the functional consequences of individual disulfide bonds in vivo. Time-resolved redox proteomics, and structural dynamics under oxidative stress will help elucidate the regulatory logic of disulfide network formation. Furthermore, integrating these redox-specific networks with transcriptomic or metabolomic data may reveal how oxidative stress reprograms cellular systems at multiple levels.

Collectively, this study advances redox proteomics by comprehensively identifying redox-sensitive proteins and disulfide-linked interactions under non-reducing conditions. Our findings reveal that disulfide bonds are selective, biologically meaningful modifications that organize signaling, metabolic, and structural networks—particularly in isoform-specific contexts. These results offer a foundation for further exploration of Cys-dependent regulation and redox-controlled protein networks in health and disease.

### Experimental Design

This study aimed to elucidate redox-dependent disulfide bond formation and associated metabolic responses in MDA-MB-231 breast cancer cells exposed to oxidative stress. Cells were treated with H₂O₂ at five concentrations (0, 0.1, 0.25, 0.5 and 1 mM) for 1 hour. Each condition was performed in triplicate using independent biological replicates. We applied an integrated approach combining LC-MS-based metabolomics and tandem mass tag (TMT)-based non-reducing proteomics. Disulfide bonds were preserved through non-reducing SDS-PAGE and identified using the DBond algorithm. Changes in metabolic pathway activity and redox-sensitive protein isoforms were quantitatively assessed.

### Cell culture

MDA-MB-231 cells were cultured in MEM alpha (CM054-050) from GenDEPOT, supplemented with 10% fetal bovine serum and penicillin/streptomycin (Sigma, USA) at 37℃ with 5% CO_2_.

### Sample preparation for metabolomics

For metabolomics analysis, we utilized the two-step method as previously described(*48*). MDA-MB-231 cells were treated with 0, 0.1, 0.25, 0.5, and 1 mM H_2_O_2_ in Hank’s Balanced Salt Solution (HBSS) for 1 h (n = 5 per concentration). Following treatment, cells were washed with phosphate-buffered saline (PBS), then their metabolites were extracted using 1 mL of cold 60% methanol and collected by scraping. For normalization, DNA concentrations were measured using a Nanodrop. After adding 400 μL of chloroform to each sample, cells were vortexed twice for 10 s each and centrifuged at 4,000 rpm for 10 min at 4℃. The supernatant from the upper hydrophilic layer was transferred to an Amicon Ultra 3K 0.5 mL centrifugal filter (Merck, USA) and centrifuged at 13,000 × g for 1 h at 4℃ to remove macromolecules including proteins. The filtrate was then concentrated using a vacuum concentrator and analyzed using Mass spectrometry.

### Redox-sensitive metabolite profiling using MS spectrometry

MS spectrometry was performed using an Acquity UPLC I-class system coupled with Q-Tof mass spectrometers (Xevo G2-XS, Waters Co., UK) for intracellular metabolites. Two different columns and mobile phases were used depending on the ionization mode.

For negative ion mode, intracellular metabolites were analyzed using an Acquity UPLC HSS T3 column (2.1 × 150 mm, 1.8 µm, Waters Co., UK). The mobile phases consisted of 0.01 M tributylamine/0.02 M acetic acid in distilled water (A) and methanol (B). The total flow rate was 0.3 mL/min. The LC gradient program started with 2% B, held for 5 min, then increased to 80% B at 30 min, held for 2 min, and returned to 2% B at 32.1 min. The column was conditioned for another 3.9 min, resulting in a total run time of 36 minutes. The injection volume was 5 µL, and the oven temperature was set to 40℃. Mass spectral data were acquired in negative electrospray ionization mode with a capillary voltage of 1.2 kV, cone voltage of 40 V, source temperature of 120℃, and cone gas and desolvation gas flow rates of 40 and 800 L/h, respectively.

For positive ion mode, intracellular metabolites were analyzed using an Intrada Amino Acid column (2.0 × 150 mm, 3 µm, Imtakt Co., Japan). The mobile phases consisted of 0.3% formic acid in acetonitrile (A) and 0.08 M ammonium formate in 20% acetonitrile (B). The total flow rate was 0.35 mL/min. The LC gradient program started with 20% B, held for 5 min, then increased to 100% B at 14 min, held for 2 min, and returned to 20% B at 16.1 min. The column was conditioned for another 3.9 min, resulting in a total run time of 20 min. The injection volume was 5 µL, and the oven temperature was set to 35℃. Mass spectral data were acquired in positive electrospray ionization mode with a capillary voltage of 2.5 kV, cone voltage of 30 V, source temperature of 100℃, and cone gas and desolvation gas flow rates of 20 and 400 L/h, respectively.

### Sample preparation for proteomics

For proteomics analysis, MDA-MB-231 cells were treated with 0, 0.1, 0.25, 0.5, and 1 mM H_2_O_2_ in HBSS buffer for 1 h. The cells were then washed with PBS and lysed in RIPA buffer (50 mM Tris, 150 mM NaCl, 0.1% SDS, 0.5% deoxycholate, 1% NP 40, pH 8.0) supplemented with protease inhibitor (Roche) and 1 mM sodium orthovanadate. The lysates were incubated on ice for 15 min with vortexing every 5 min, then centrifuged at 14,000 rpm for 15 min at 4℃. The protein concentration in the supernatant was measured using the BCA assay (Pierce, USA). Each lysate was prepared with non-reducing SDS-PAGE sample buffer and boiled at 95℃ for 5 min.

### Silver staining

To maintain protein oxidation state, the lysates were prepared using a non-reducing SDS-PAGE sample buffer. The samples were loaded (70 μg of protein) onto a PROTEAN II xi 2-D Cell system (BIO-RAD, USA) for separation and further analysis. The entire gel was silver stained based on the instructions on the PlusOne Silver Staining Kit (GE Healthcare, USA) and divided into 18 bands.

### Redox-sensitive proteomics using in-gel digestion, protein quantification, and modification analysis with TMT labeling and mass spectrometry

Gel bands were excised with a scalpel into 18 fractions based on molecular weight, destained with 15 mM potassium ferricyanide and 50 mM sodium thiosulfate. After destaining, the gel bands were washed with distilled water (DW) to completely remove the destaining reagent. The pH was adjusted to 8.0 using 200 mM triethylammonium bicarbonate buffer (TEAB) to facilitate trypsin digestion. The gels were dehydrated with acetonitrile (ACN), rehydrated with 10–20 μL of 25 mM TEAB containing 20 ng/μL sequencing-grade trypsin (Promega, Madison, WI, USA), and incubated at 37°C for 15–17 h. Peptides were extracted using 30 μL of 60% ACN solution. The extracts were pooled and dried using a vacuum concentrator for MS analysis. Acid was omitted during TMT labeling to maintain a final peptide extract pH of ∼8.5, optimizing labeling efficiency.

Each fraction from MDA-MB-231 was resuspended in 30 μL of TMT Dissolution Buffer (Thermo Fisher Scientific, Waltham, MA, USA) and aminolabeled with one of the six TMT tags at 25°C for 1 h, and subsequently pooled in equal amounts.

TMT-labeled peptides were concentrated using a vacuum concentrator and dissolved in 10% ACN containing 0.1% formic acid. The dissolved peptides were desalted on-line using a trap column (180 μm x 20 mm, Symmetry® C18, Waters Co., UK) prior to separation. Peptide separation was performed over 80 min on a C18 reversed-phase analytical column (75 μm x 250 mm, 1.7 μm particle size, BEH130 C18, Waters Co. UK) equipped with an integrated electrospray ionization PicoTip^TM^ (± 10 μm, New Objective, USA). The nanoAcquity^TM^ UPLC system coupled with ESI-MS (nanoLC-MS, SYNAPT^TM^ G2Si HDMS^TM^, Waters Co., UK) was used for analysis as described below(*13*).

The mobile phases consisted of 0.1% formic acid in DW (Solvent A) and 0.1% formic acid in ACN (Solvent B). The optimized LC gradient program began with 95% Solvent A and 5% Solvent B, held for 0.5 min, followed by a gradual increase to 40% Solvent B at 40 min and 60% Solvent B at 50 min. The gradient then reached 80% Solvent B at 60 min, held for 5 min, and returned to the initial condition of 95% Solvent A and 5% Solvent B at 70 min. The column was conditioned for an additional 10 min to ensure stability. The flow rate was maintained at 0.35 mL/min throughout the run, and the injection volume was set to 5 μL.

Mass spectrometry was performed under optimized conditions to ensure accurate and reproducible results. The instrument was operated in positive electrospray ionization mode (ES+), with a capillary voltage of 2.5 kV and a source temperature of 100°C. The sampling cone was set to 40 eV, and the source offset was adjusted to 80 eV. The source gas flow was maintained at 0 mL/min, while the desolvation temperature was set to 250°C. Cone gas flow and desolvation gas flow rates were set to 50 L/h and 600 L/h, respectively. Nanoflow gas pressure was maintained at 0.5 bar, purge gas flow at 600 mL/h, and nebulizer gas flow at 6.5 bar.

The raw data files obtained from the MS were converted to mascot generic files using Mascot Daemon (version 2.8.0). Tandem MS (MS/MS) spectra were matched to amino acid sequences in the SwissProt human database (version 57.15, 20,233 entries). The search parameters included a 0.3 Da tolerance for peptide and fragment ions, digestion with trypsin allowing up to one missed cleavage, and variable modifications such as acetylation (K), dehydroalanine (C), deamidation (NQ), glygly (K), oxidation (M), phosphorylation (ST and Y), propionamide (C), and trioxidation (C).

After identifying potential PTMs with the Mascot search, additional searches were performed using DBond (http://prix.hanyang.ac.kr/), a specialized algorithm for detecting disulfide-linked peptides in MS/MS spectra, against FASTA files of each protein, downloaded from NCBI (http://www.ncbi.nlm.nih.gov/protein/). Only significant hits, indicated by MASCOT probability analysis (probability based Mowse score p < 0.05), were considered. All assignments were verified through automatic and manual spectral interpretation.

### Puromycin assay

MDA-MB-231 cells were seeded in 60 mm cell culture dishes at a density of 6.0 x 10^5^ cells. After 24 h, cells were treated with H_2_O_2_ (0.1, 0.25, 0.5, or 1 mM) in PBS for 1 h. The buffer was then replaced with a fresh medium containing puromycin, and cells were incubated for 0.5, 2, 6, or 12 h to label nascent peptides. The cells were then incubated with 1 μg/mL of puromycin in the medium for 1 h before harvest. The samples were lysed, and puromycin-incorporated nascent peptides were detected using a puromycin antibody (Millipore, USA).

### Endogenous immunoprecipitation in non-reducing condition

MDA-MB-231 cells (2.2 x 10^6^ cells) were seeded in 100 mm cell culture dishes and harvested 24 h later. Subsequently, the cells were lysed using RIPA buffer, and the lysate concentration was determined via BCA assay. 500 μg of lysates were used for endogenous immunoprecipitation. The lysates were incubated overnight at 4℃ with 10 μg of normal rabbit IgG (Cell signaling 2729) as a control or γ-actin rabbit monoclonal antibody (Abcam, ab200046) to detect endogenous actin. Protein G Sepharose 4 Fast Flow (Cytiva, 17-0618-01) 40 μl were added to each solution and incubated for an additional 3 hours at 4℃. The lysates were then centrifuged to precipitate beads, washed 3 times, and boiled for 5 min at 95℃ in a non-reducing sample buffer. The samples were separated and silver stained, as mentioned above.

## Supporting information

Disulfide bond raw data

MS, PTM raw data

## Statistical Analysis

All quantitative values were normalized to the 0 mM H₂O₂ condition. Statistical significance was assessed using one-way ANOVA followed by Tukey’s post hoc test to compare differences across treatment groups. For comparisons of individual treatment conditions against the control (0 mM), two-tailed unpaired Student’s t-tests were also used. A p-value < 0.05 was considered statistically significant.

## Data analysis

Pathway analysis, enrichment analysis, and principal component analysis (PCA) were performed using MetaboAnalyst 6.0 (https://www.metaboanalyst.ca/) on metabolites exhibiting over 20% variation after treatment with 1 mM H_2_O_2_ compared to that at 0 mM H_2_O_2_. Metabolite graphs were generated using GraphPad Prism 9. Protein–protein interaction networks and protein classifications were analyzed using STRING (STRING version 12.0, https://string-db.org/) and PANTHER (https://www.pantherdb.org/). Graphic comparisons of sensitive and insensitive protein subgroups according to PANTHER classification were performed using the graphing functions of SRplot (https://www.bioinformatics.com.cn/srplot). Intra-disulfide or Inter-disulfide bonds were identified using the DBond program from PRIX (https://prix.hanyang.ac.kr/download/dbond.jsp) and visualized using Cytoscape(*19*).

## Author contributions

Conceptualization: YL, KJL, EJS

Methodology: KJL, SN

Investigation: YL, TKK, JJ

Visualization: YL

Supervision: KJL, EJS

Writing—original draft: YL

Writing—review & editing: YL, KJL, JJ, SN, EJS

## Data and materials availability

All data are available in the main text or the supplementary materials.

