## Supplementary Figure for "Deciphering the oxidative modifications via disulfide mapping"

### Slide 1
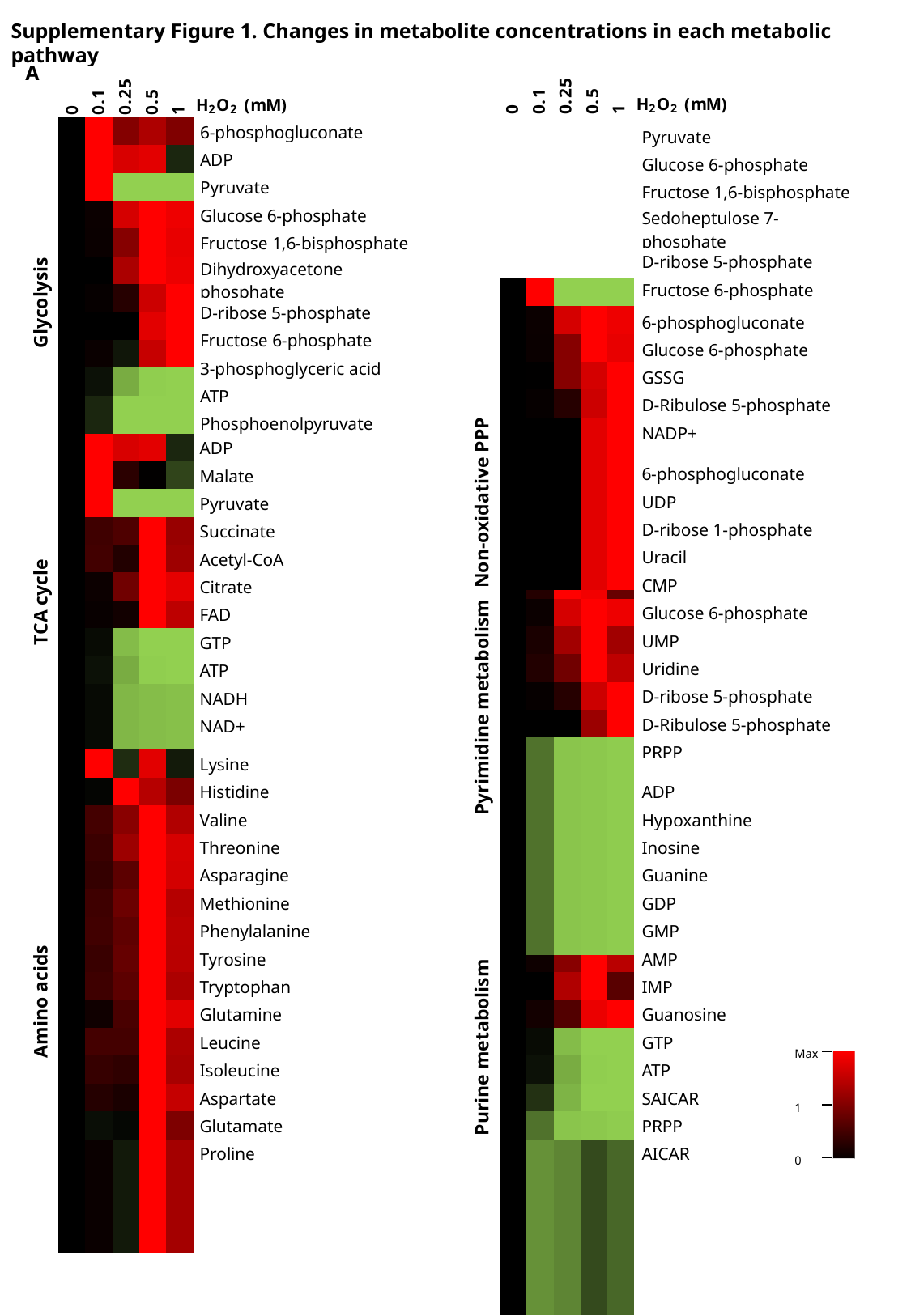

Supplementary Figure 1. Changes in metabolite concentrations in each metabolic pathway
A
| Non-oxidative PPP |
| --- |
| Glycolysis |
| --- |
| 6-phosphogluconate |
| --- |
| ADP |
| Pyruvate |
| Glucose 6-phosphate |
| Fructose 1,6-bisphosphate |
| Dihydroxyacetone phosphate |
| D-ribose 5-phosphate |
| Fructose 6-phosphate |
| 3-phosphoglyceric acid |
| ATP |
| Phosphoenolpyruvate |
| Pyruvate |
| --- |
| Glucose 6-phosphate |
| Fructose 1,6-bisphosphate |
| Sedoheptulose 7-phosphate |
| D-ribose 5-phosphate |
| Fructose 6-phosphate |
| Oxidative PPP |
| --- |
| 6-phosphogluconate |
| --- |
| Glucose 6-phosphate |
| GSSG |
| D-Ribulose 5-phosphate |
| NADP+ |
| TCA cycle |
| --- |
| ADP |
| --- |
| Malate |
| Pyruvate |
| Succinate |
| Acetyl-CoA |
| Citrate |
| FAD |
| GTP |
| ATP |
| NADH |
| NAD+ |
| Pyrimidine metabolism |
| --- |
| 6-phosphogluconate |
| --- |
| UDP |
| D-ribose 1-phosphate |
| Uracil |
| CMP |
| Glucose 6-phosphate |
| UMP |
| Uridine |
| D-ribose 5-phosphate |
| D-Ribulose 5-phosphate |
| PRPP |
| Amino acids |
| --- |
| Lysine |
| --- |
| Histidine |
| Valine |
| Threonine |
| Asparagine |
| Methionine |
| Phenylalanine |
| Tyrosine |
| Tryptophan |
| Glutamine |
| Leucine |
| Isoleucine |
| Aspartate |
| Glutamate |
| Proline |
| Purine metabolism |
| --- |
| ADP |
| --- |
| Hypoxanthine |
| Inosine |
| Guanine |
| GDP |
| GMP |
| AMP |
| IMP |
| Guanosine |
| GTP |
| ATP |
| SAICAR |
| PRPP |
| AICAR |
Max
1
0

### Slide 2
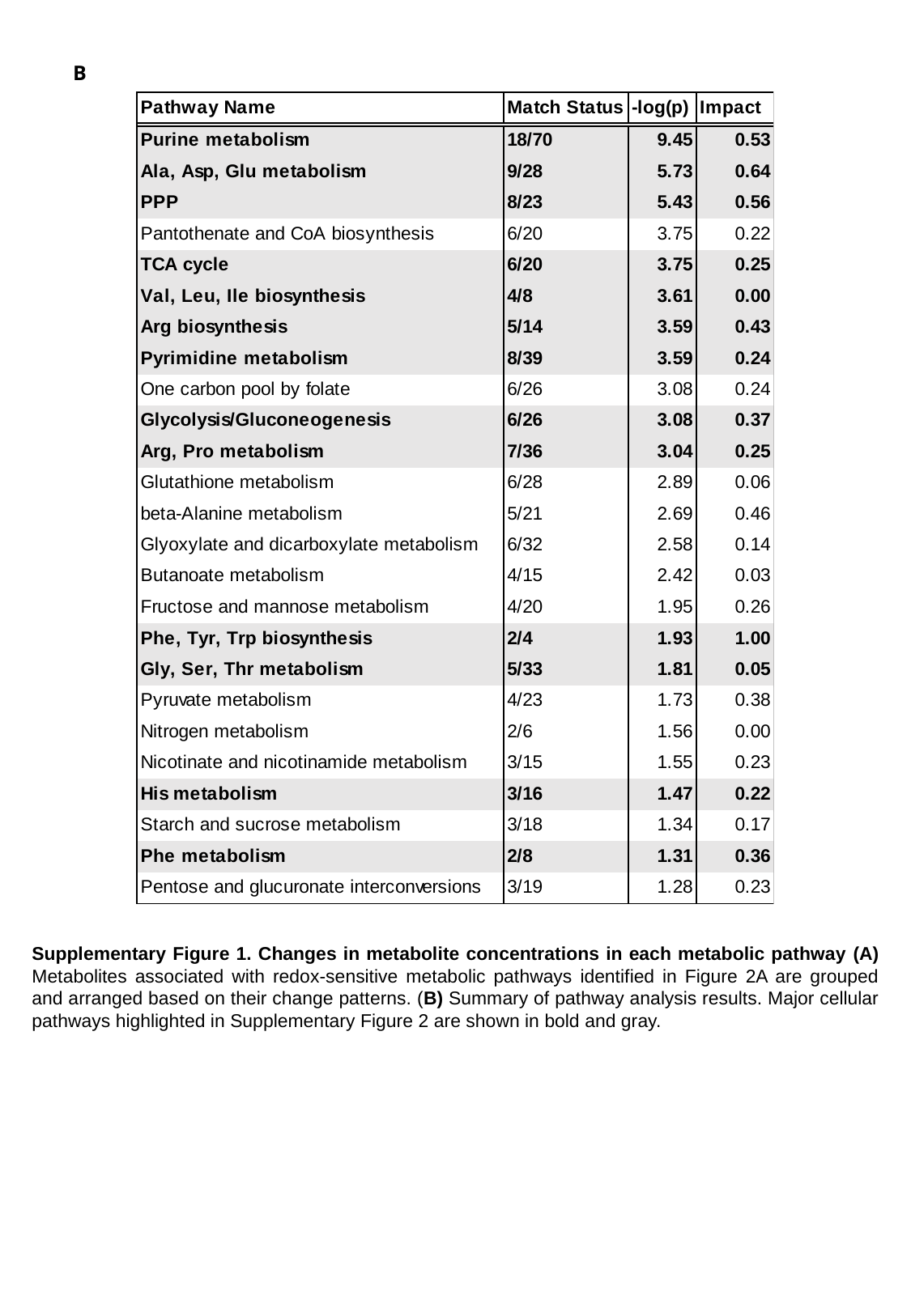

B
Supplementary Figure 1. Changes in metabolite concentrations in each metabolic pathway (A) Metabolites associated with redox-sensitive metabolic pathways identified in Figure 2A are grouped and arranged based on their change patterns. (B) Summary of pathway analysis results. Major cellular pathways highlighted in Supplementary Figure 2 are shown in bold and gray.

### Slide 3
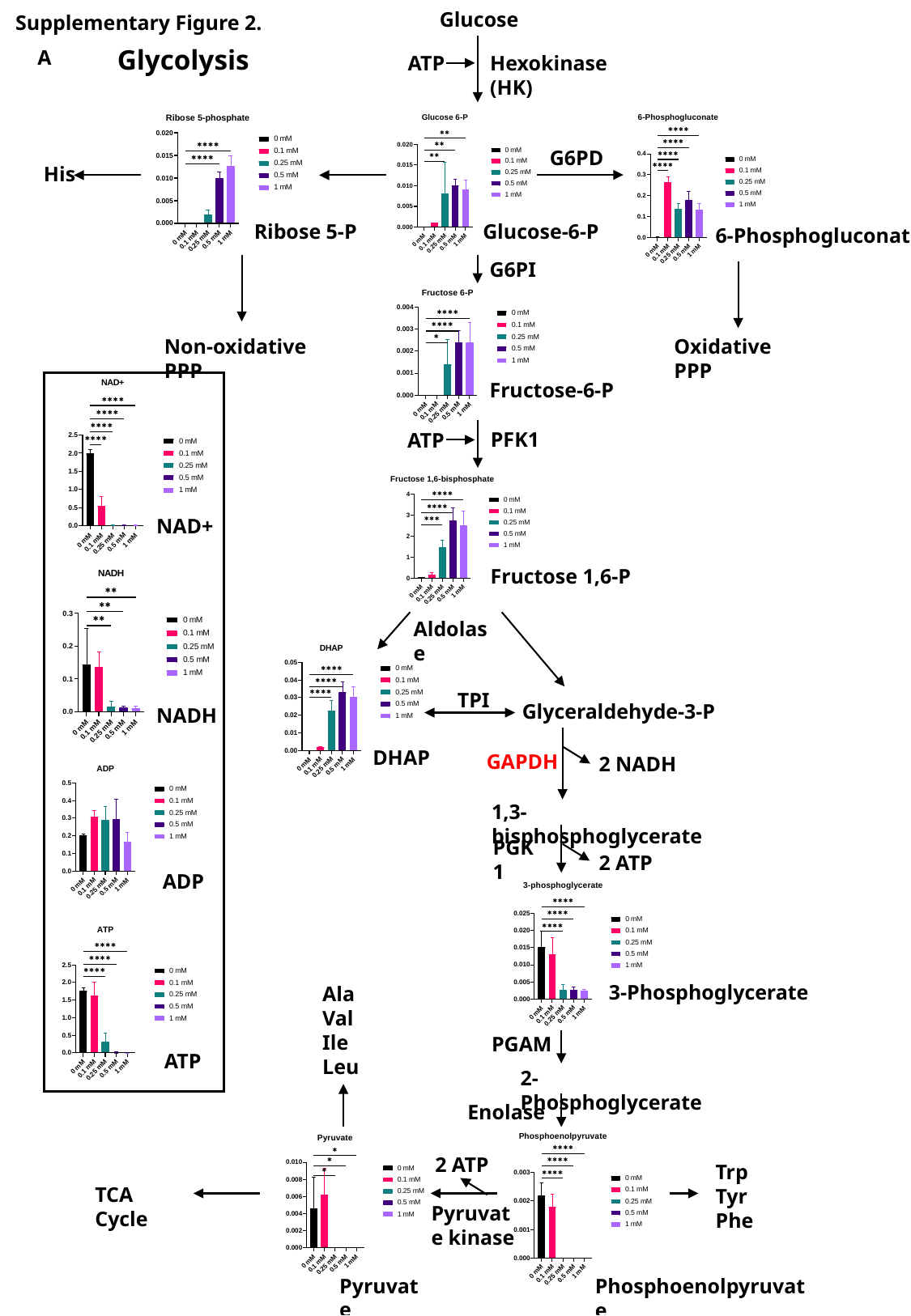

Glucose
Supplementary Figure 2.
Glycolysis
A
ATP
Hexokinase (HK)
G6PD
His
Ribose 5-P
Glucose-6-P
6-Phosphogluconate
G6PI
Oxidative PPP
Non-oxidative PPP
Fructose-6-P
PFK1
ATP
NAD+
Fructose 1,6-P
Aldolase
TPI
Glyceraldehyde-3-P
NADH
DHAP
GAPDH
2 NADH
1,3-bisphosphoglycerate
PGK1
2 ATP
ADP
3-Phosphoglycerate
Ala
Val
Ile
Leu
PGAM
ATP
2-Phosphoglycerate
Enolase
2 ATP
Trp
Tyr
Phe
TCA Cycle
Pyruvate kinase
Pyruvate
Phosphoenolpyruvate

### Slide 4
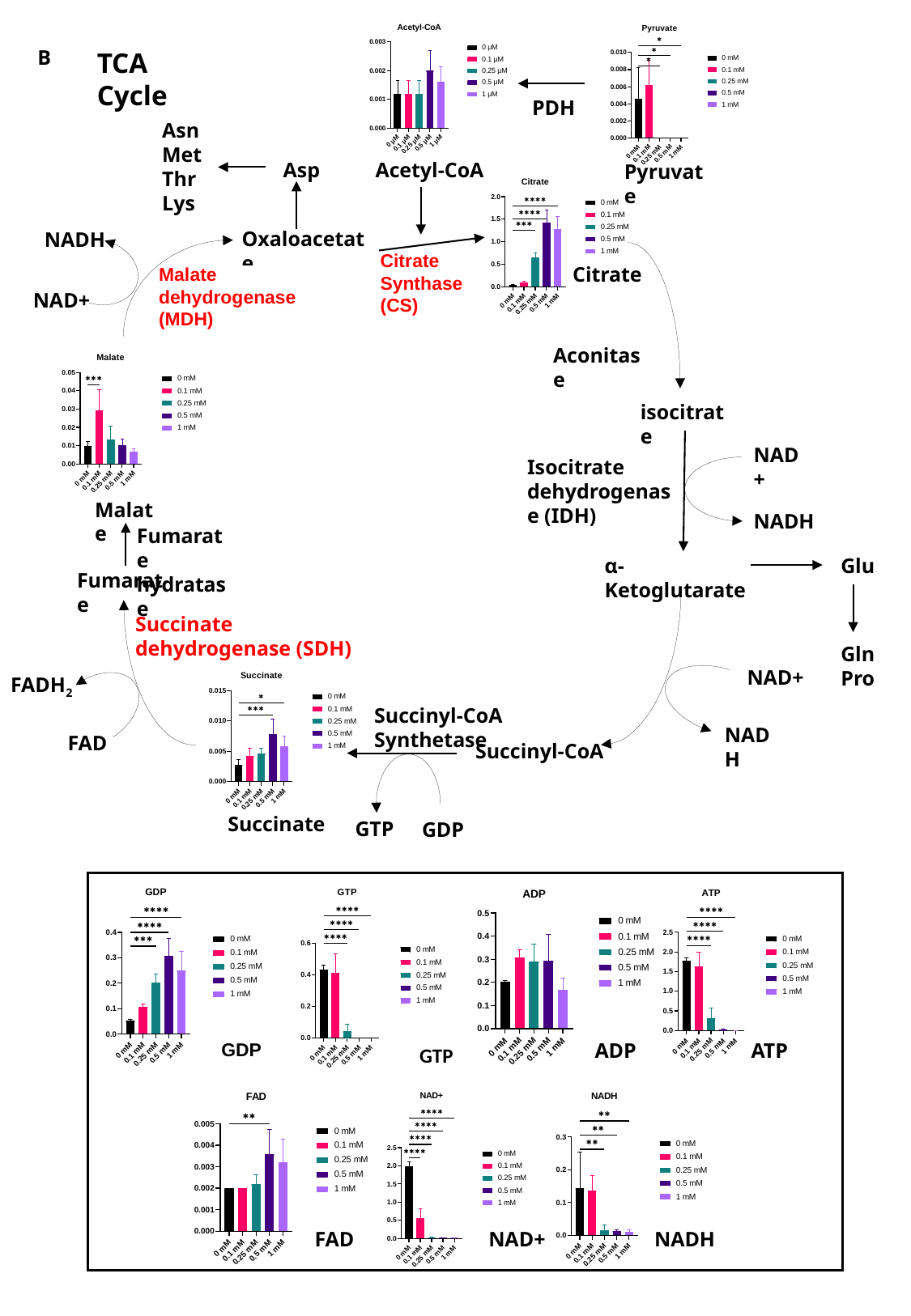

B
TCA Cycle
PDH
Asn
Met
Thr
Lys
Acetyl-CoA
Asp
Pyruvate
Oxaloacetate
NADH
Citrate Synthase (CS)
Citrate
Malate dehydrogenase (MDH)
Malate dehydrogenase (MDH)
NAD+
Aconitase
isocitrate
NAD+
Isocitrate dehydrogenase (IDH)
Malate
NADH
Fumarate hydratase
Glu
α-Ketoglutarate
Fumarate
Succinate dehydrogenase (SDH)
Gln
Pro
NAD+
FADH_2_
Succinyl-CoA Synthetase
NADH
FAD
Succinyl-CoA
Succinate
GTP
GDP
ADP
ATP
GDP
GTP
FAD
NAD+
NADH

### Slide 5
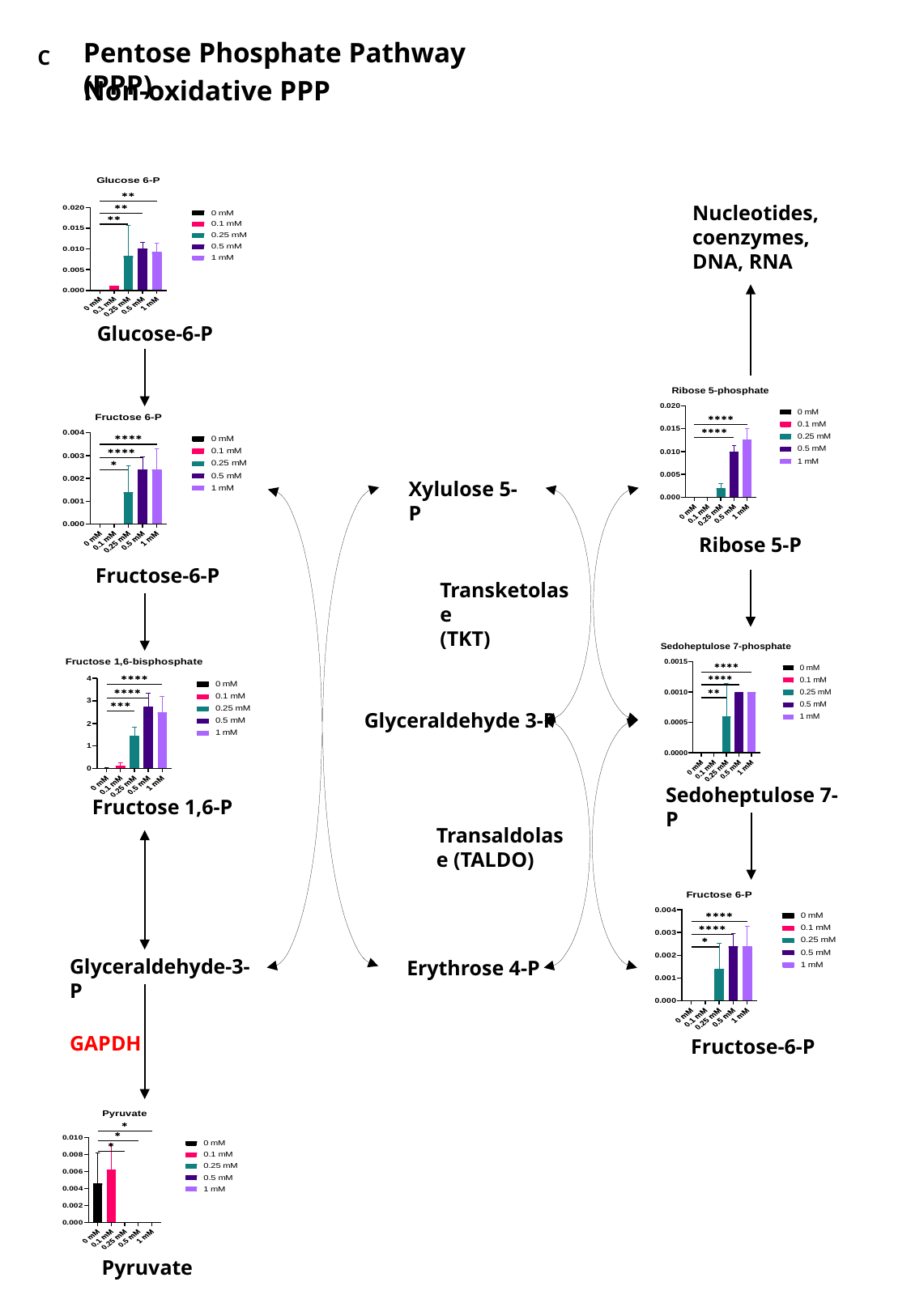

Pentose Phosphate Pathway (PPP)
C
Non-oxidative PPP
Nucleotides, coenzymes, DNA, RNA
Glucose-6-P
Xylulose 5-P
Ribose 5-P
Fructose-6-P
Transketolase
(TKT)
Glyceraldehyde 3-P
Sedoheptulose 7-P
Fructose 1,6-P
Transaldolase (TALDO)
Glyceraldehyde-3-P
Erythrose 4-P
GAPDH
Fructose-6-P
Pyruvate

### Slide 6
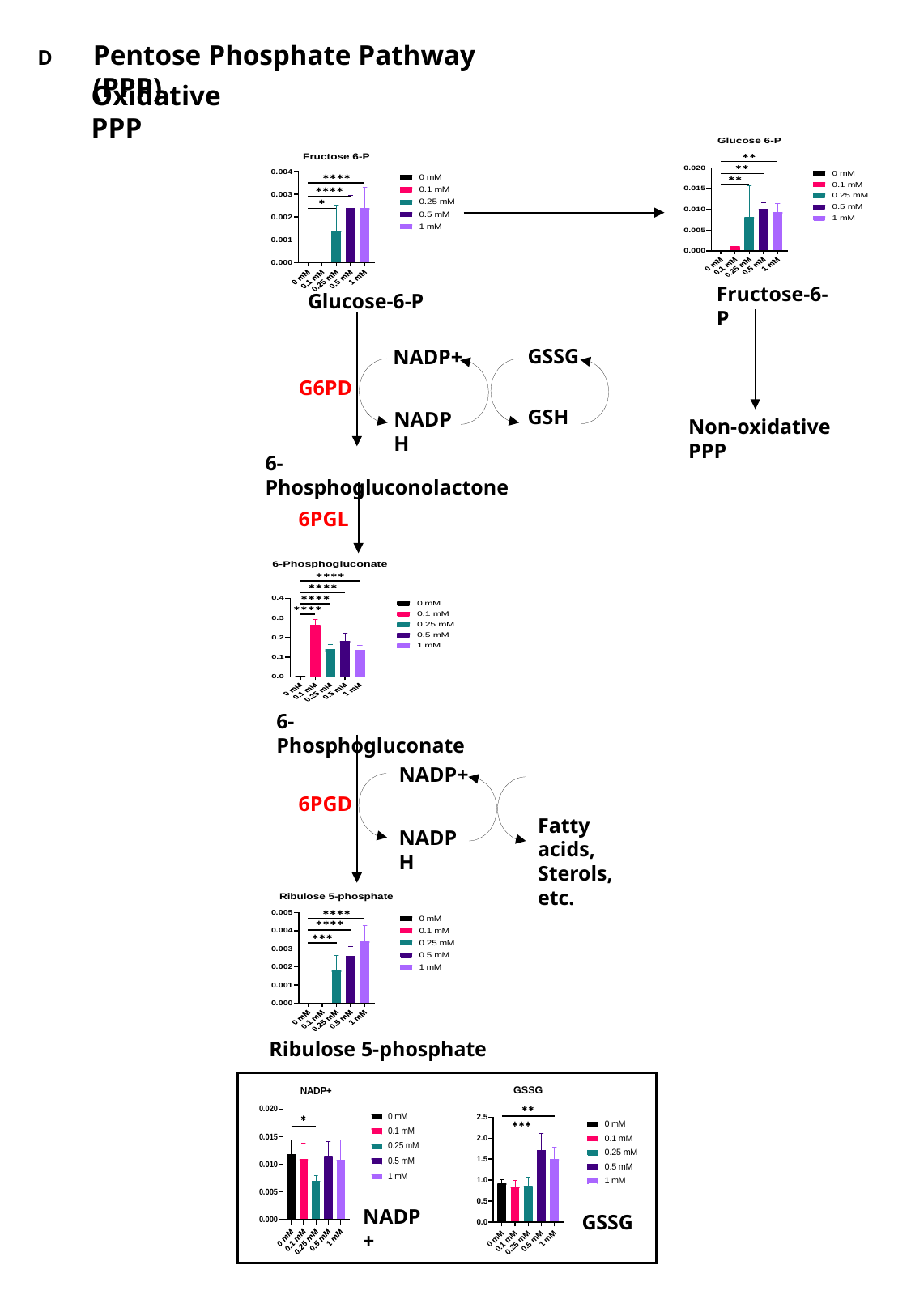

Pentose Phosphate Pathway (PPP)
D
Oxidative PPP
Fructose-6-P
Glucose-6-P
GSSG
NADP+
G6PD
GSH
NADPH
Non-oxidative PPP
6-Phosphogluconolactone
6PGL
6-Phosphogluconate
NADP+
6PGD
Fatty acids, Sterols, etc.
NADPH
Ribulose 5-phosphate
NADP+
GSSG

### Slide 7
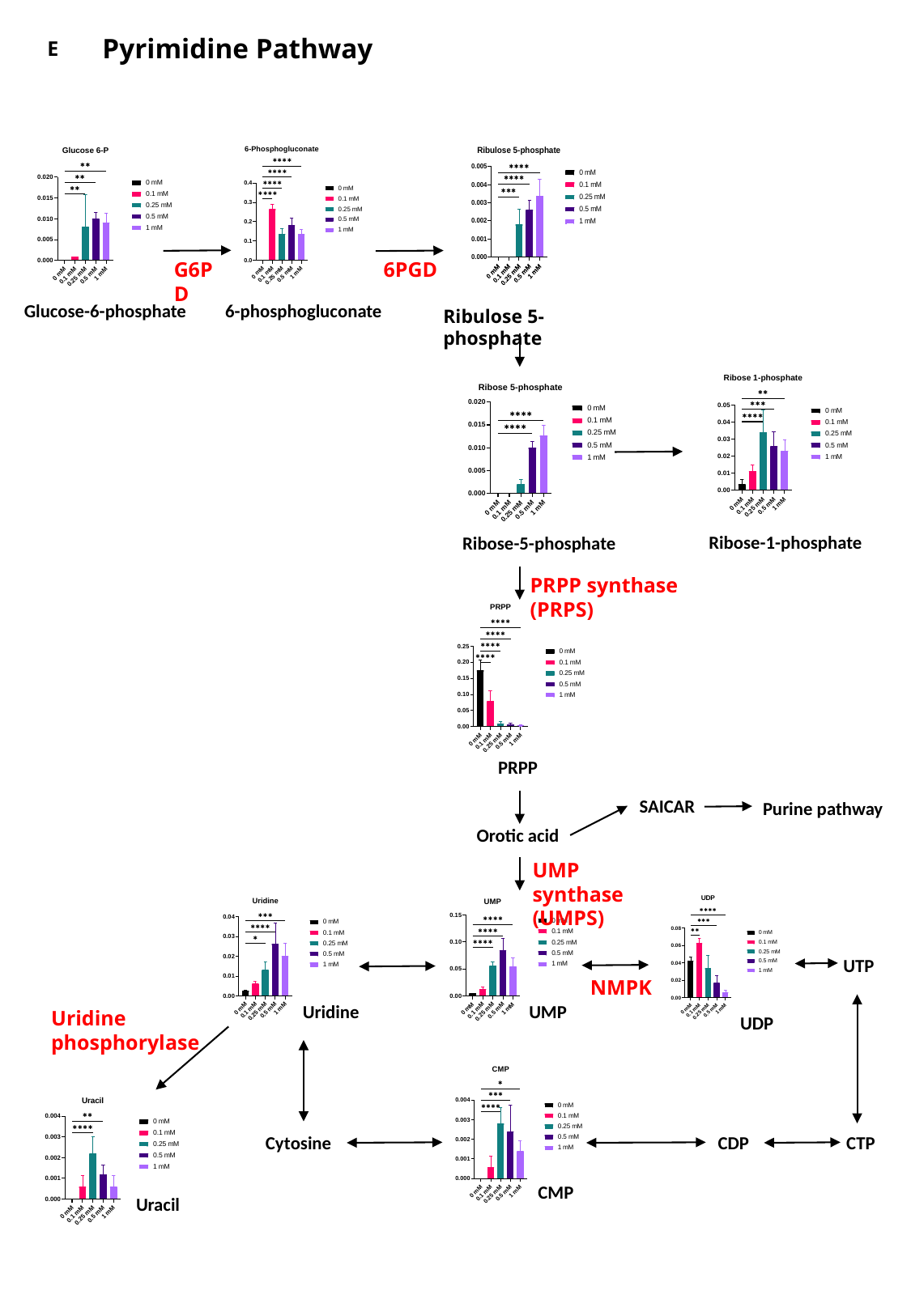

Pyrimidine Pathway
E
G6PD
6PGD
Glucose-6-phosphate
6-phosphogluconate
Ribulose 5-phosphate
Ribose-1-phosphate
Ribose-5-phosphate
PRPP synthase (PRPS)
PRPP
SAICAR
Purine pathway
Orotic acid
UMP synthase (UMPS)
UTP
NMPK
Uridine
UMP
Uridine phosphorylase
UDP
Cytosine
CDP
CTP
CMP
Uracil

### Slide 8
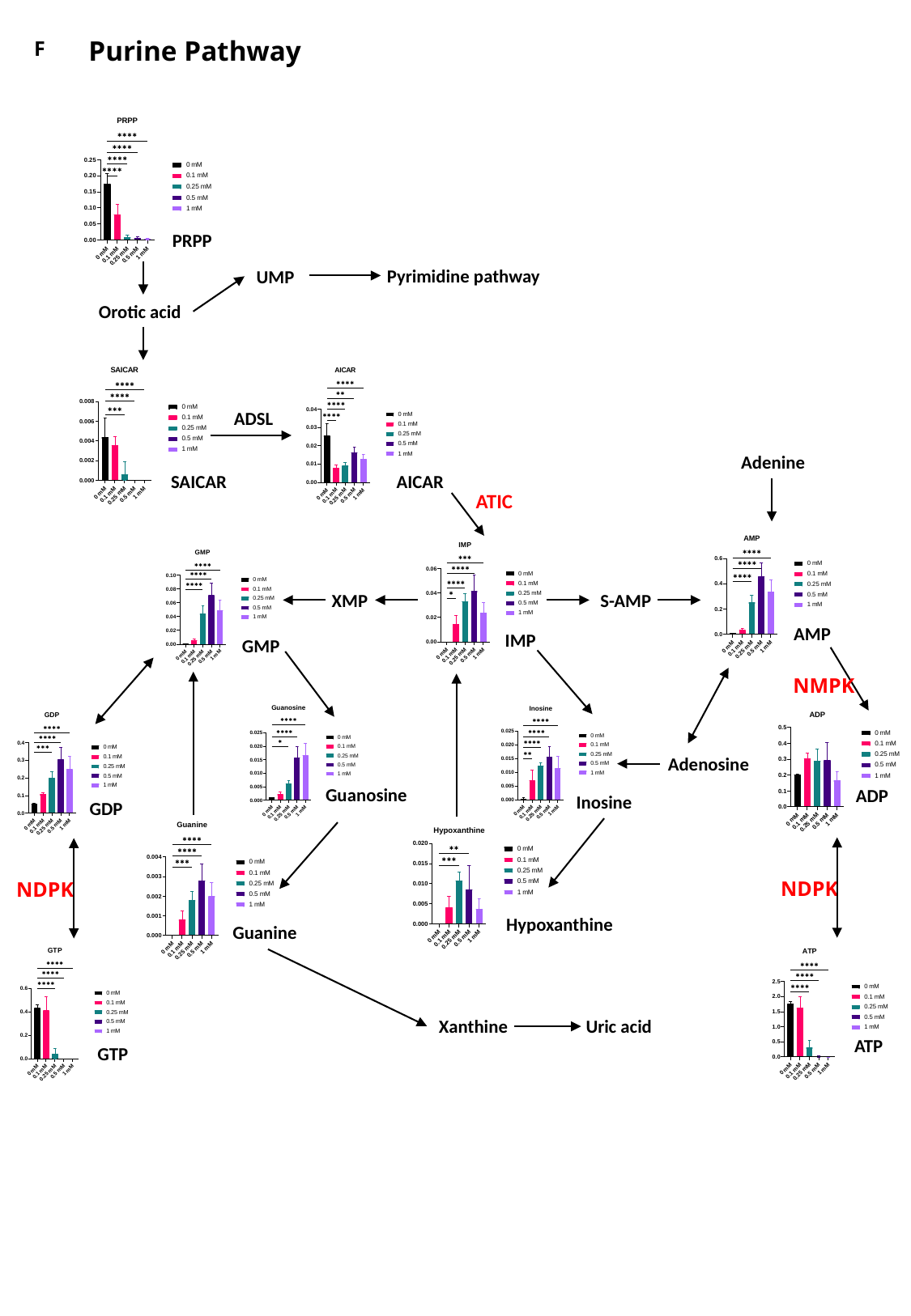

Purine Pathway
F
PRPP
Pyrimidine pathway
UMP
Orotic acid
ADSL
Adenine
AICAR
SAICAR
ATIC
XMP
S-AMP
AMP
IMP
GMP
NMPK
Adenosine
Guanosine
ADP
Inosine
GDP
NDPK
NDPK
Hypoxanthine
Guanine
Xanthine
Uric acid
ATP
GTP

### Slide 9
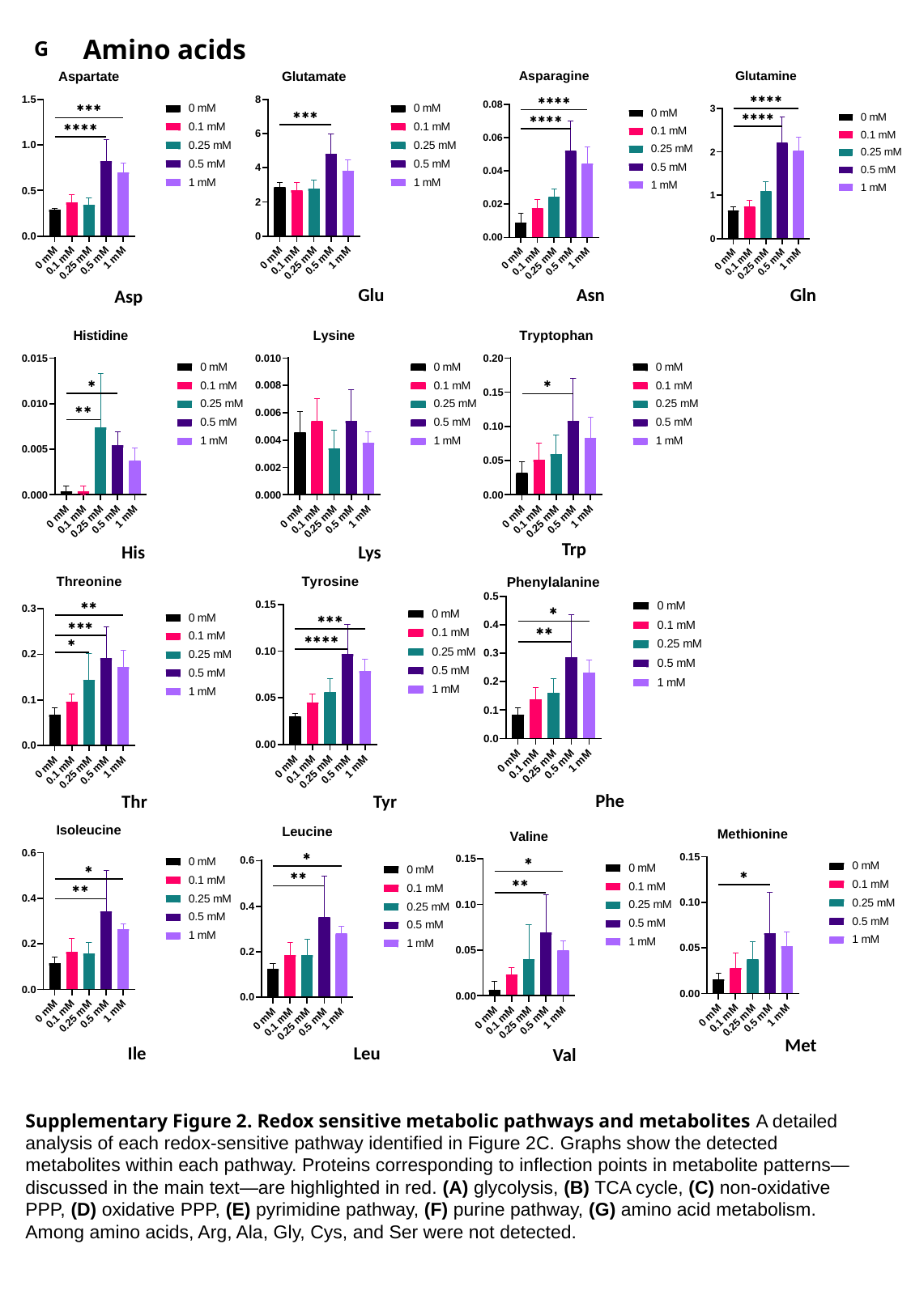

Amino acids
G
Gln
Asn
Glu
Asp
Trp
His
Lys
Phe
Tyr
Thr
Met
Ile
Leu
Val
Supplementary Figure 2. Redox sensitive metabolic pathways and metabolites A detailed analysis of each redox-sensitive pathway identified in Figure 2C. Graphs show the detected metabolites within each pathway. Proteins corresponding to inflection points in metabolite patterns—discussed in the main text—are highlighted in red. (A) glycolysis, (B) TCA cycle, (C) non-oxidative PPP, (D) oxidative PPP, (E) pyrimidine pathway, (F) purine pathway, (G) amino acid metabolism. Among amino acids, Arg, Ala, Gly, Cys, and Ser were not detected.

### Slide 10
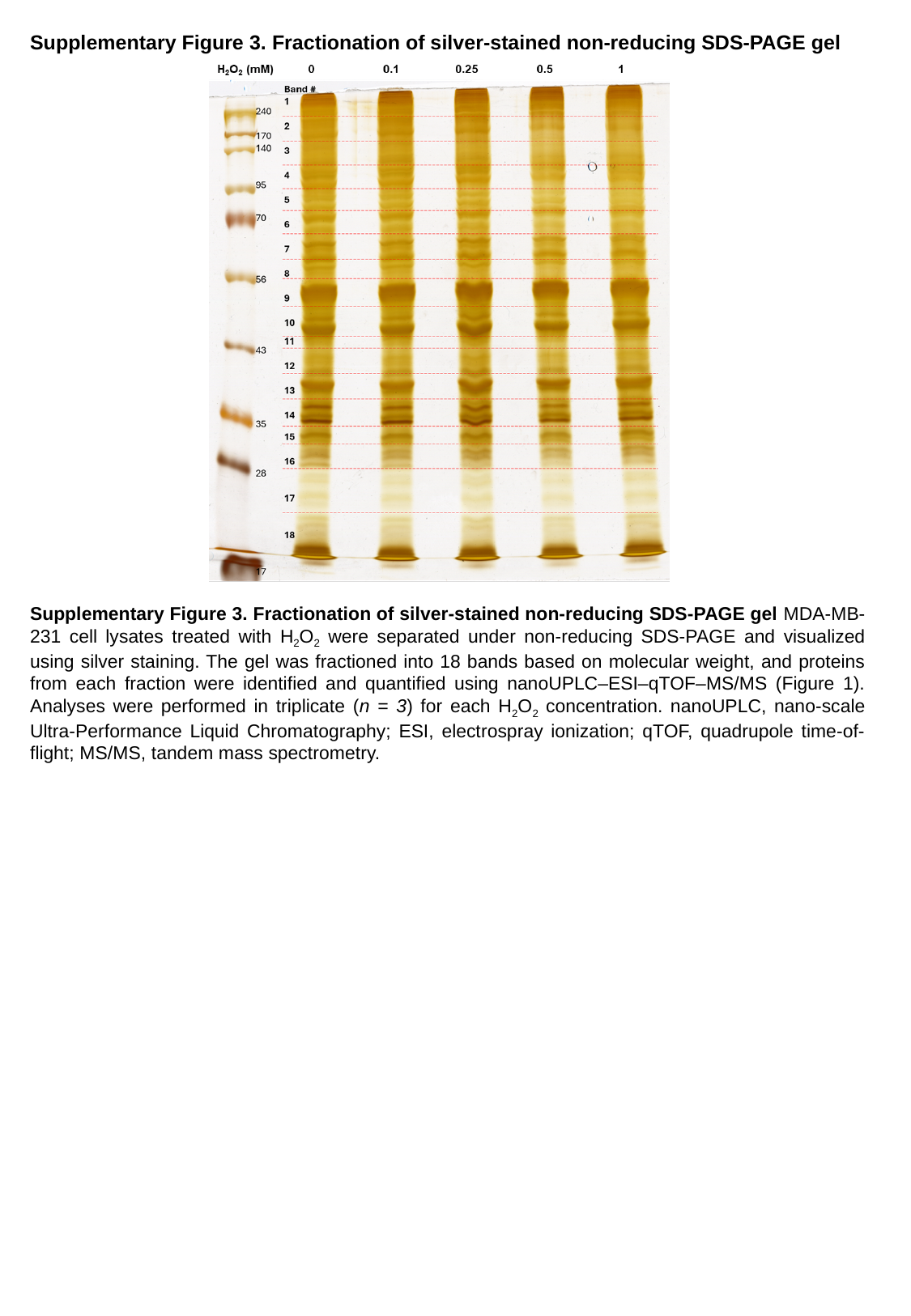

Supplementary Figure 3. Fractionation of silver-stained non-reducing SDS-PAGE gel
Supplementary Figure 3. Fractionation of silver-stained non-reducing SDS-PAGE gel MDA-MB-231 cell lysates treated with H_2_O_2_ were separated under non-reducing SDS-PAGE and visualized using silver staining. The gel was fractioned into 18 bands based on molecular weight, and proteins from each fraction were identified and quantified using nanoUPLC–ESI–qTOF–MS/MS (Figure 1). Analyses were performed in triplicate (n = 3) for each H_2_O_2_ concentration. nanoUPLC, nano-scale Ultra-Performance Liquid Chromatography; ESI, electrospray ionization; qTOF, quadrupole time-of-flight; MS/MS, tandem mass spectrometry.

### Slide 11
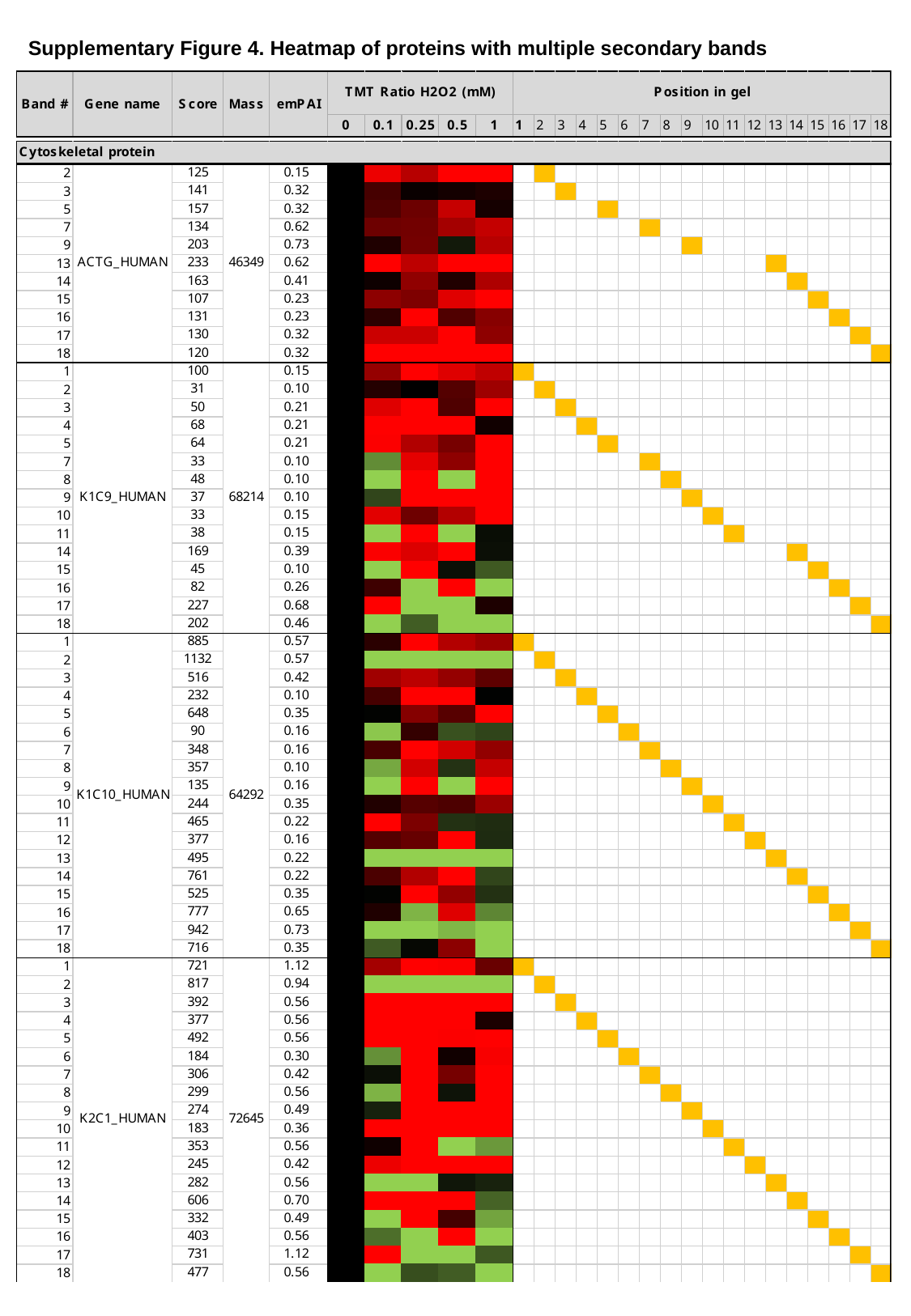

Supplementary Figure 4. Heatmap of proteins with multiple secondary bands

### Slide 12
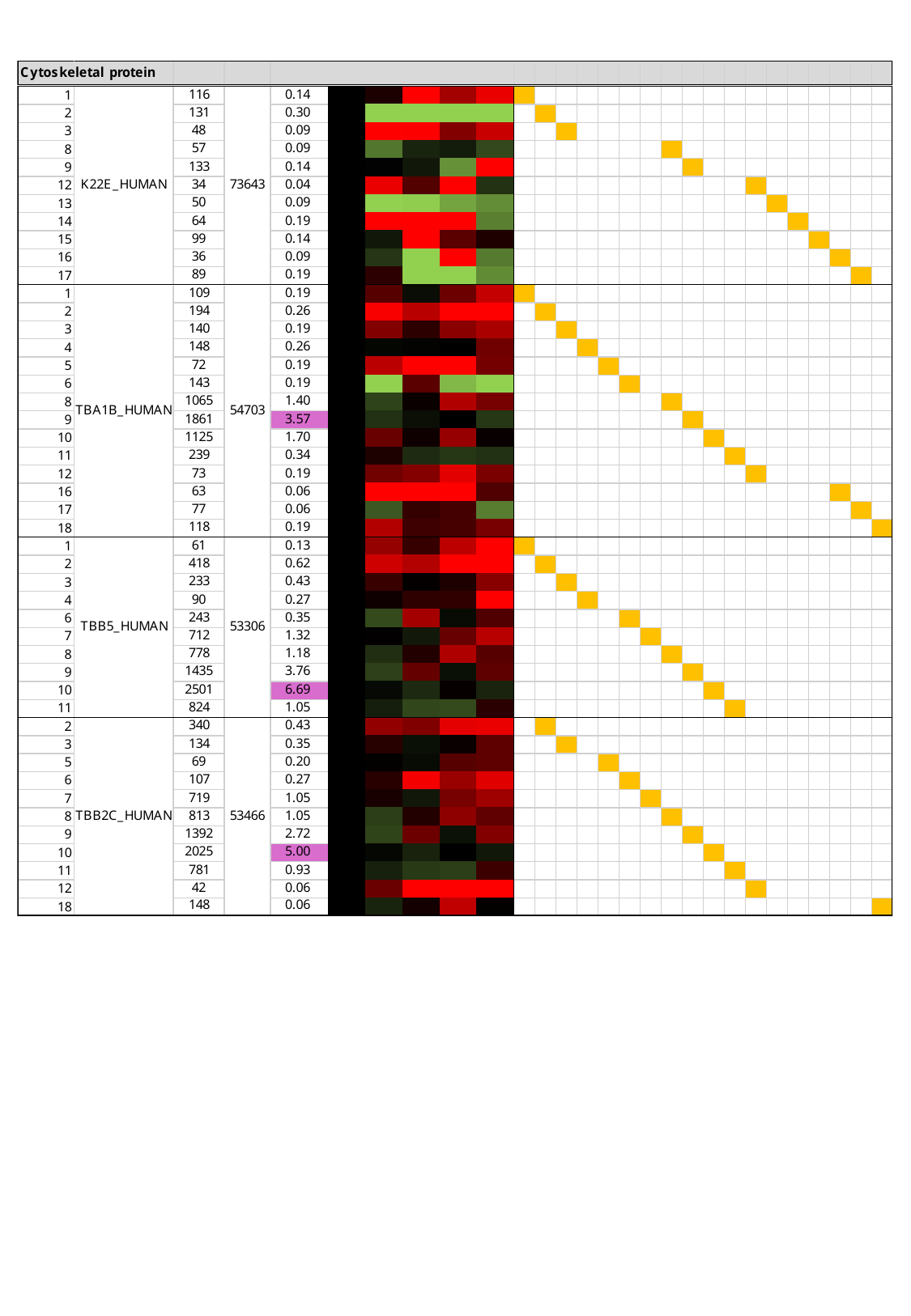

### Slide 13
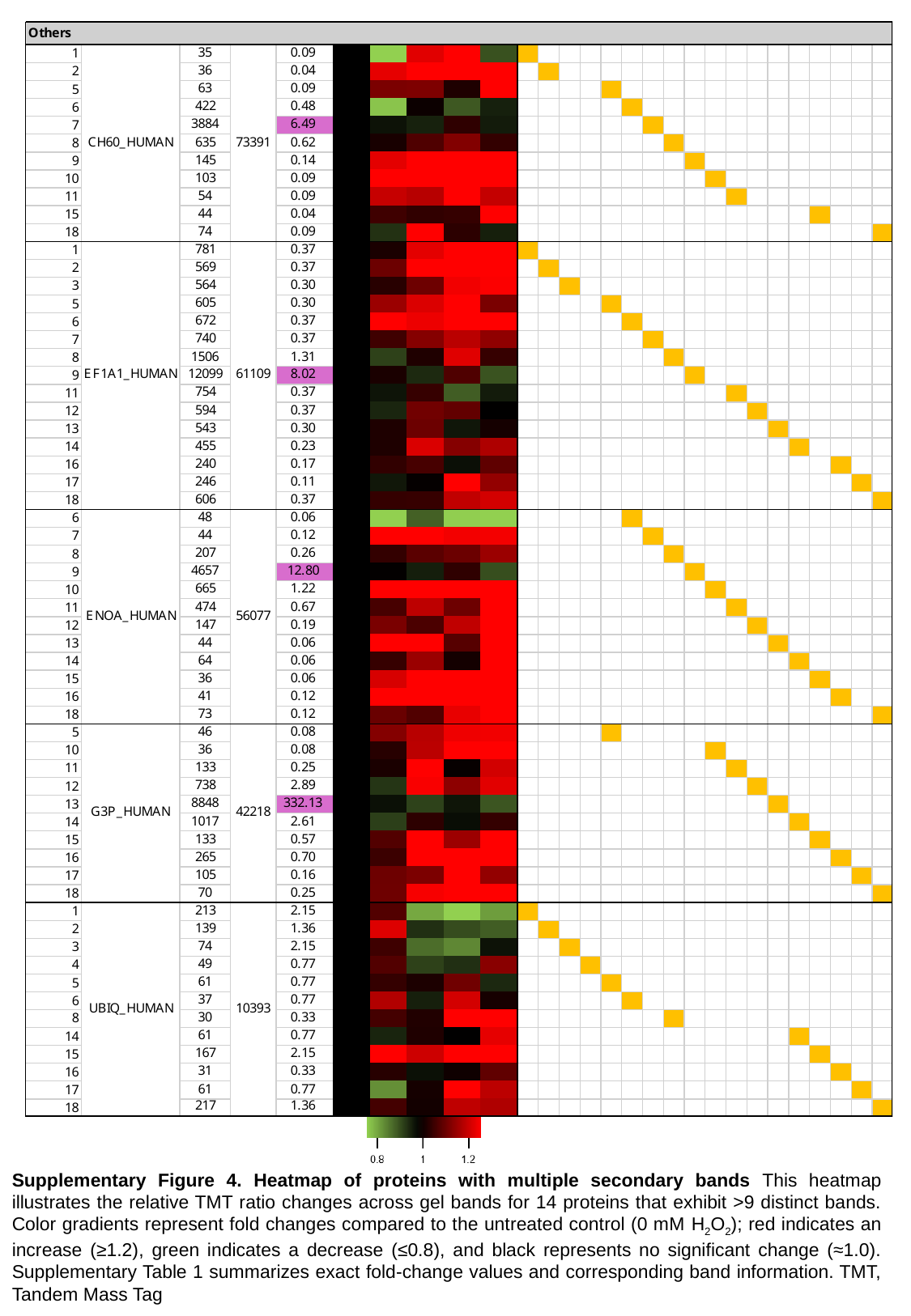

Supplementary Figure 4. Heatmap of proteins with multiple secondary bands This heatmap illustrates the relative TMT ratio changes across gel bands for 14 proteins that exhibit >9 distinct bands. Color gradients represent fold changes compared to the untreated control (0 mM H_2_O_2_); red indicates an increase (≥1.2), green indicates a decrease (≤0.8), and black represents no significant change (≈1.0). Supplementary Table 1 summarizes exact fold-change values and corresponding band information. TMT, Tandem Mass Tag

### Slide 14
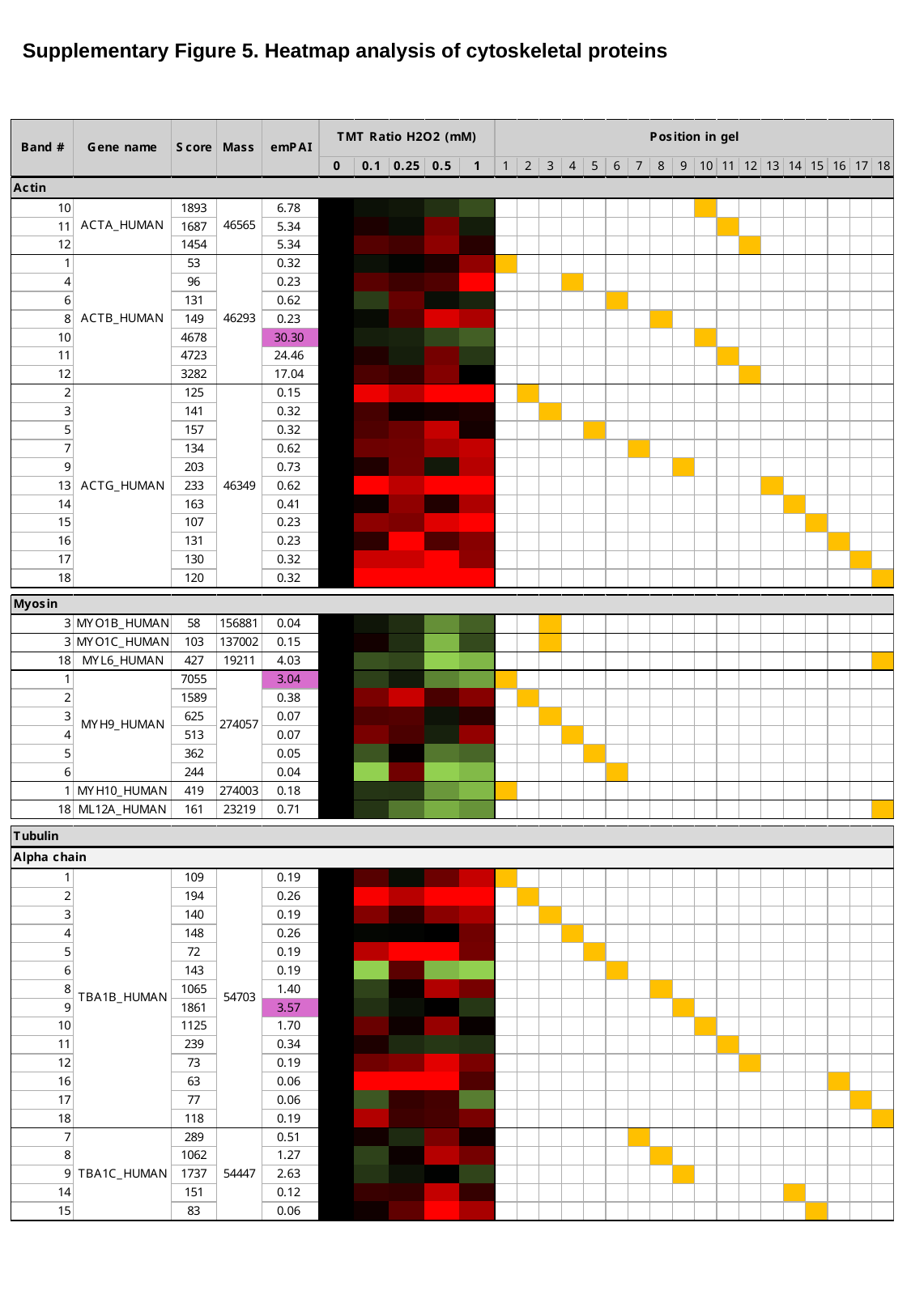

Supplementary Figure 5. Heatmap analysis of cytoskeletal proteins

### Slide 15
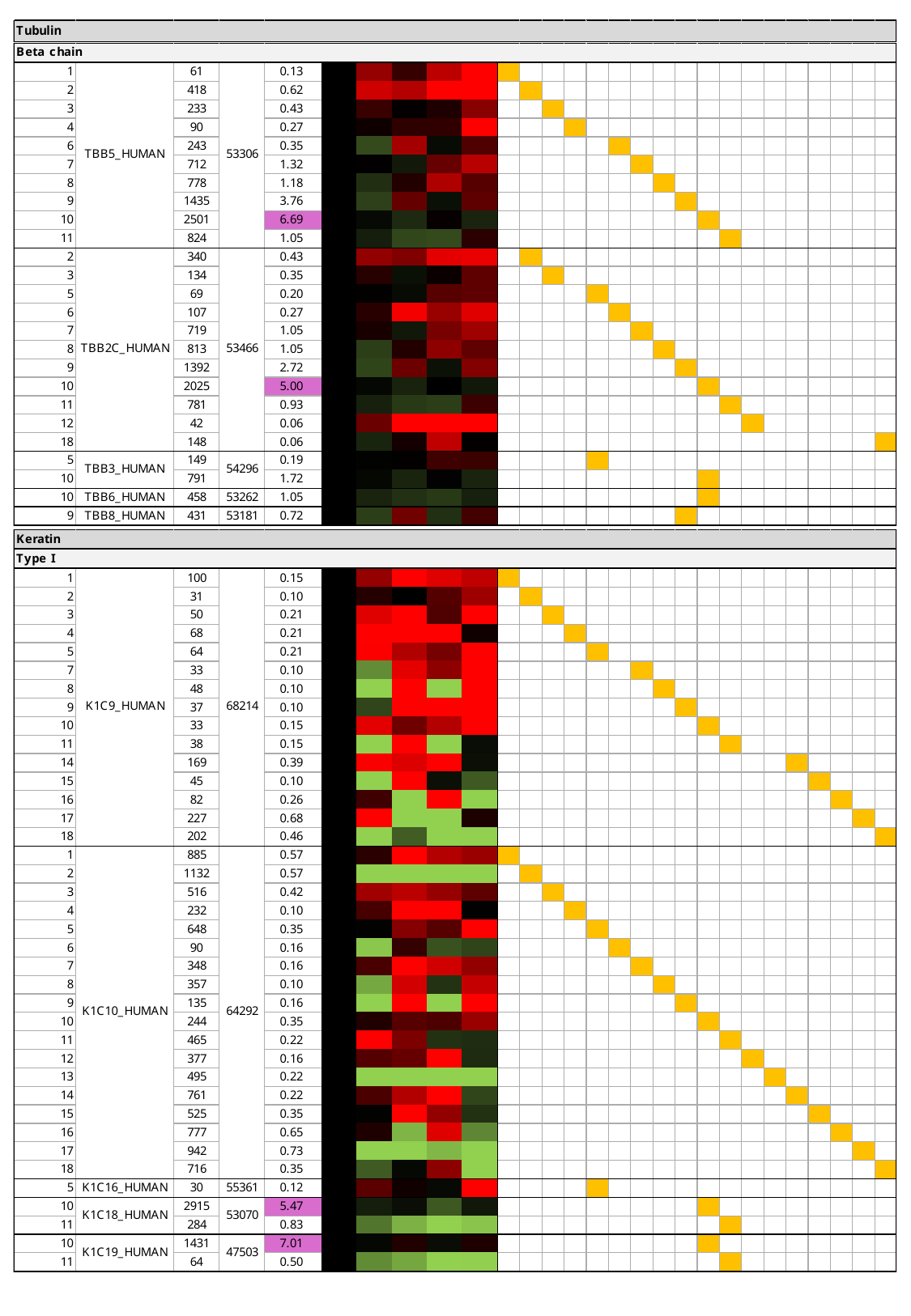

### Slide 16
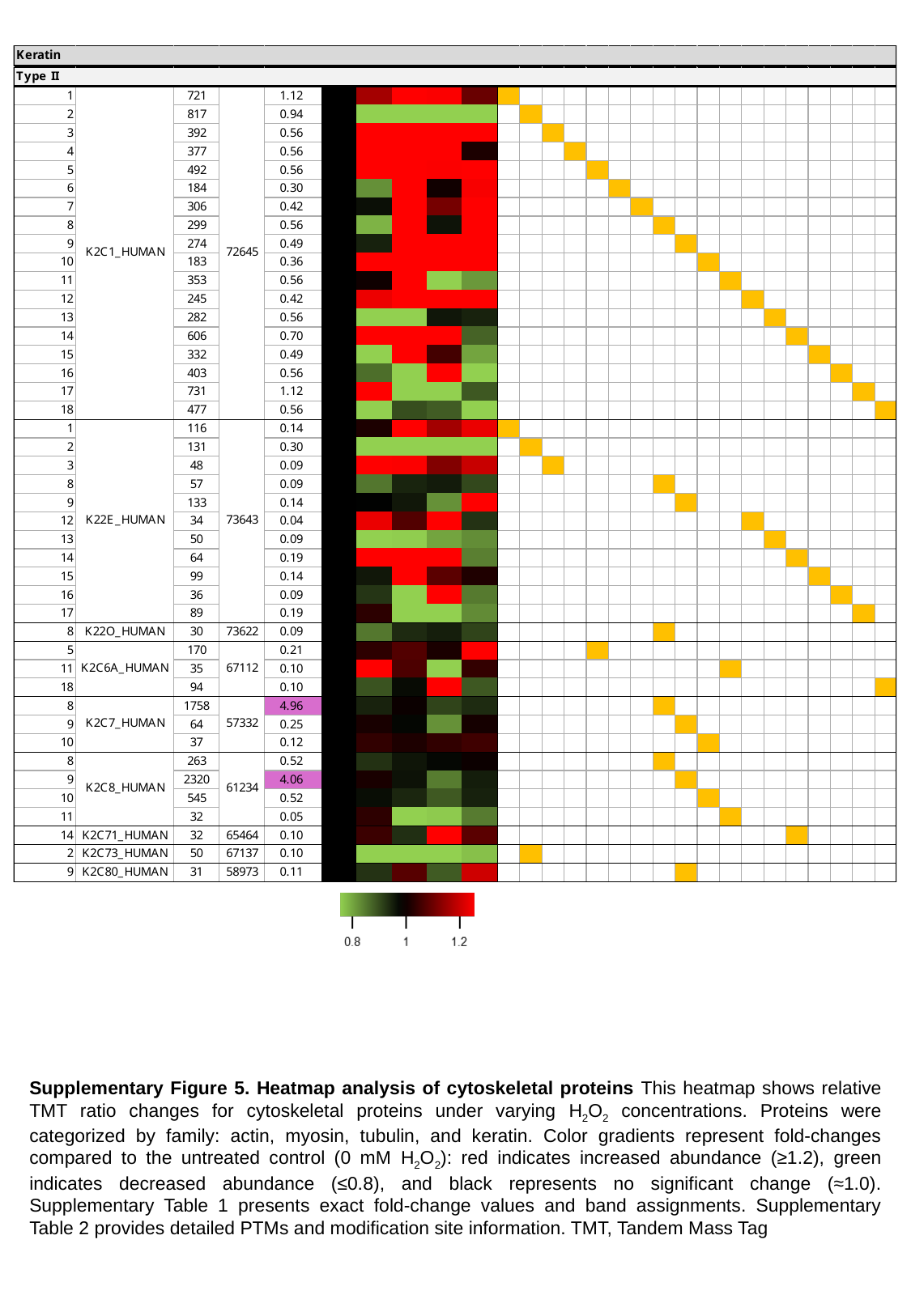

Supplementary Figure 5. Heatmap analysis of cytoskeletal proteins This heatmap shows relative TMT ratio changes for cytoskeletal proteins under varying H_2_O_2_ concentrations. Proteins were categorized by family: actin, myosin, tubulin, and keratin. Color gradients represent fold-changes compared to the untreated control (0 mM H_2_O_2_): red indicates increased abundance (≥1.2), green indicates decreased abundance (≤0.8), and black represents no significant change (≈1.0). Supplementary Table 1 presents exact fold-change values and band assignments. Supplementary Table 2 provides detailed PTMs and modification site information. TMT, Tandem Mass Tag

### Slide 17
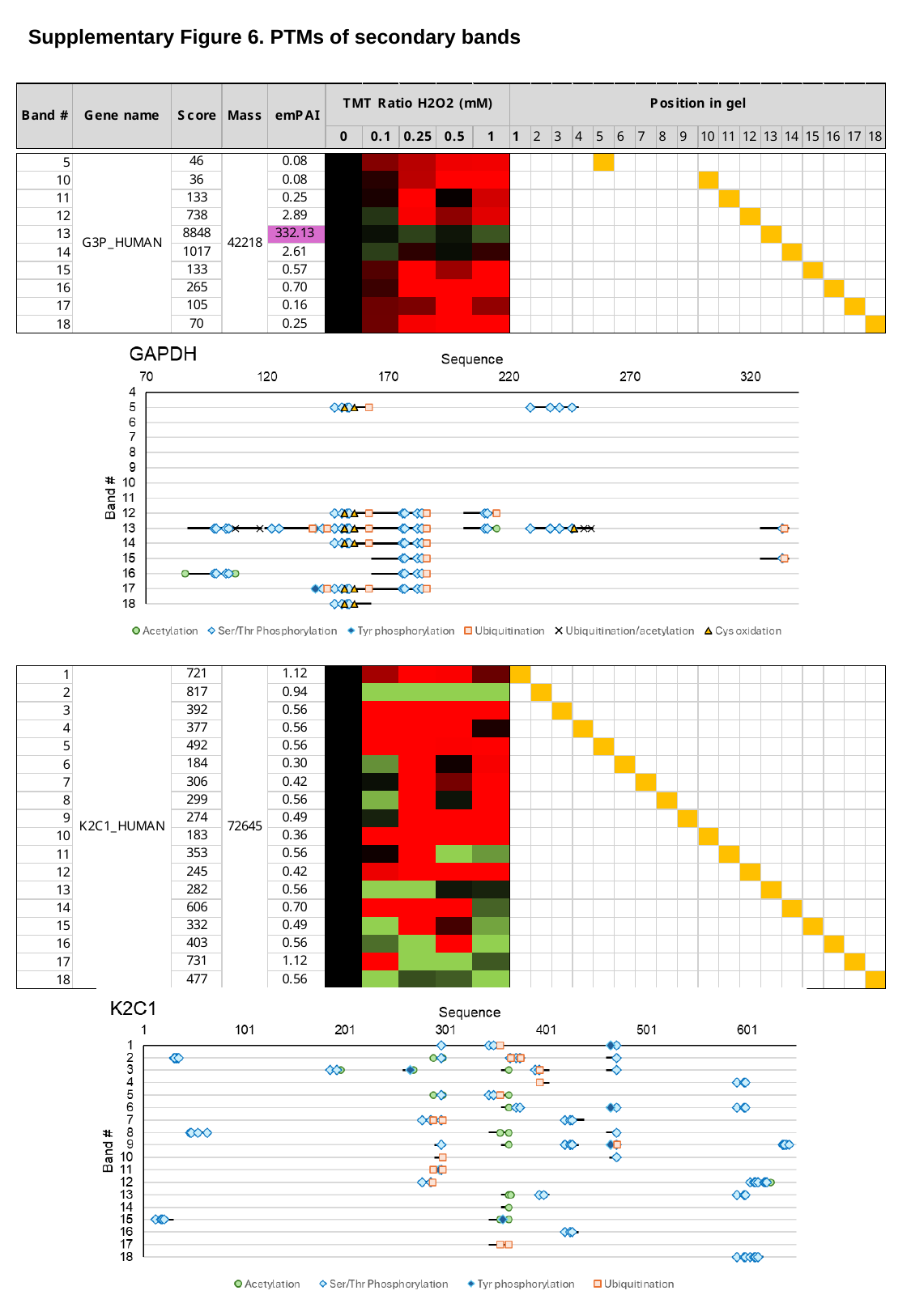

Supplementary Figure 6. PTMs of secondary bands

### Slide 18
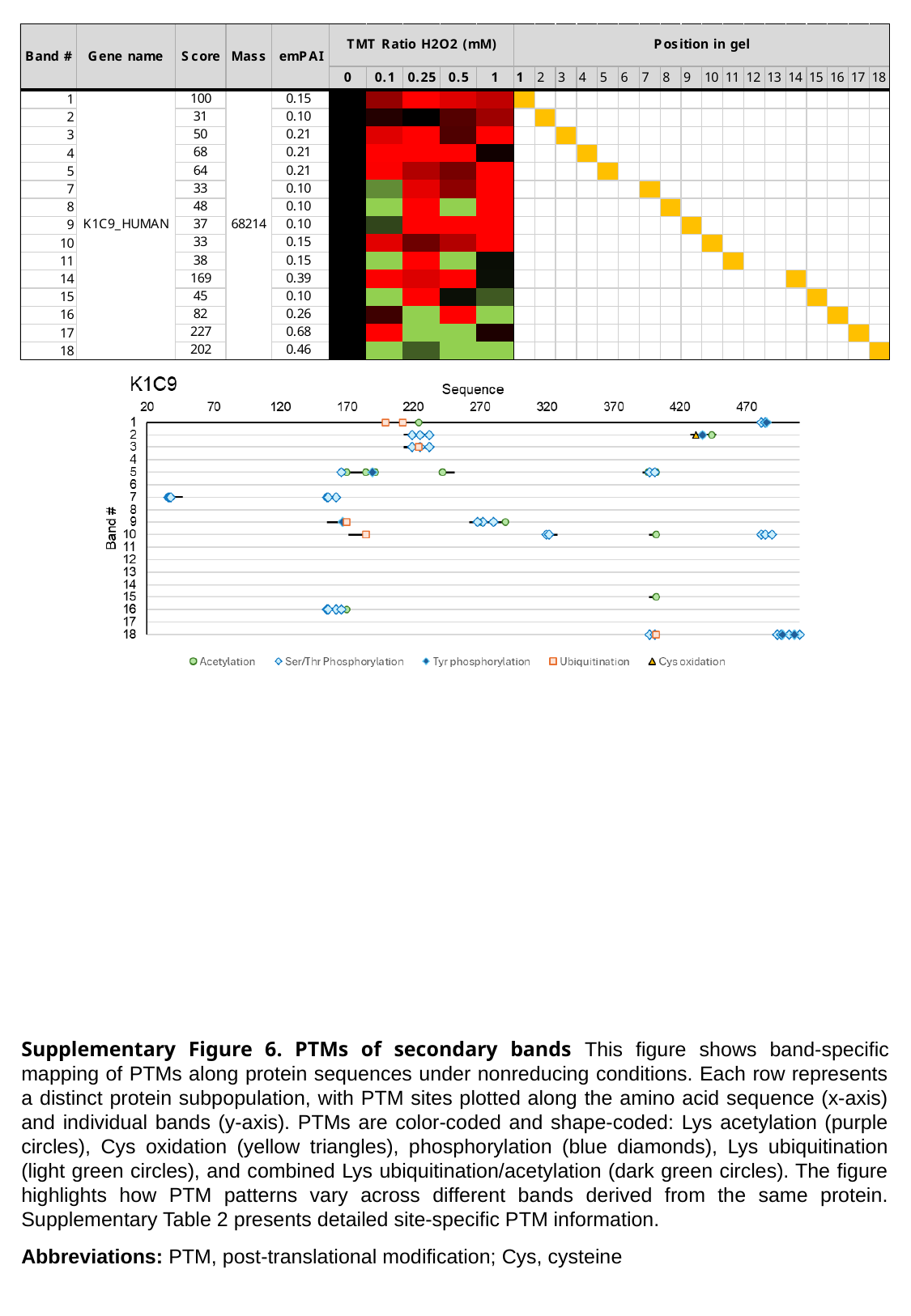

Supplementary Figure 6. PTMs of secondary bands This figure shows band-specific mapping of PTMs along protein sequences under nonreducing conditions. Each row represents a distinct protein subpopulation, with PTM sites plotted along the amino acid sequence (x-axis) and individual bands (y-axis). PTMs are color-coded and shape-coded: Lys acetylation (purple circles), Cys oxidation (yellow triangles), phosphorylation (blue diamonds), Lys ubiquitination (light green circles), and combined Lys ubiquitination/acetylation (dark green circles). The figure highlights how PTM patterns vary across different bands derived from the same protein. Supplementary Table 2 presents detailed site-specific PTM information.
Abbreviations: PTM, post-translational modification; Cys, cysteine

### Slide 19
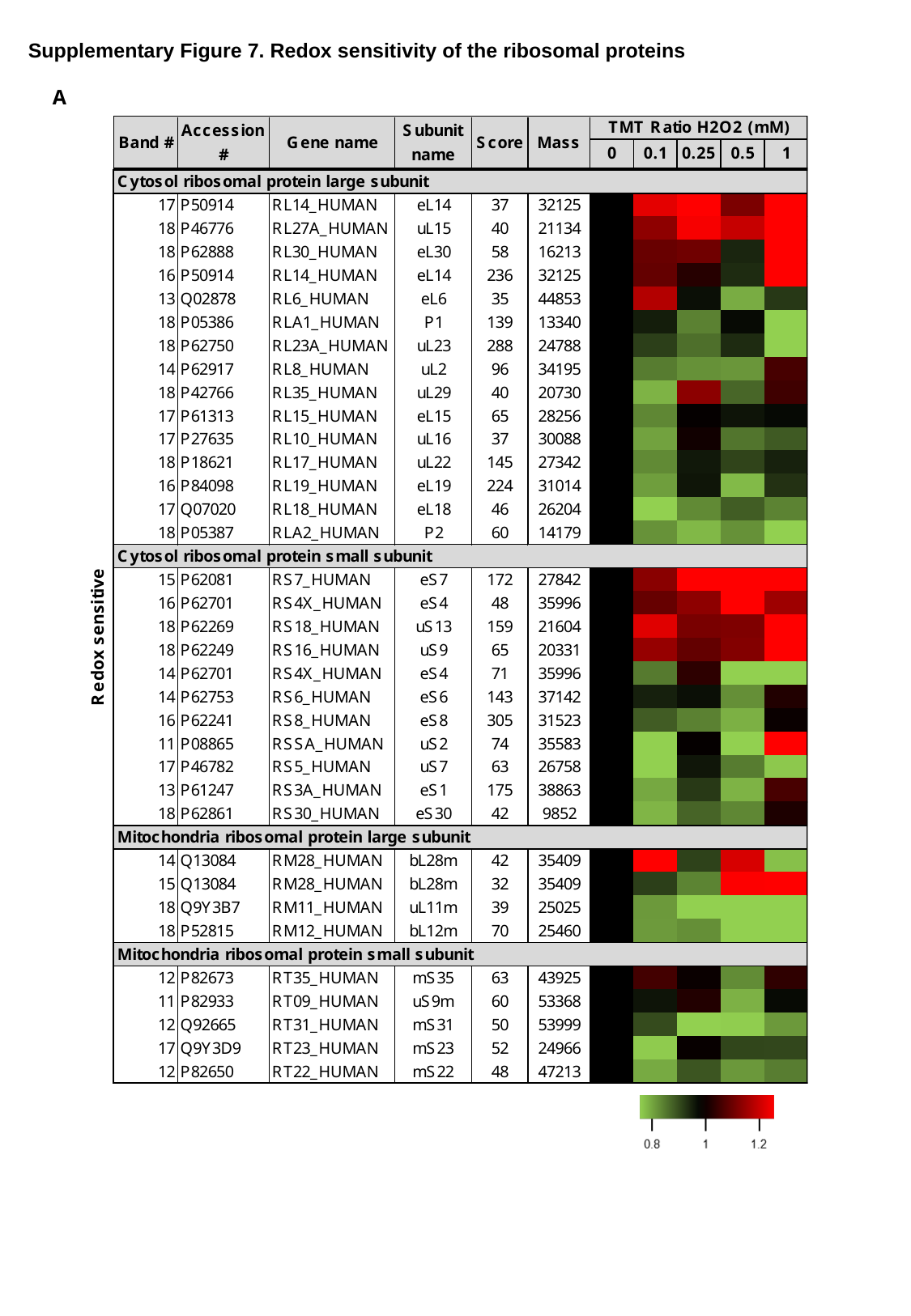

Supplementary Figure 7. Redox sensitivity of the ribosomal proteins
A

### Slide 20
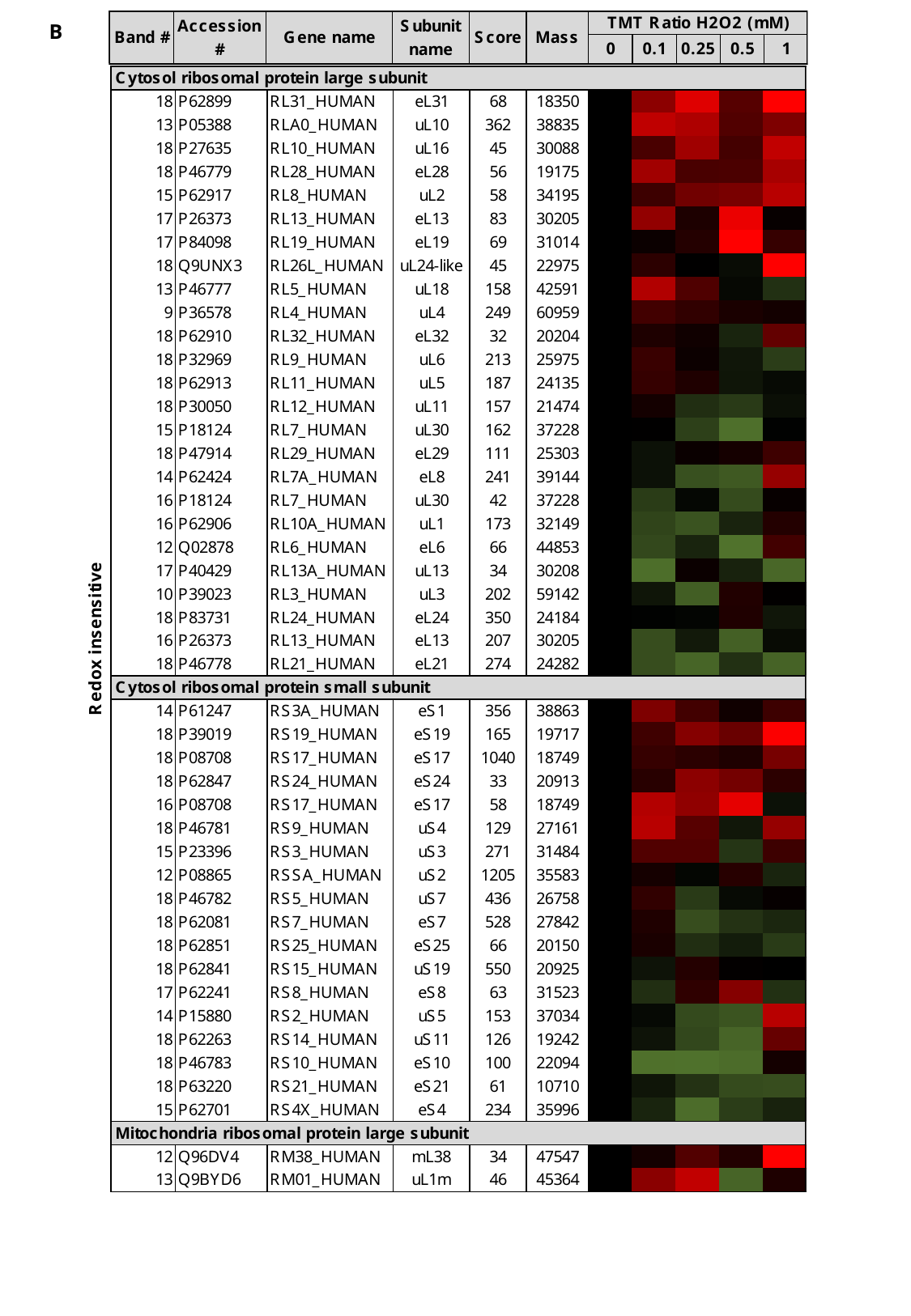

B

### Slide 21
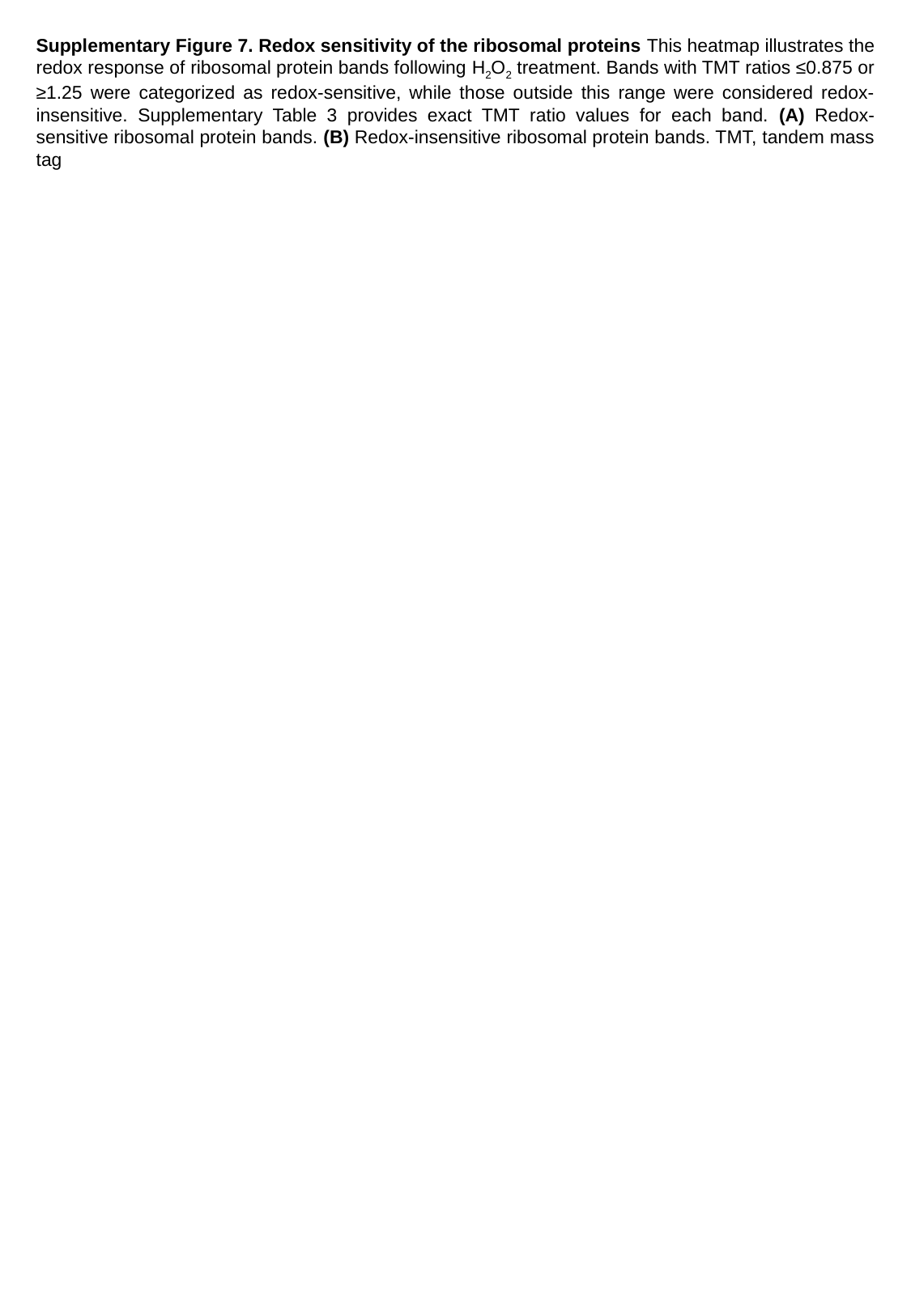

Supplementary Figure 7. Redox sensitivity of the ribosomal proteins This heatmap illustrates the redox response of ribosomal protein bands following H_2_O_2_ treatment. Bands with TMT ratios ≤0.875 or ≥1.25 were categorized as redox-sensitive, while those outside this range were considered redox-insensitive. Supplementary Table 3 provides exact TMT ratio values for each band. (A) Redox-sensitive ribosomal protein bands. (B) Redox-insensitive ribosomal protein bands. TMT, tandem mass tag

### Slide 22
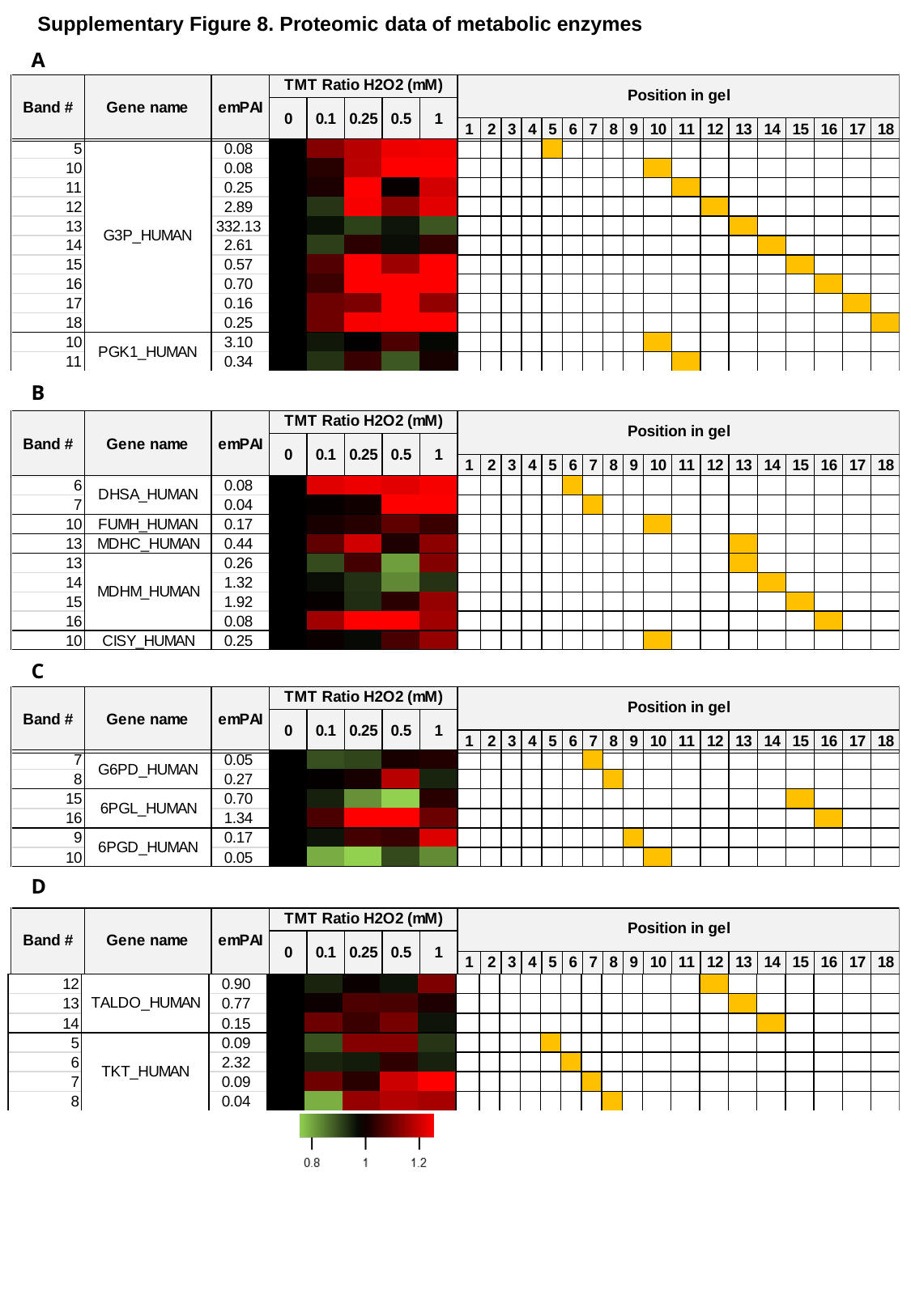

Supplementary Figure 8. Proteomic data of metabolic enzymes
A
B
C
D

### Slide 23
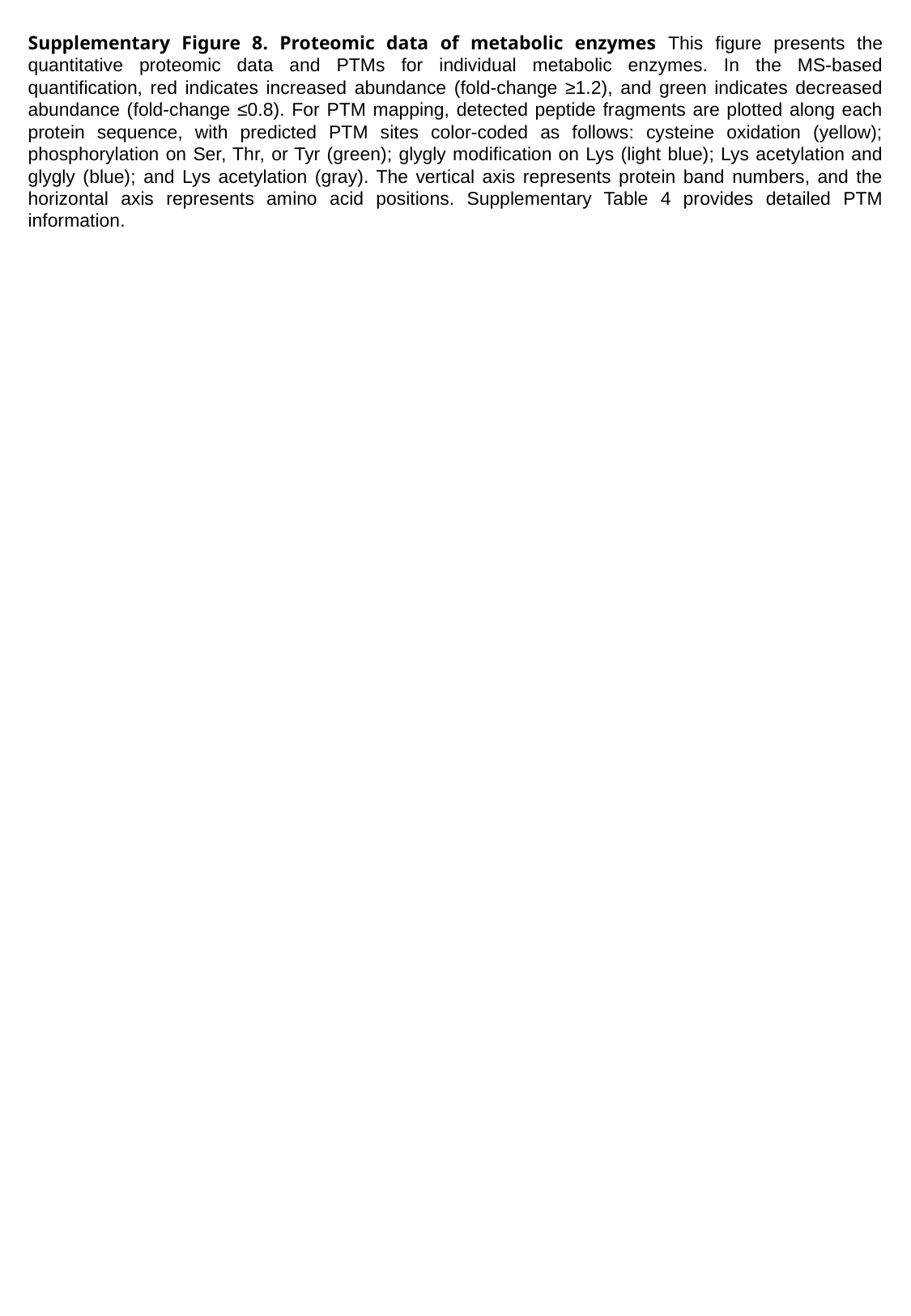

Supplementary Figure 8. Proteomic data of metabolic enzymes This figure presents the quantitative proteomic data and PTMs for individual metabolic enzymes. In the MS-based quantification, red indicates increased abundance (fold-change ≥1.2), and green indicates decreased abundance (fold-change ≤0.8). For PTM mapping, detected peptide fragments are plotted along each protein sequence, with predicted PTM sites color-coded as follows: cysteine oxidation (yellow); phosphorylation on Ser, Thr, or Tyr (green); glygly modification on Lys (light blue); Lys acetylation and glygly (blue); and Lys acetylation (gray). The vertical axis represents protein band numbers, and the horizontal axis represents amino acid positions. Supplementary Table 4 provides detailed PTM information.

### Slide 24
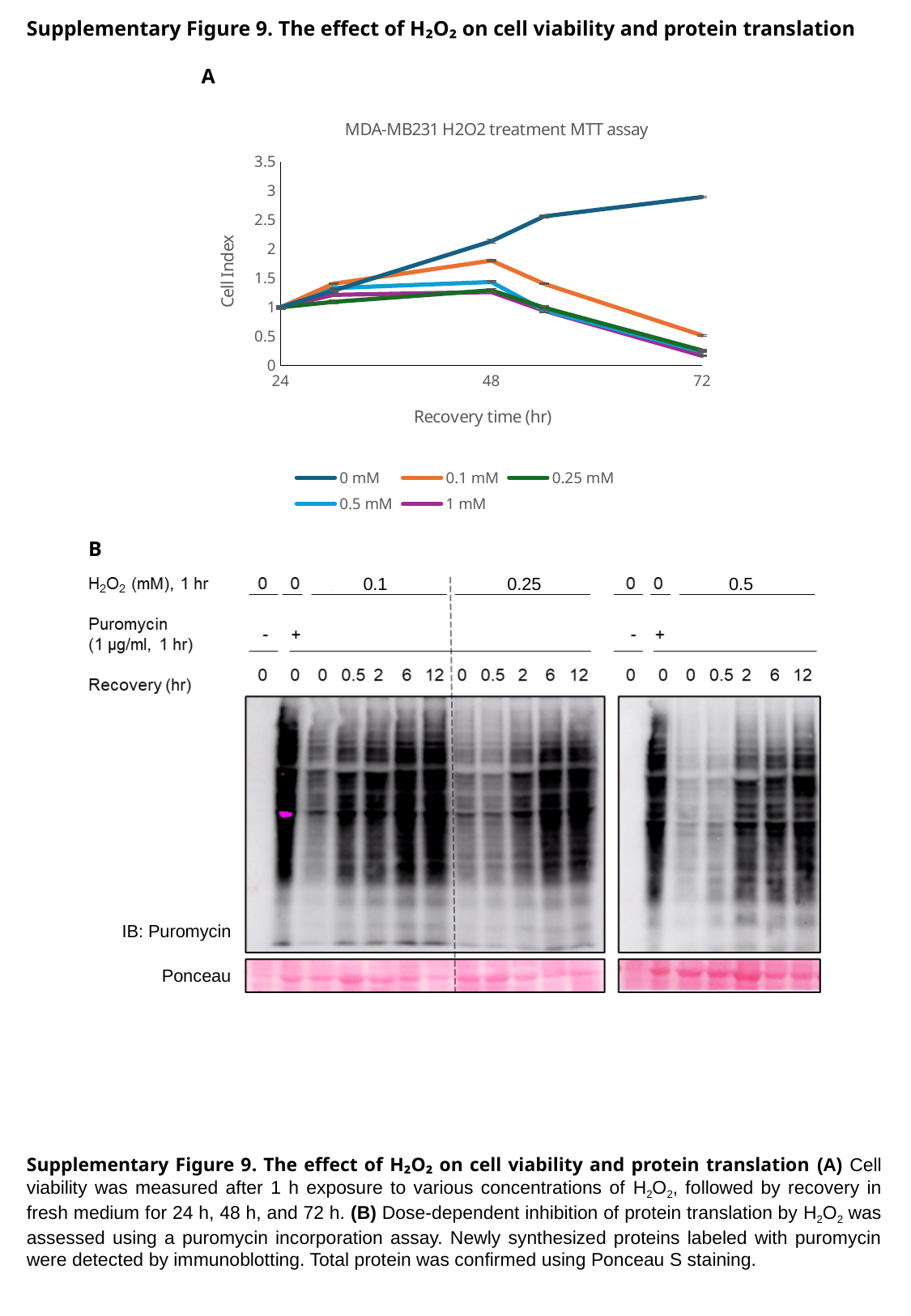

Supplementary Figure 9. The effect of H₂O₂ on cell viability and protein translation
A
#### Chart: MDA-MB231 H2O2 treatment MTT assay
| Category | 0 mM | 0.1 mM | 0.25 mM | 0.5 mM | 1 mM |
|---|---|---|---|---|---|B
0.1
0.25
0.5
IB: Puromycin
Ponceau
Supplementary Figure 9. The effect of H₂O₂ on cell viability and protein translation (A) Cell viability was measured after 1 h exposure to various concentrations of H_2_O_2_, followed by recovery in fresh medium for 24 h, 48 h, and 72 h. (B) Dose-dependent inhibition of protein translation by H_2_O_2_ was assessed using a puromycin incorporation assay. Newly synthesized proteins labeled with puromycin were detected by immunoblotting. Total protein was confirmed using Ponceau S staining.

### Slide 25
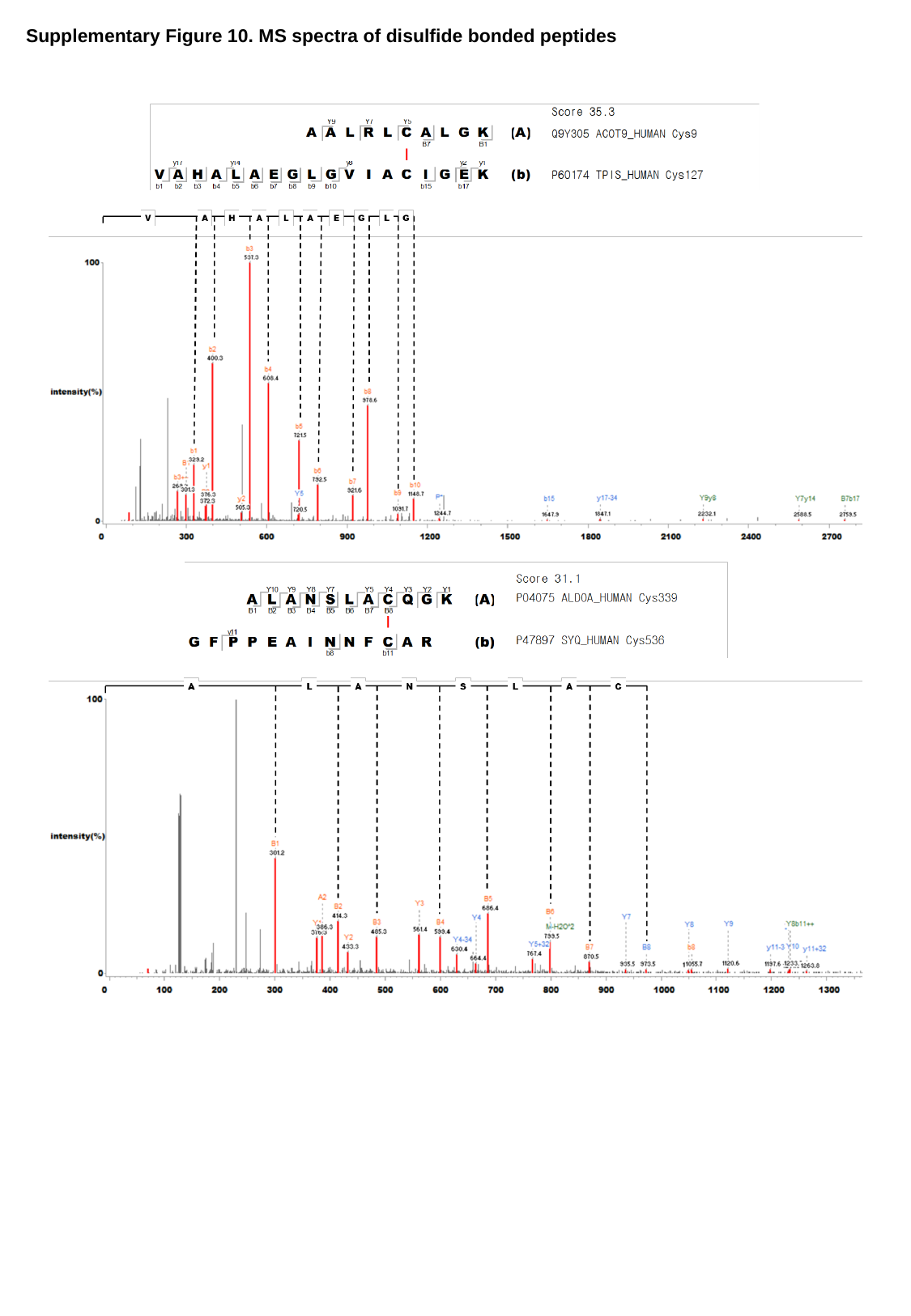

Supplementary Figure 10. MS spectra of disulfide bonded peptides

### Slide 26
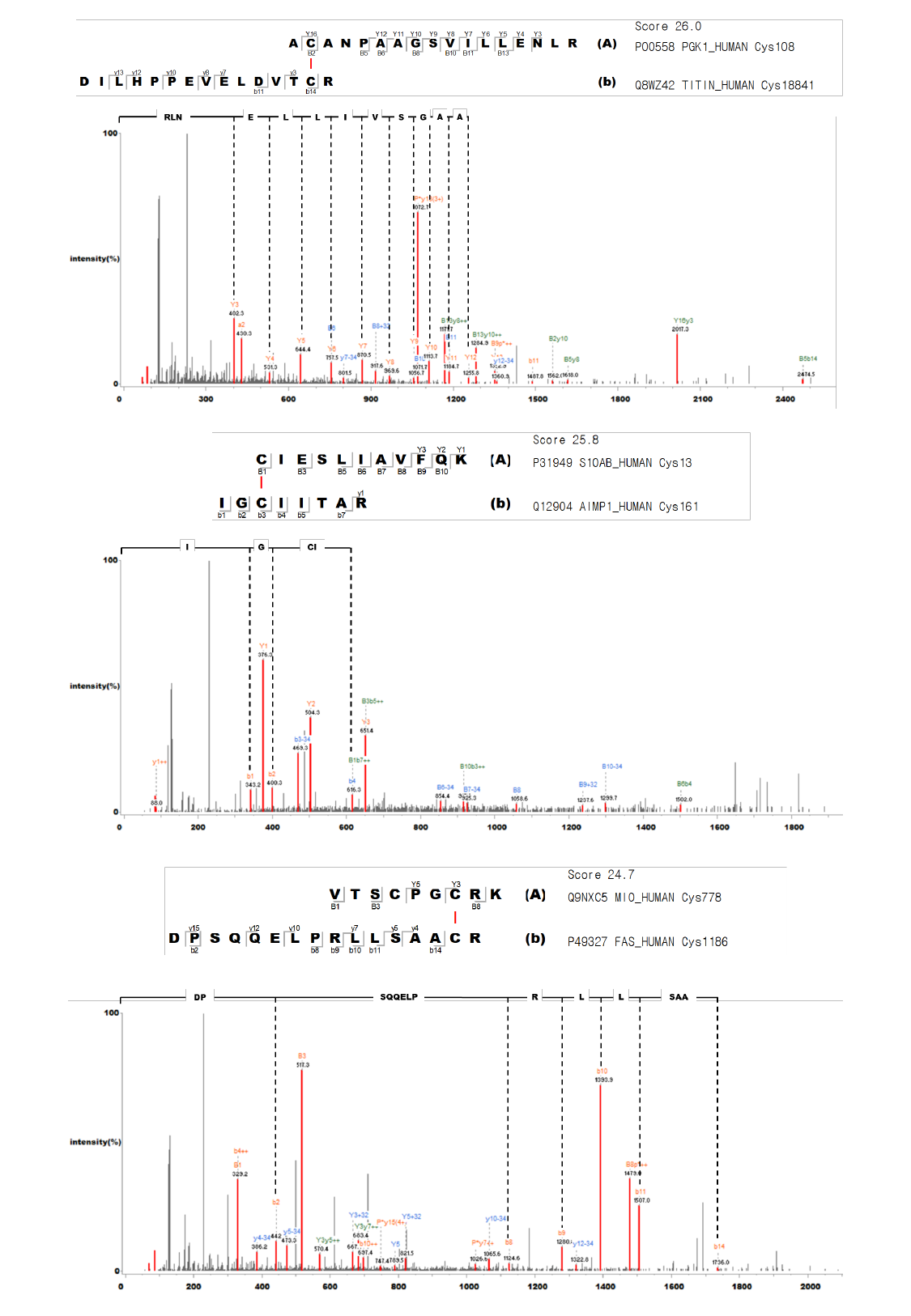

### Slide 27
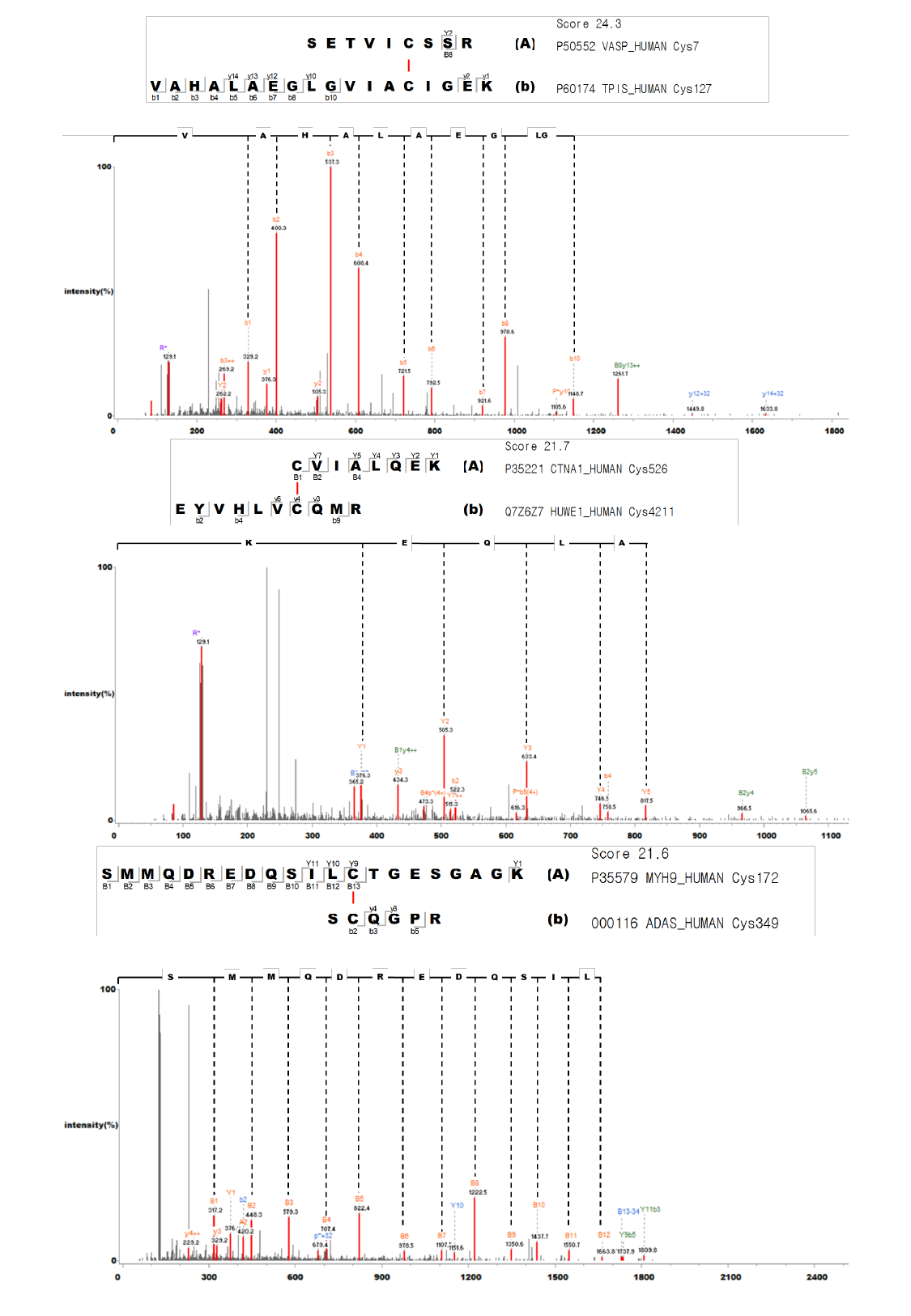

### Slide 28
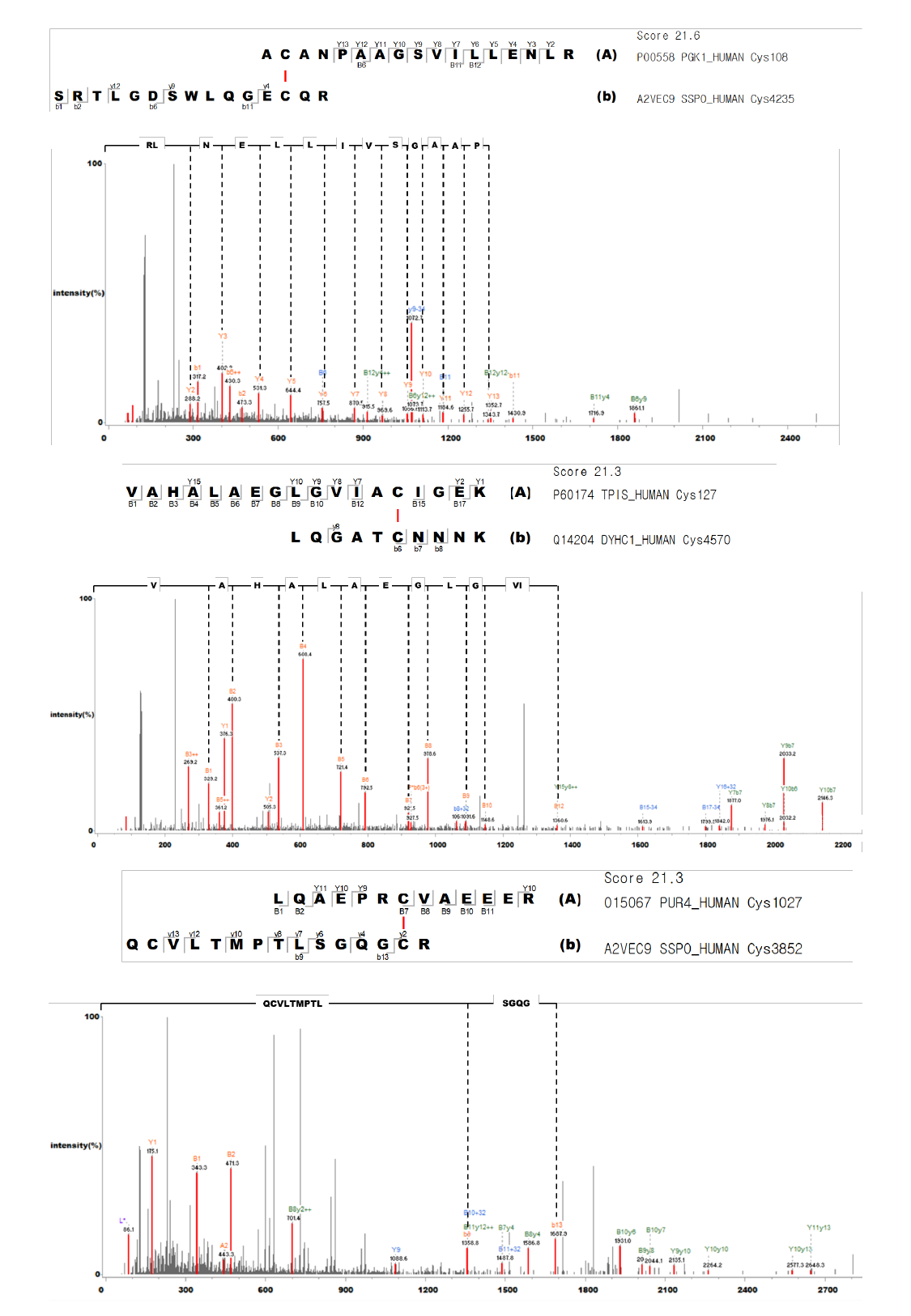

### Slide 29
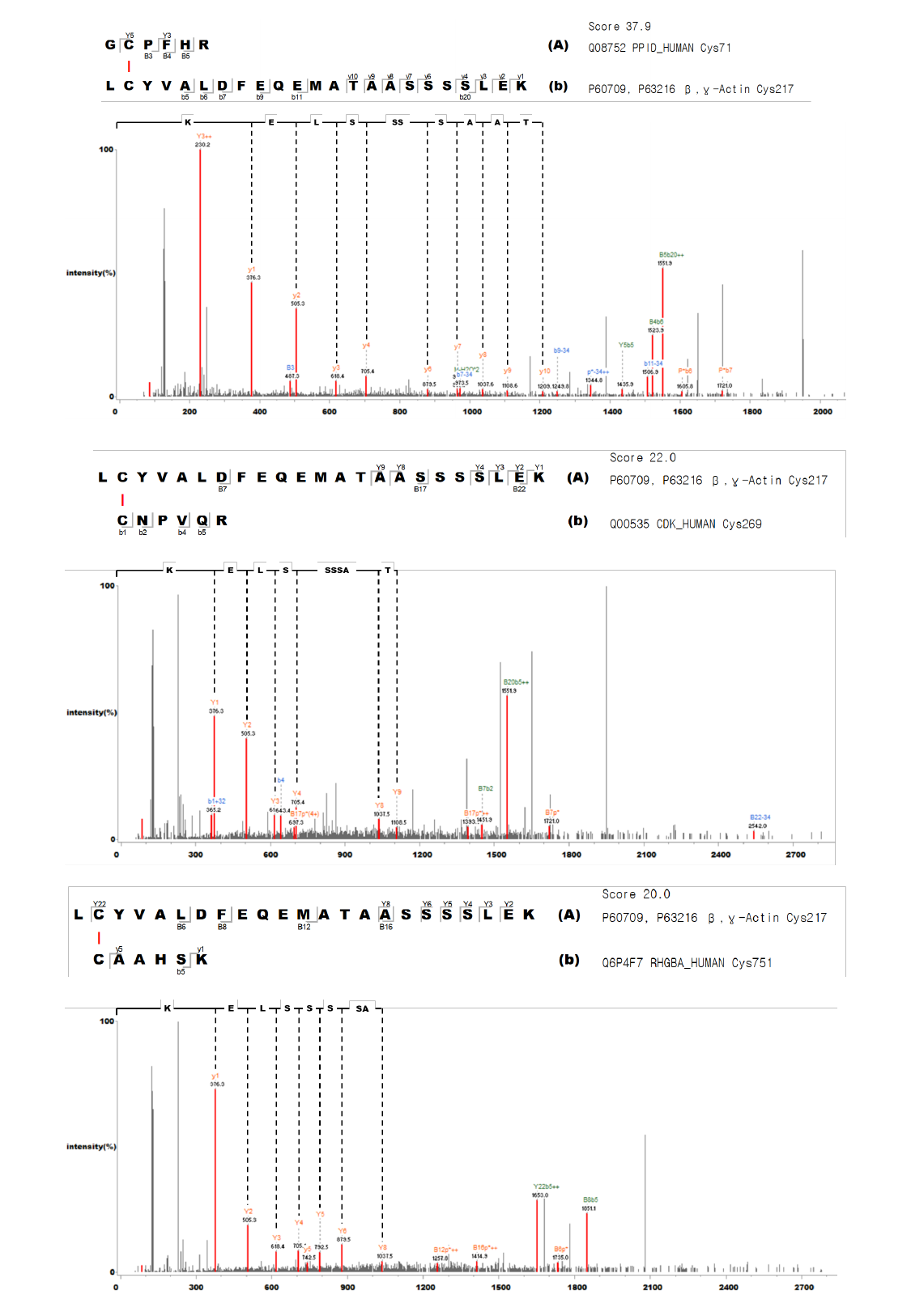

### Slide 30
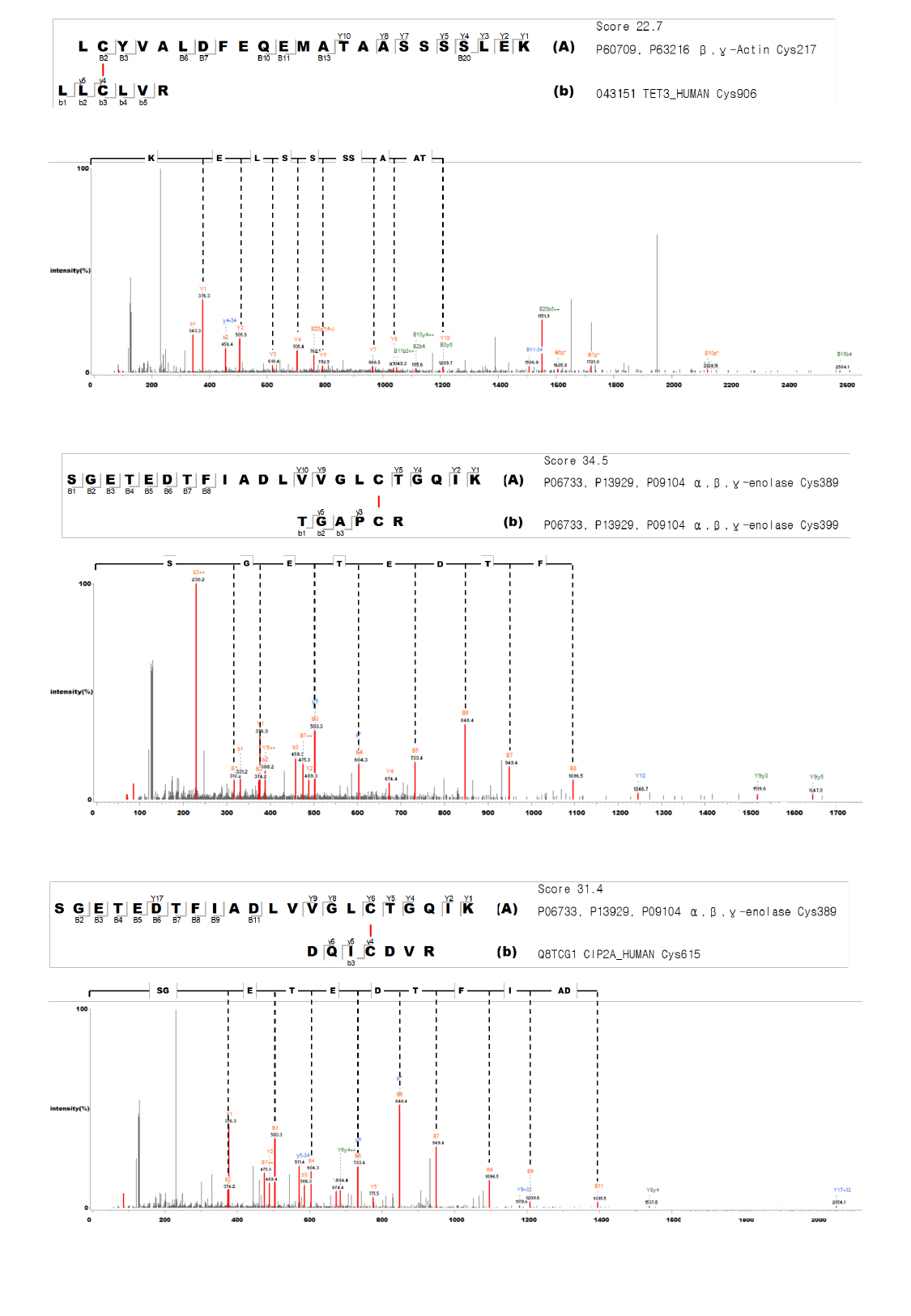

### Slide 31
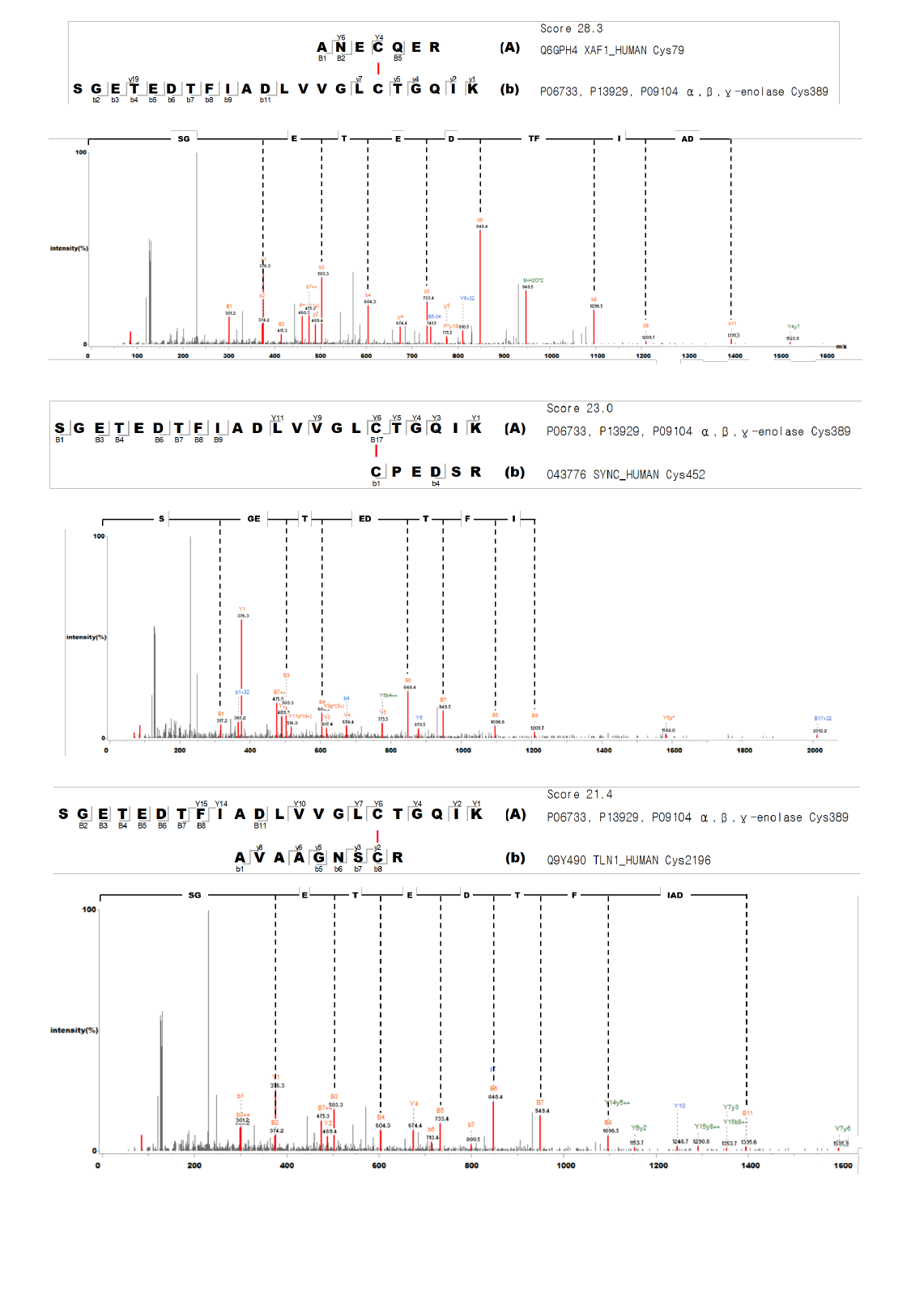

### Slide 32
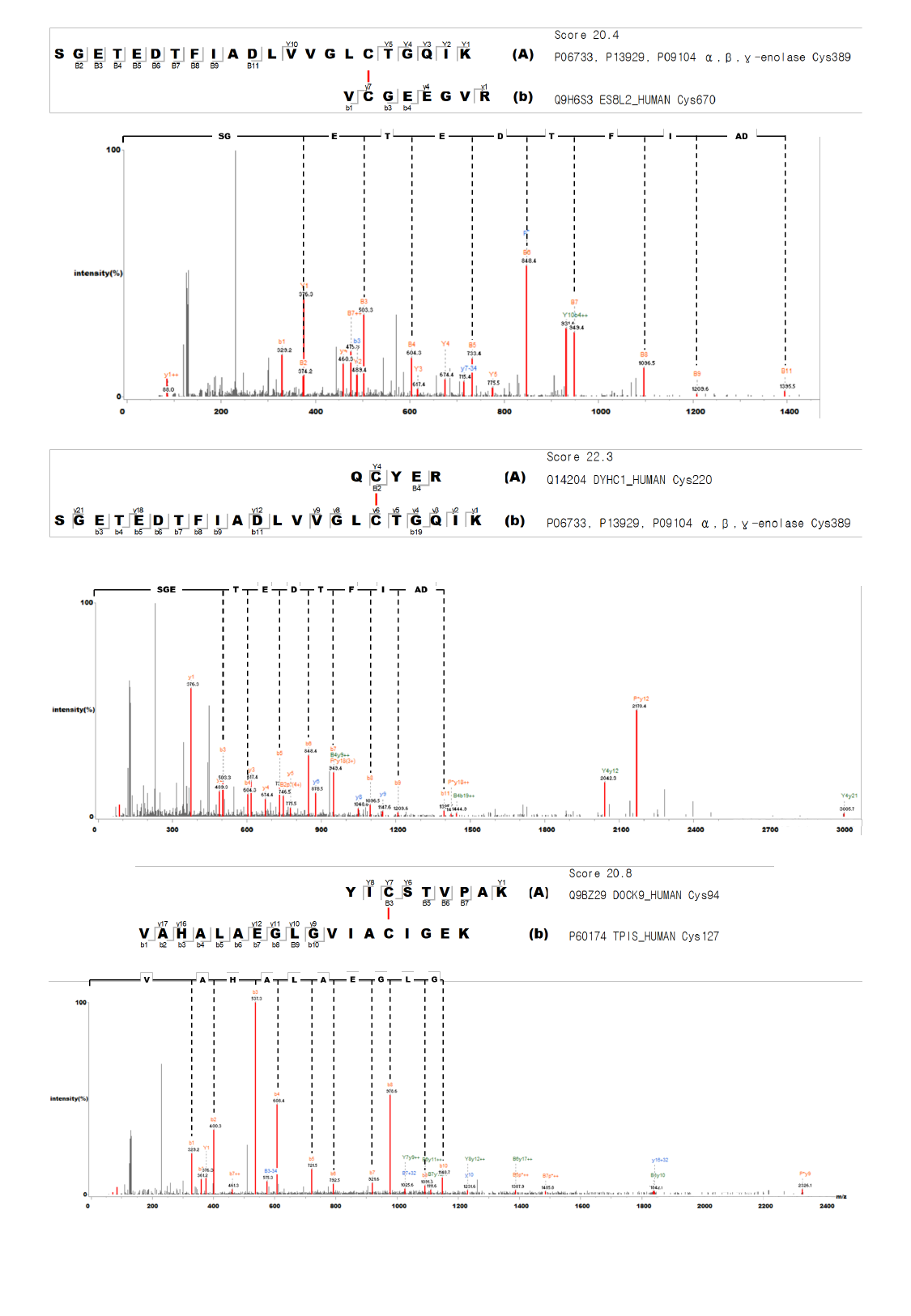

### Slide 33
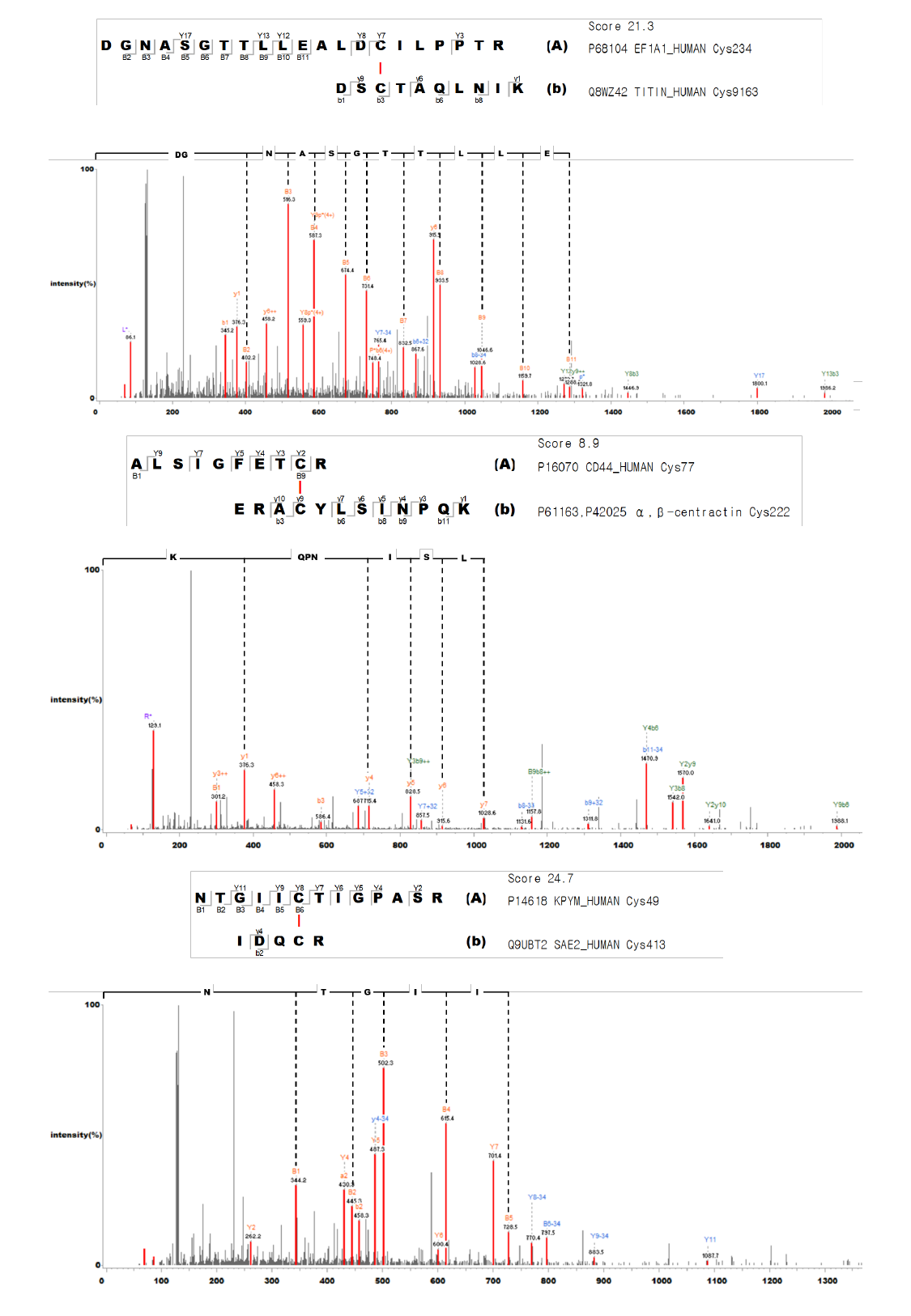

### Slide 34
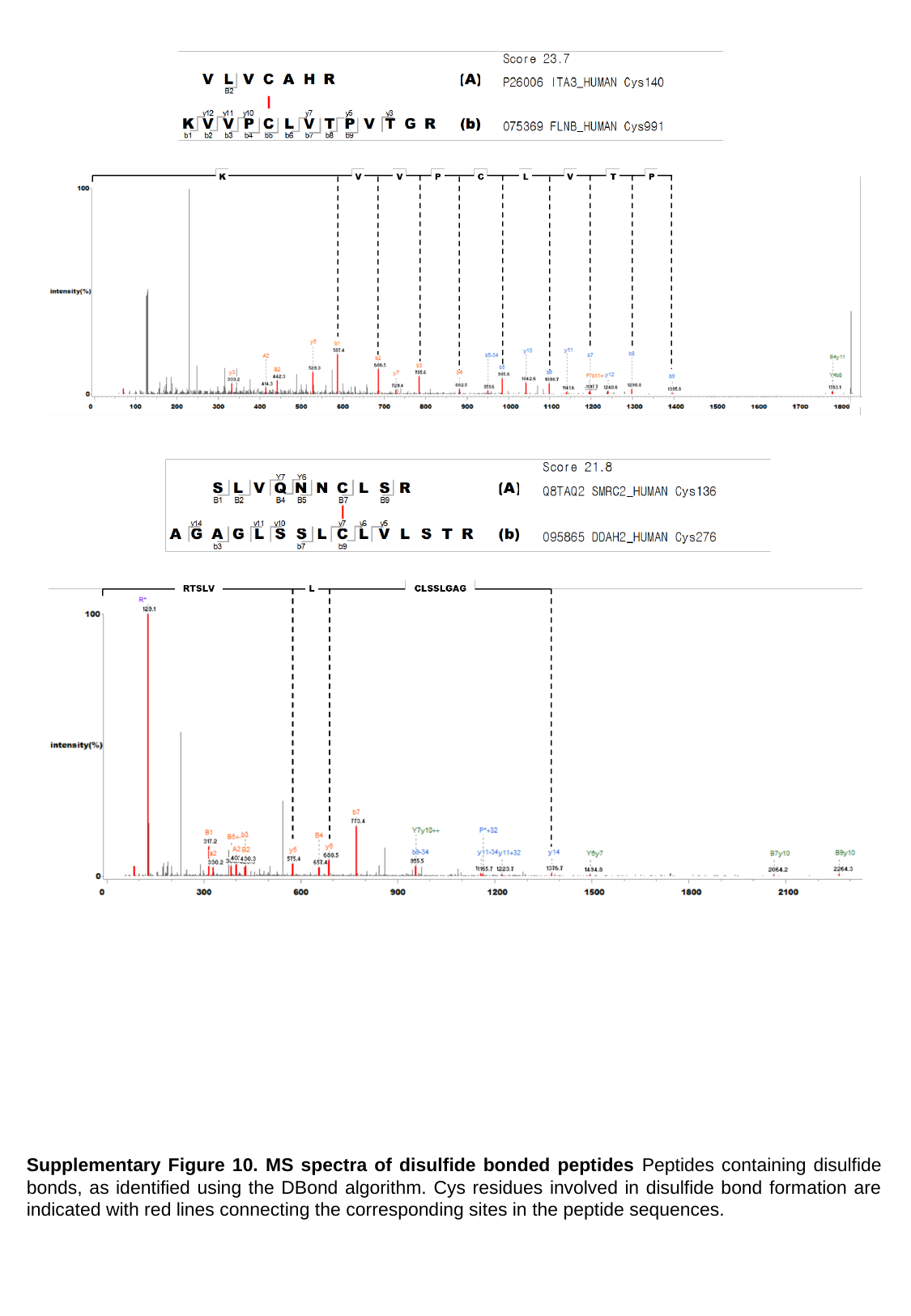

Supplementary Figure 10. MS spectra of disulfide bonded peptides Peptides containing disulfide bonds, as identified using the DBond algorithm. Cys residues involved in disulfide bond formation are indicated with red lines connecting the corresponding sites in the peptide sequences.

### Slide 35
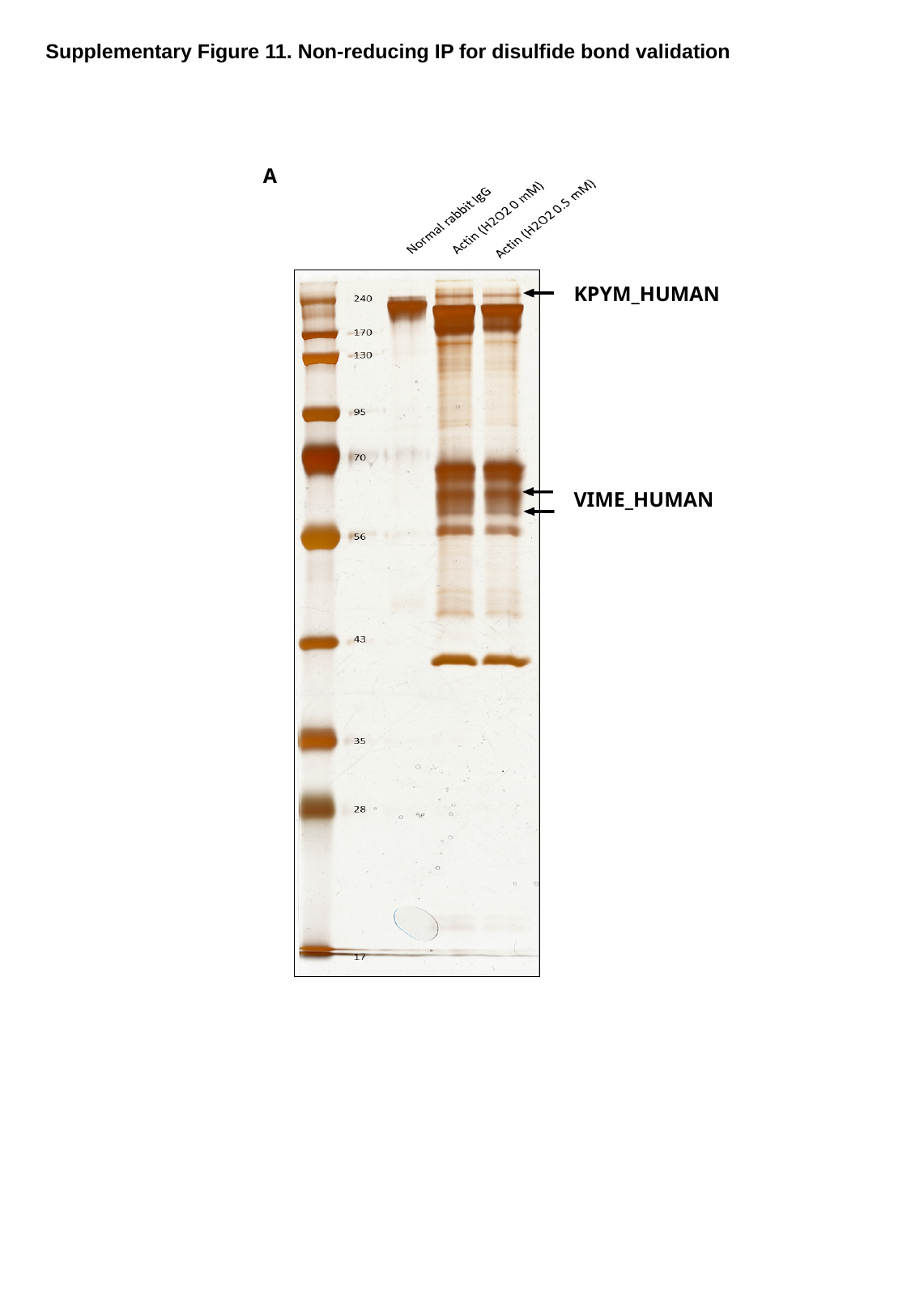

Supplementary Figure 11. Non-reducing IP for disulfide bond validation
A
KPYM_HUMAN
VIME_HUMAN

### Slide 36
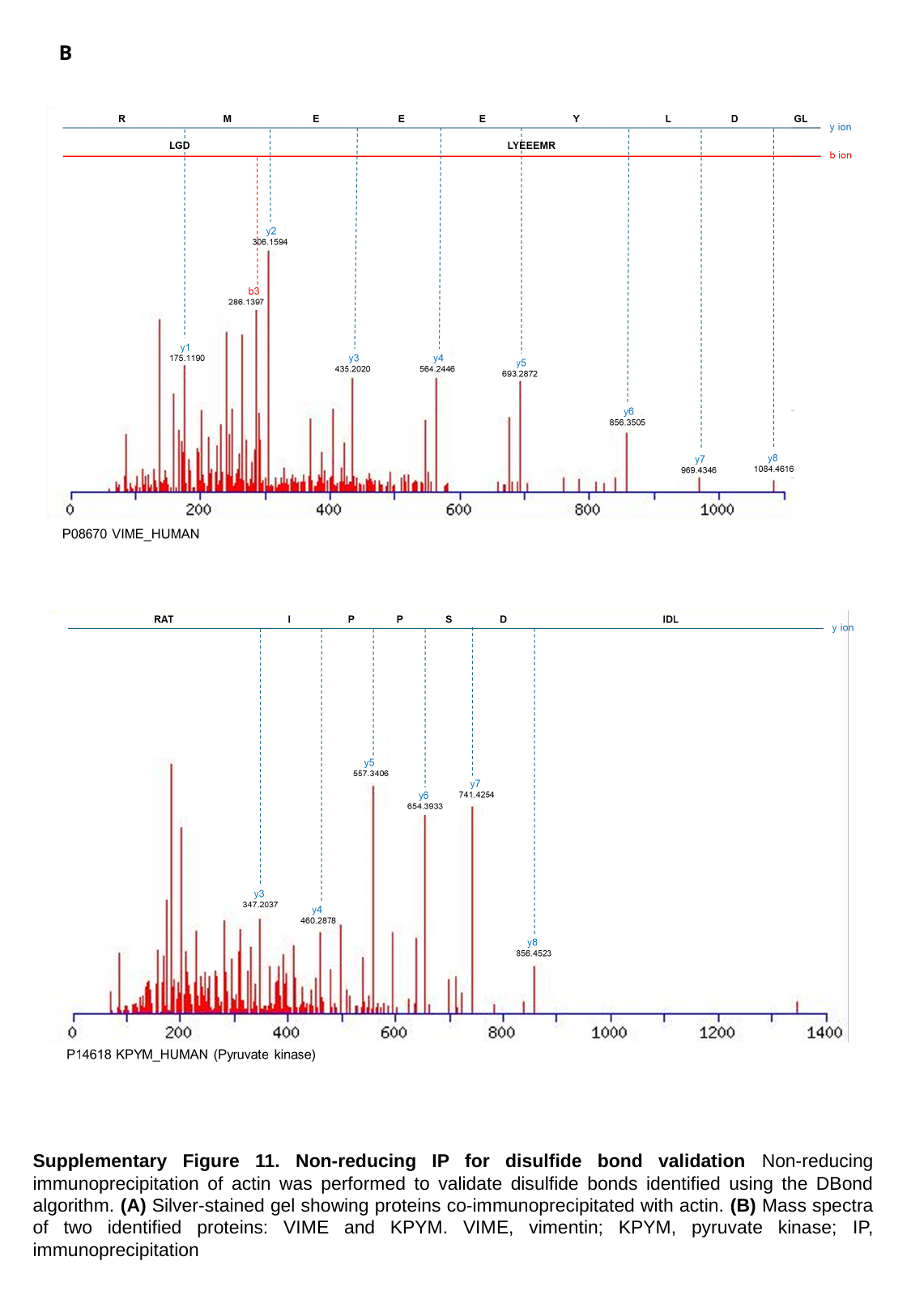

B
Supplementary Figure 11. Non-reducing IP for disulfide bond validation Non-reducing immunoprecipitation of actin was performed to validate disulfide bonds identified using the DBond algorithm. (A) Silver-stained gel showing proteins co-immunoprecipitated with actin. (B) Mass spectra of two identified proteins: VIME and KPYM. VIME, vimentin; KPYM, pyruvate kinase; IP, immunoprecipitation

### Slide 37
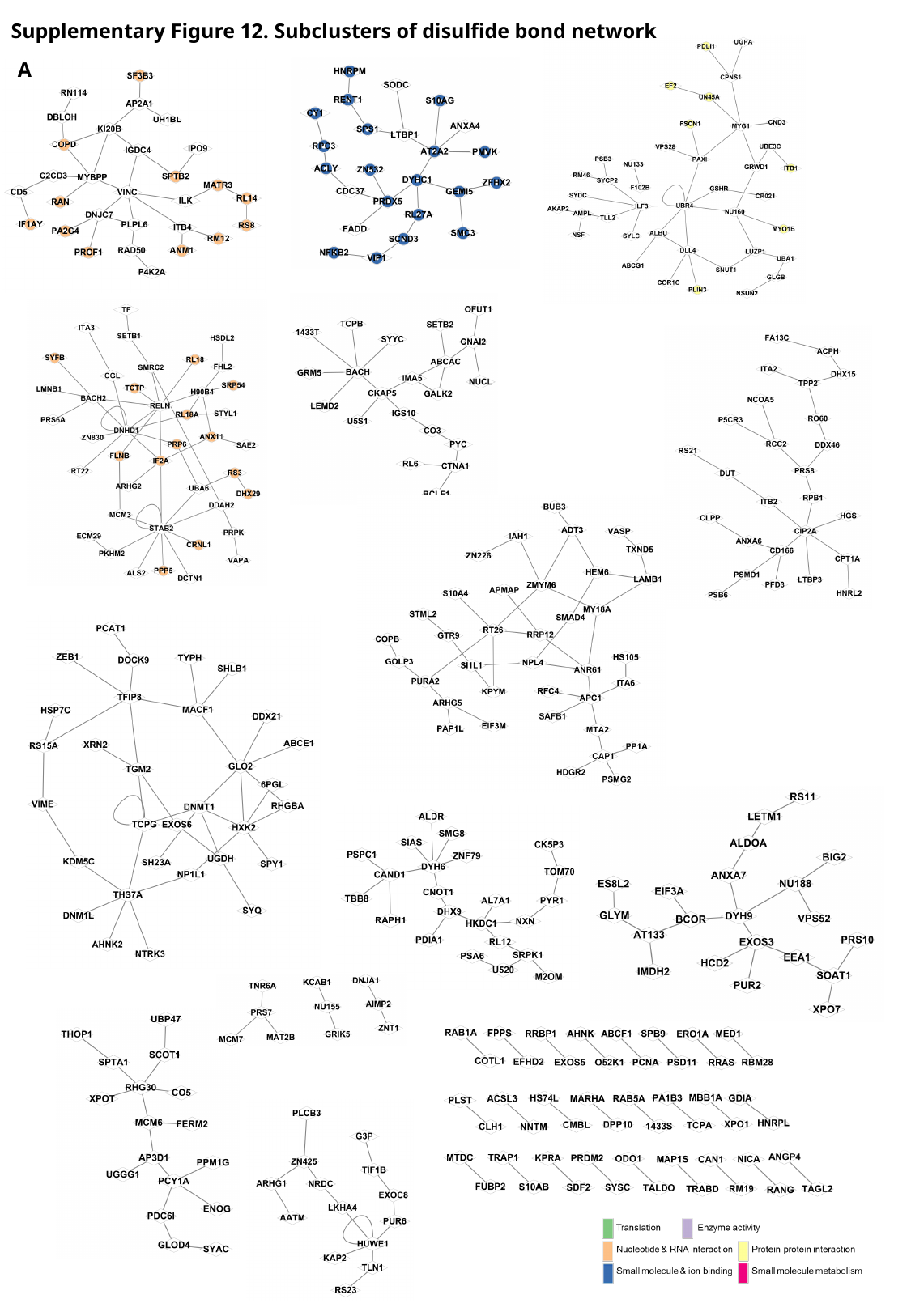

Supplementary Figure 12. Subclusters of disulfide bond network
A

### Slide 38
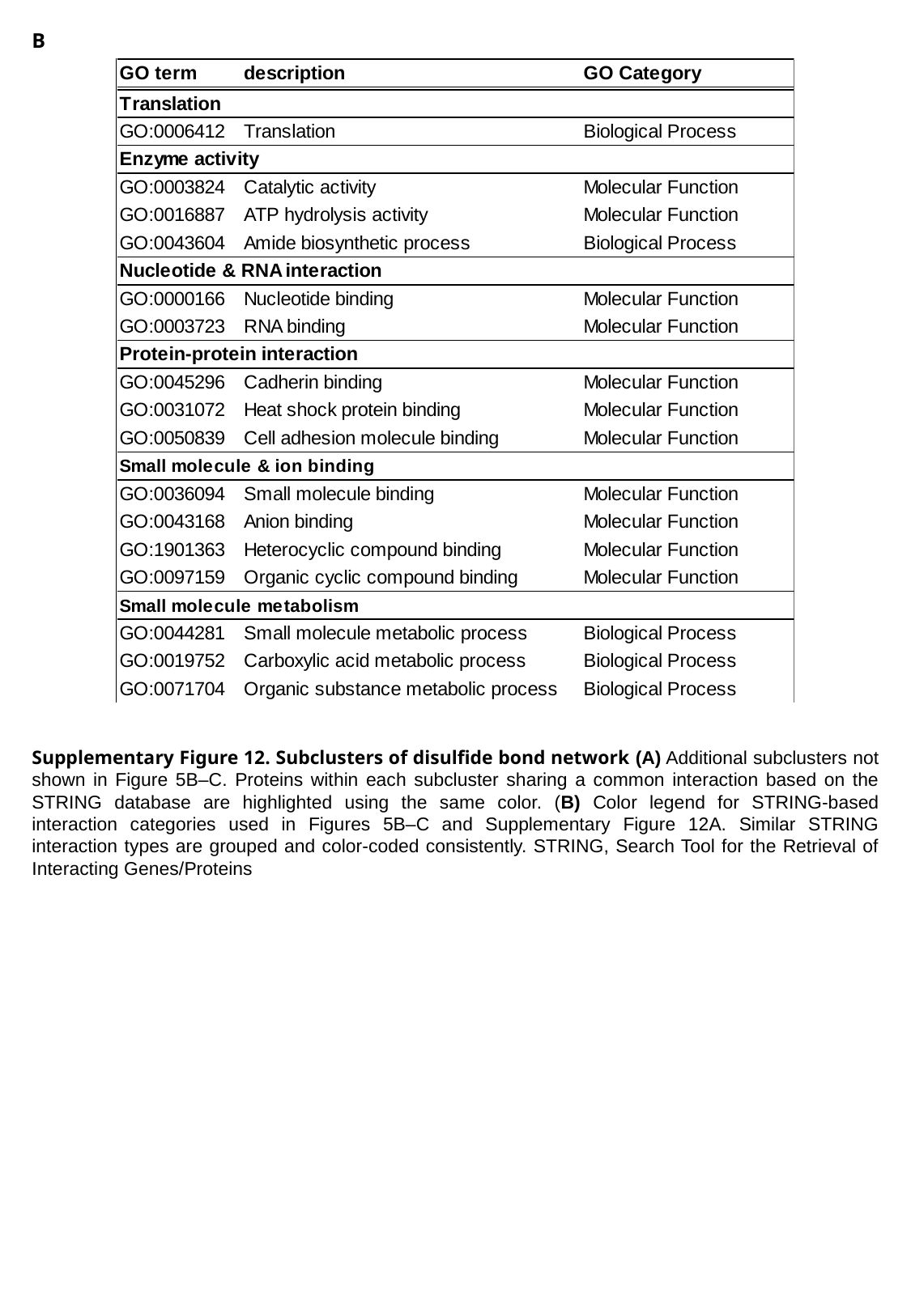

B
Supplementary Figure 12. Subclusters of disulfide bond network (A) Additional subclusters not shown in Figure 5B–C. Proteins within each subcluster sharing a common interaction based on the STRING database are highlighted using the same color. (B) Color legend for STRING-based interaction categories used in Figures 5B–C and Supplementary Figure 12A. Similar STRING interaction types are grouped and color-coded consistently. STRING, Search Tool for the Retrieval of Interacting Genes/Proteins

### Slide 39
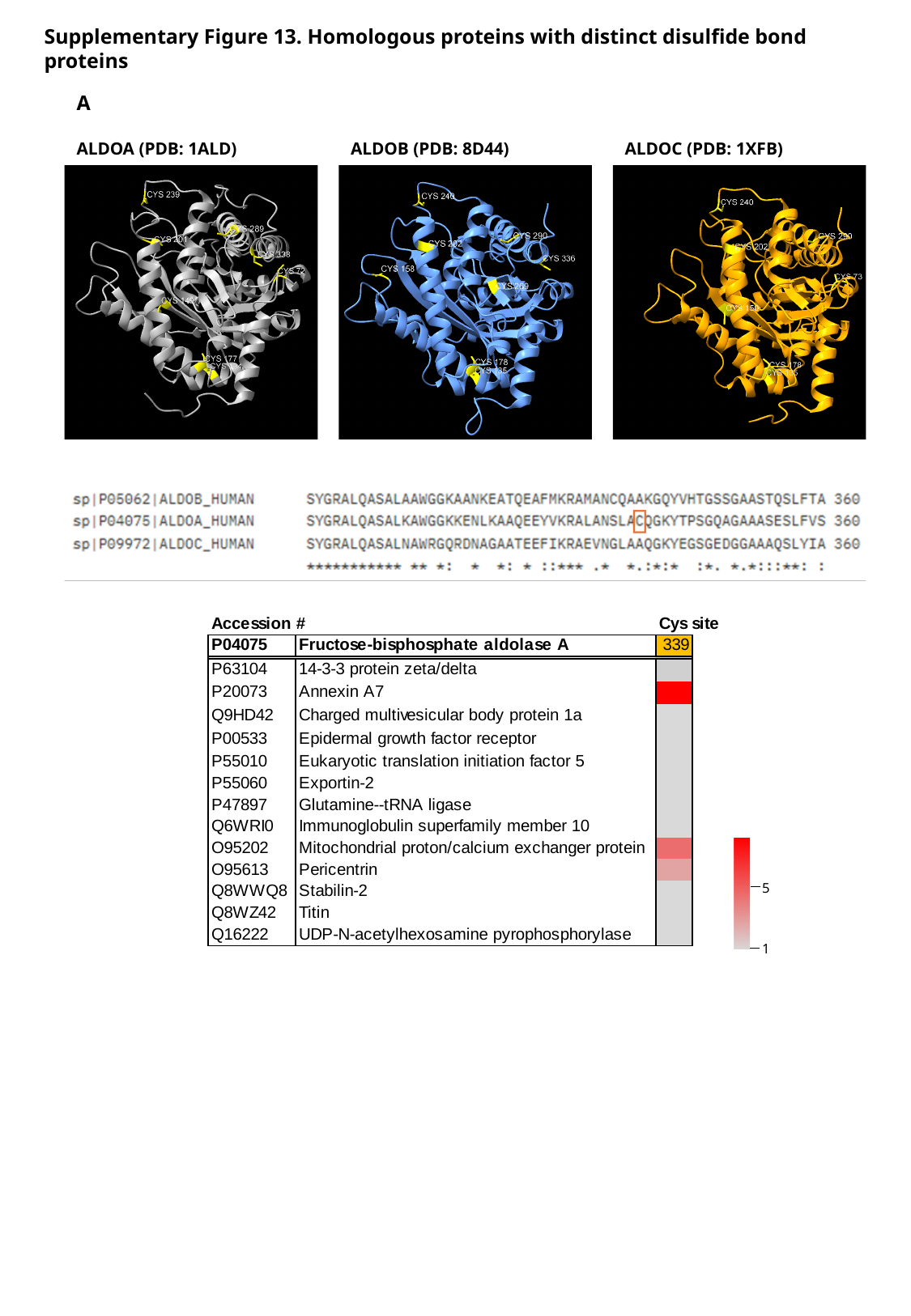

Supplementary Figure 13. Homologous proteins with distinct disulfide bond proteins
A
ALDOA (PDB: 1ALD)
ALDOC (PDB: 1XFB)
ALDOB (PDB: 8D44)
5
1

### Slide 40
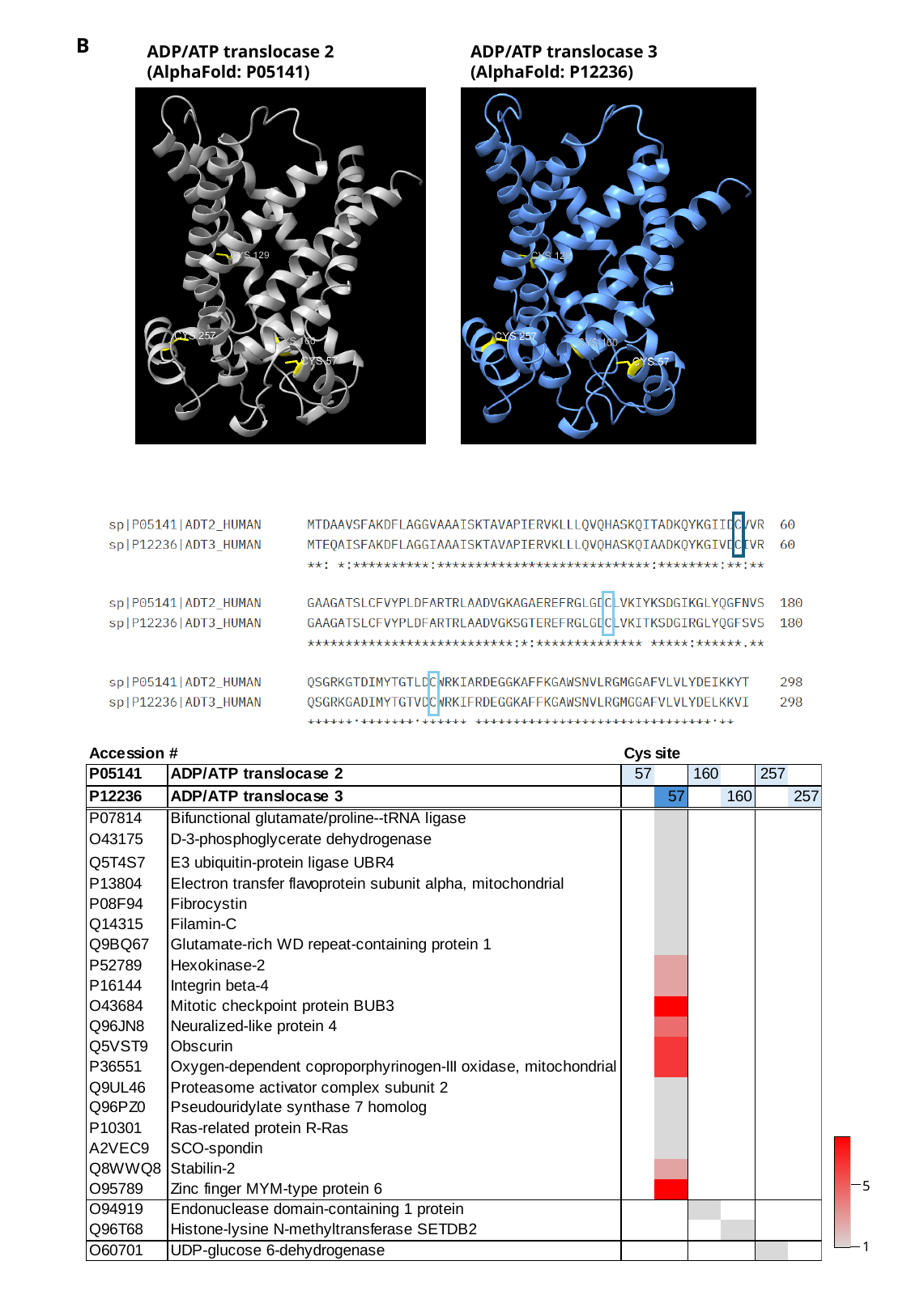

B
ADP/ATP translocase 2 (AlphaFold: P05141)
ADP/ATP translocase 3 (AlphaFold: P12236)
5
1

### Slide 41
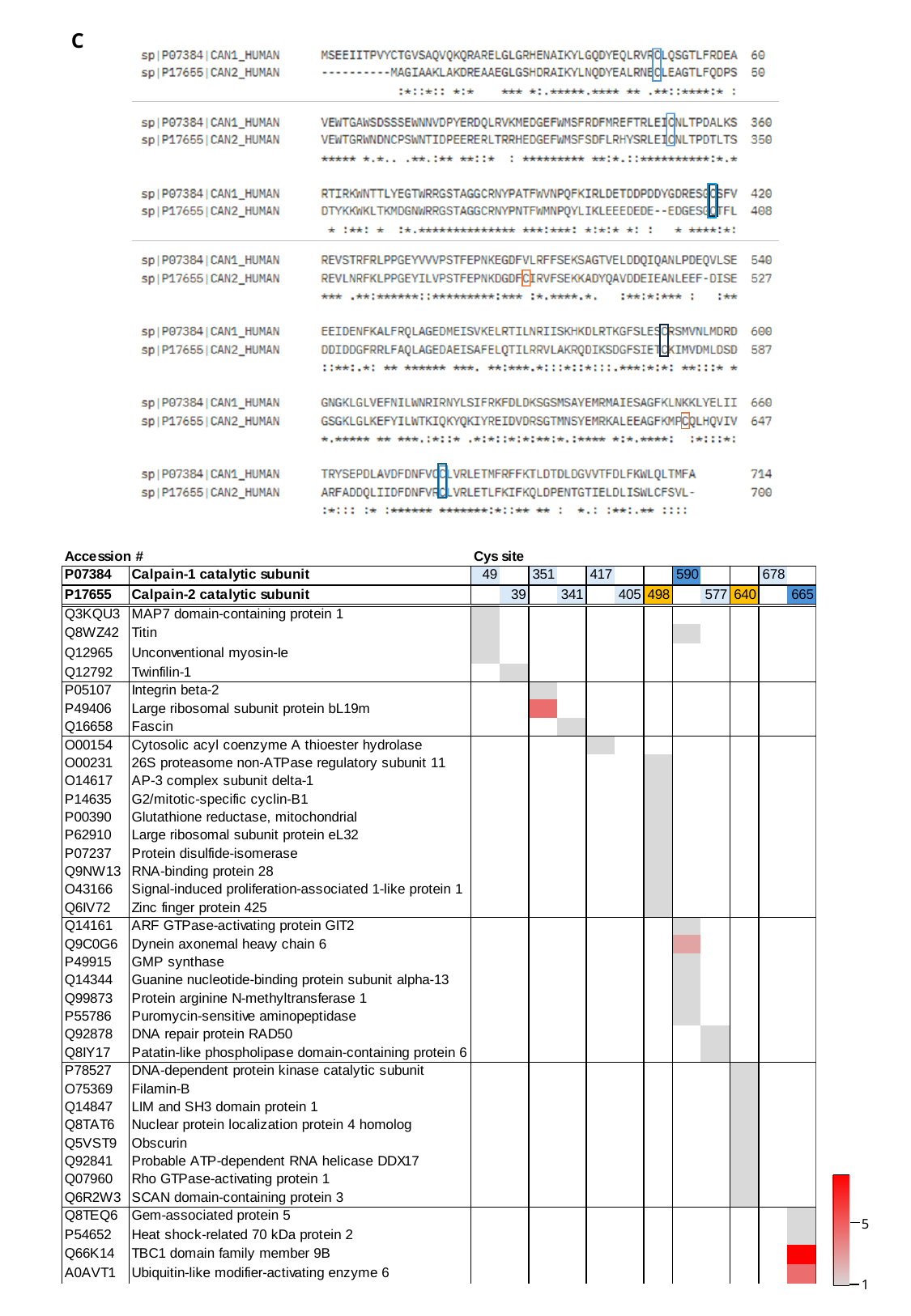

C
5
1

### Slide 42
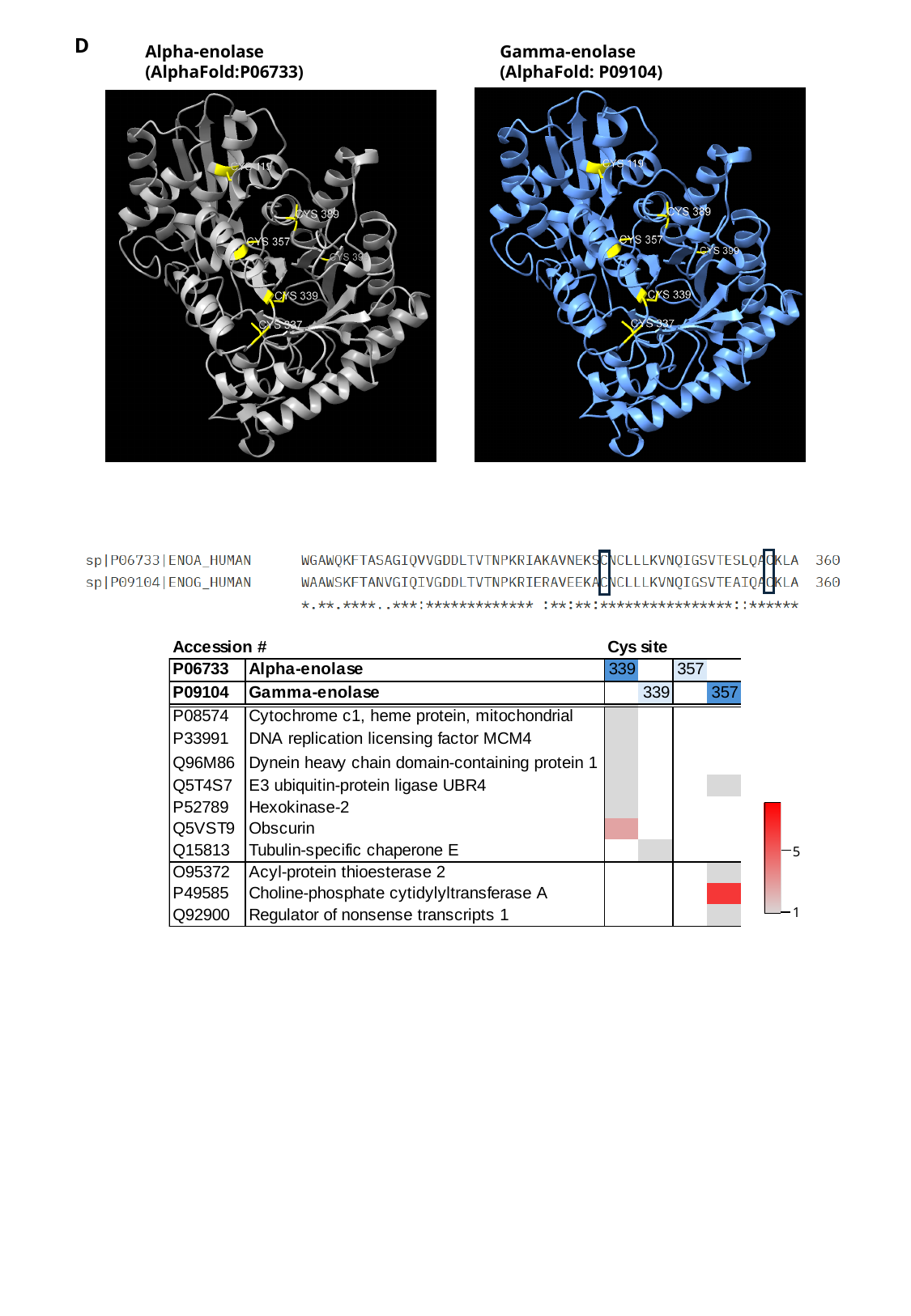

D
Alpha-enolase
(AlphaFold:P06733)
Gamma-enolase
(AlphaFold: P09104)
5
1

### Slide 43
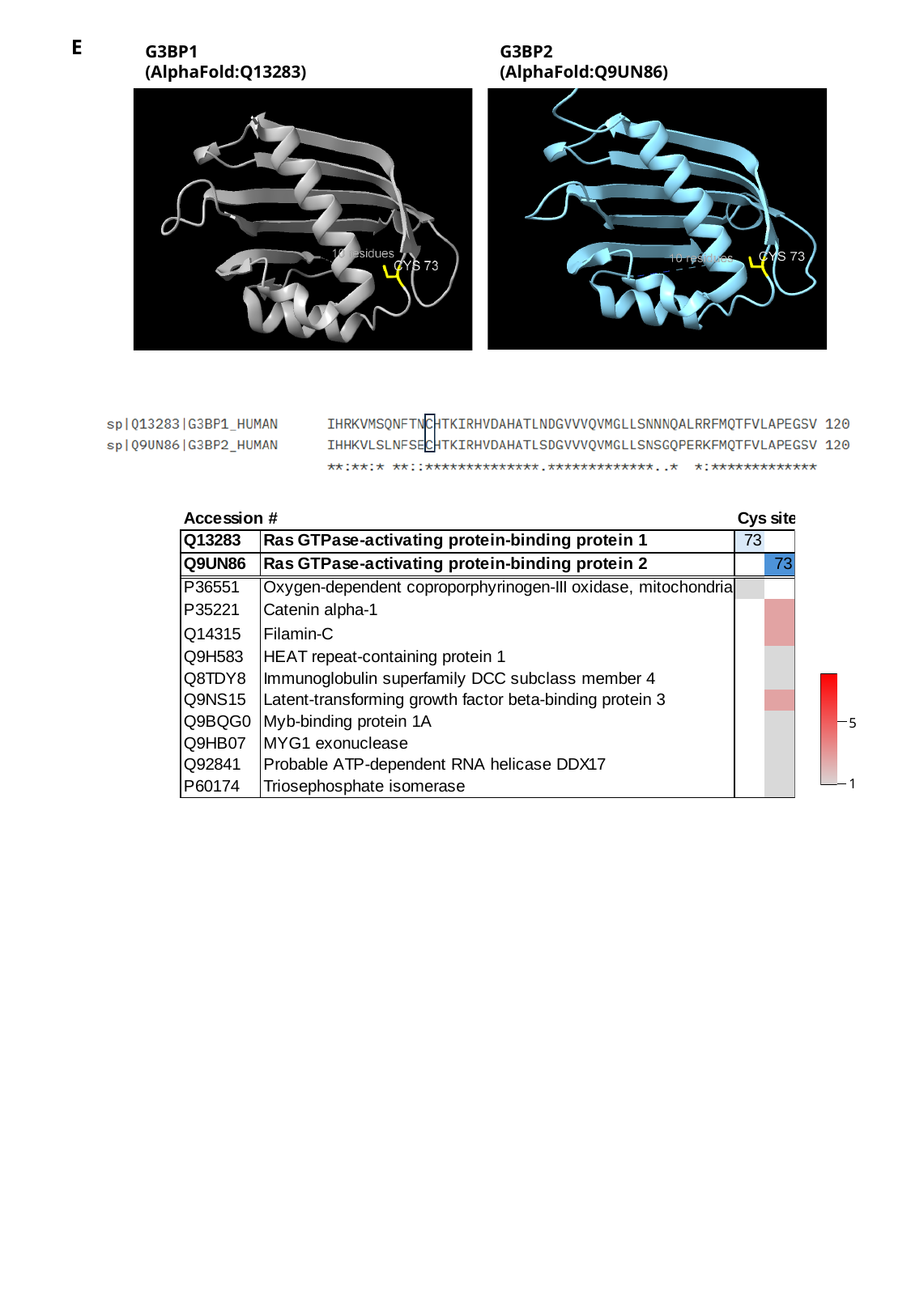

E
G3BP1
(AlphaFold:Q13283)
G3BP2
(AlphaFold:Q9UN86)
5
1

### Slide 44
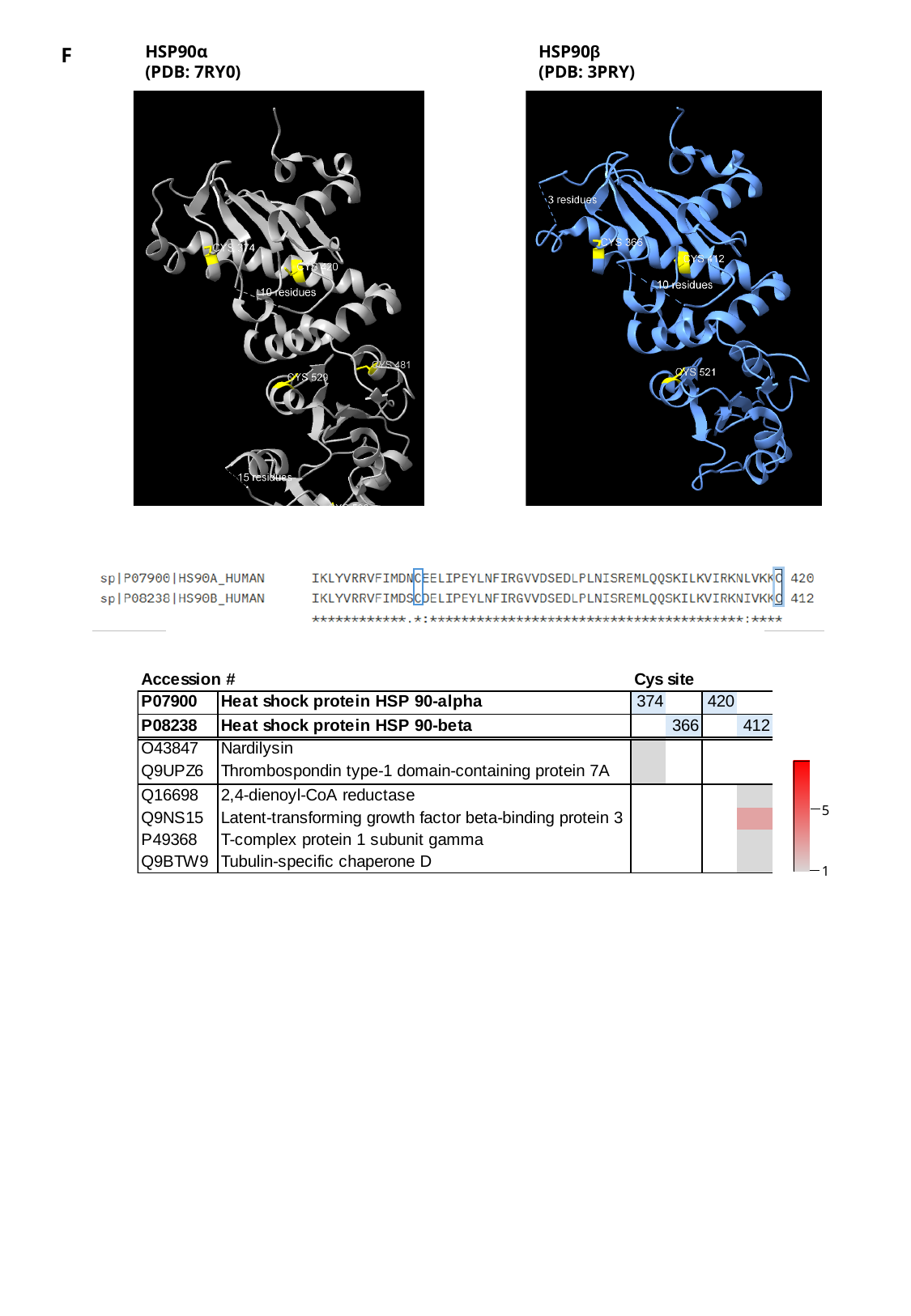

HSP90α
(PDB: 7RY0)
HSP90β
(PDB: 3PRY)
F
5
1

### Slide 45
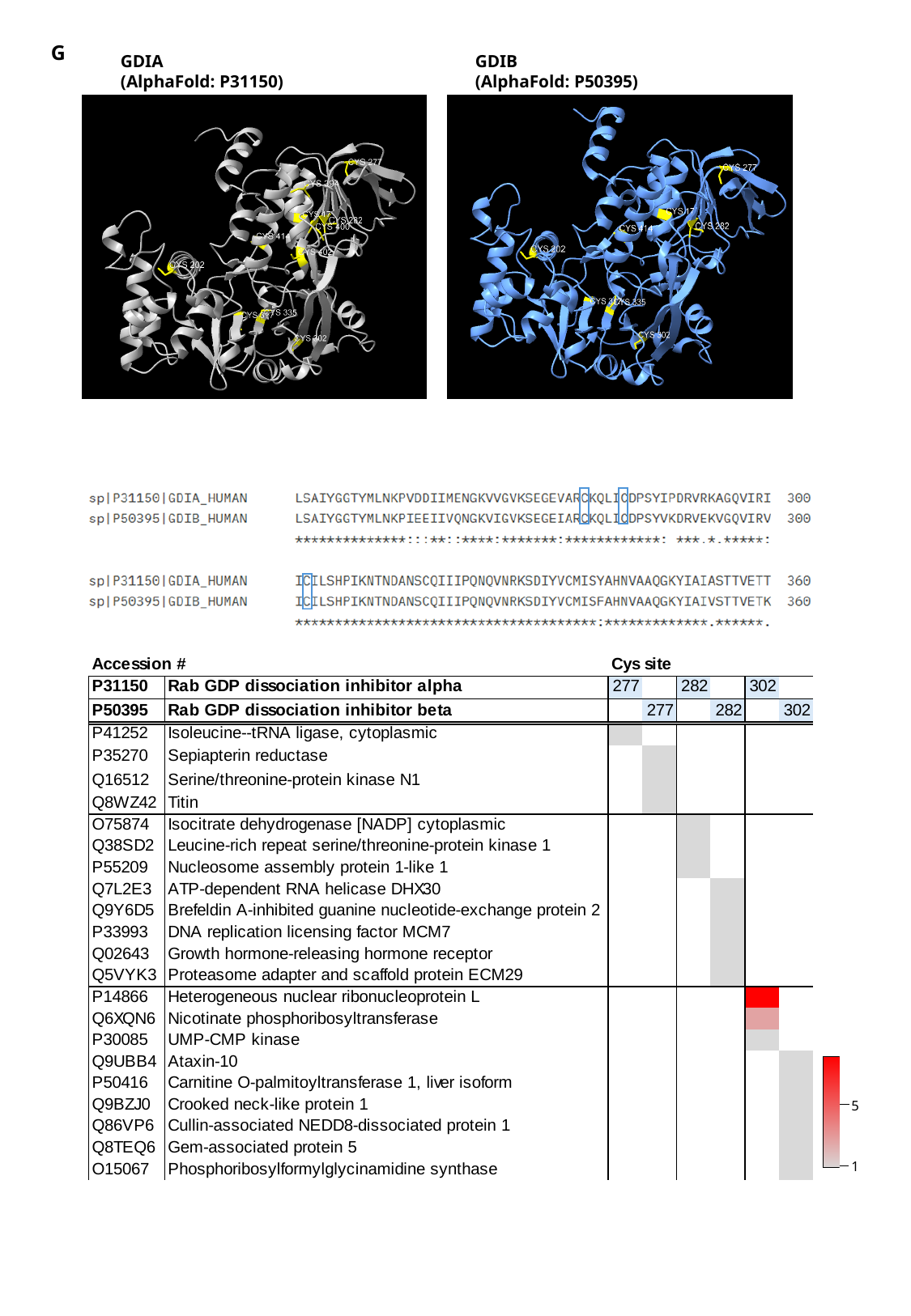

G
GDIA
(AlphaFold: P31150)
GDIB
(AlphaFold: P50395)
5
1

### Slide 46
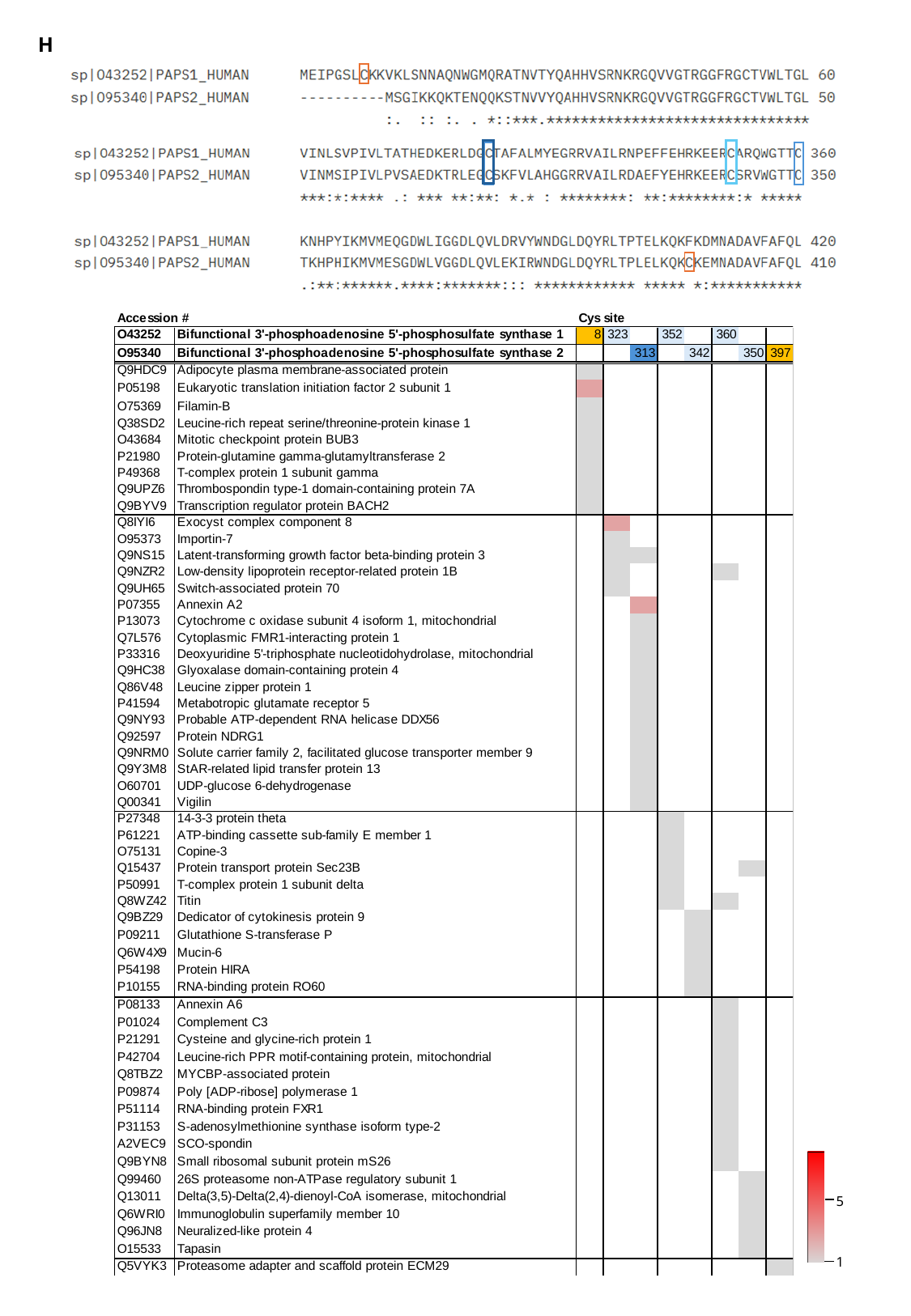

H
5
1

### Slide 47
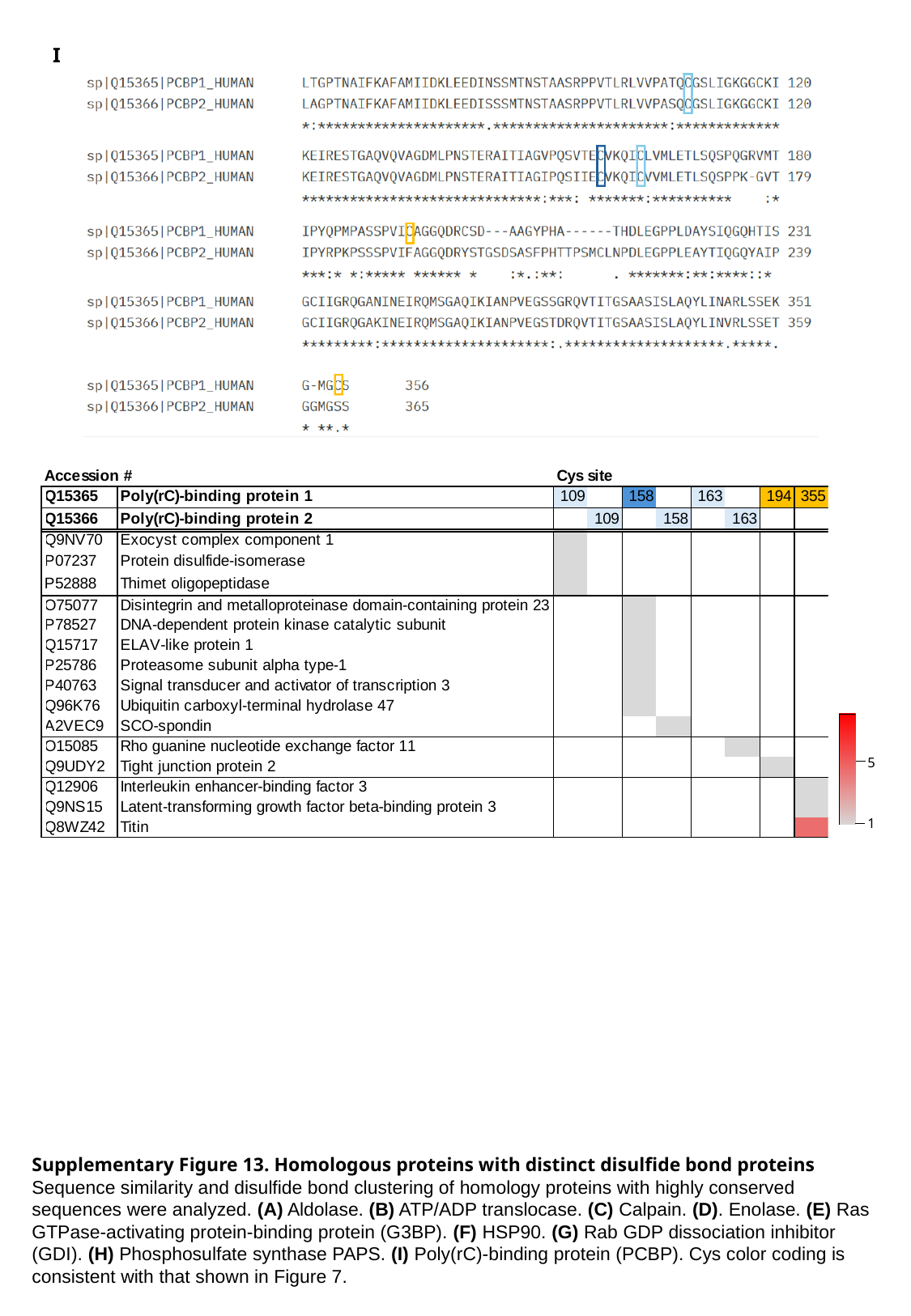

I
5
1
Supplementary Figure 13. Homologous proteins with distinct disulfide bond proteins Sequence similarity and disulfide bond clustering of homology proteins with highly conserved sequences were analyzed. (A) Aldolase. (B) ATP/ADP translocase. (C) Calpain. (D). Enolase. (E) Ras GTPase-activating protein-binding protein (G3BP). (F) HSP90. (G) Rab GDP dissociation inhibitor (GDI). (H) Phosphosulfate synthase PAPS. (I) Poly(rC)-binding protein (PCBP). Cys color coding is consistent with that shown in Figure 7.

### Slide 48
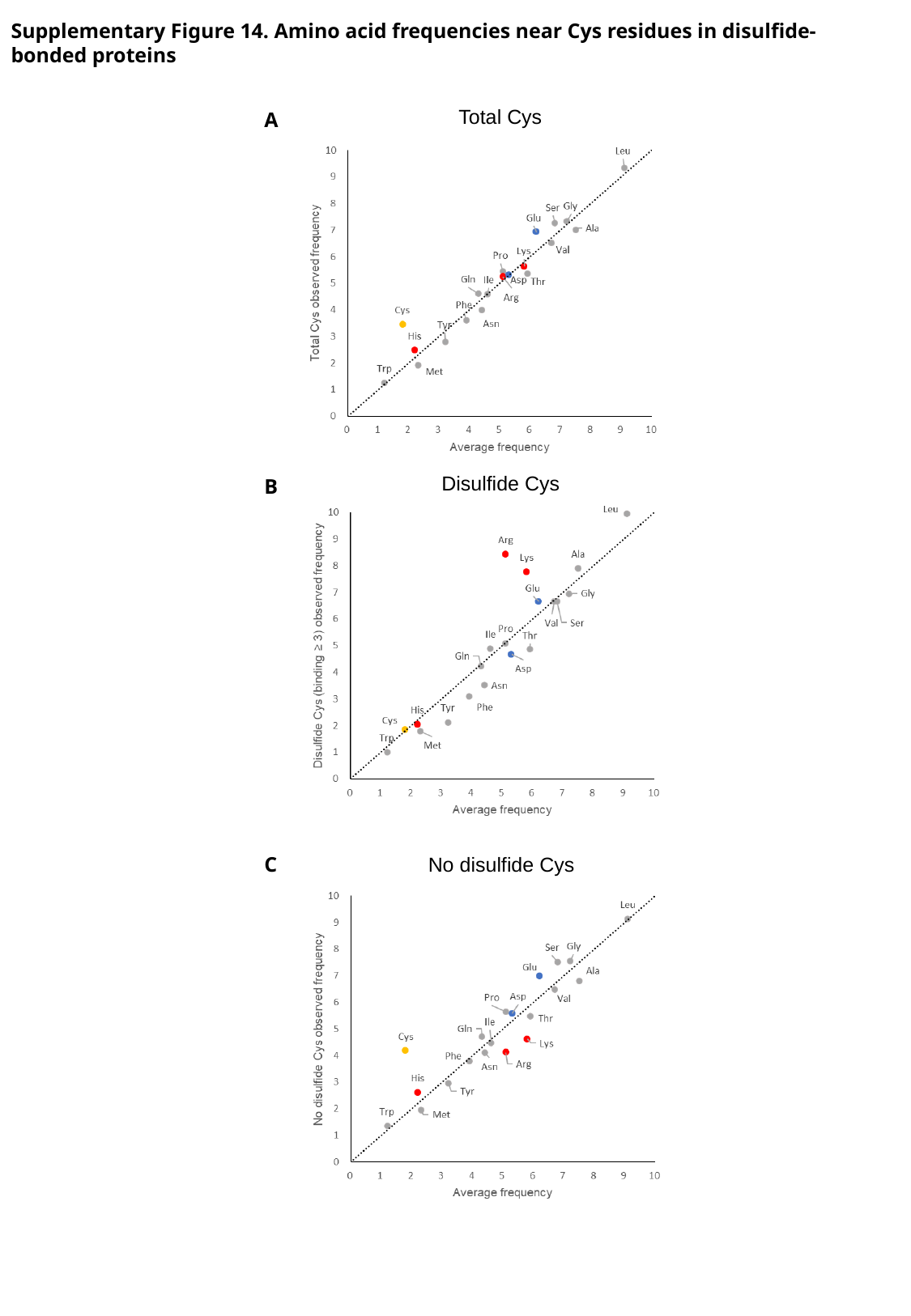

Supplementary Figure 14. Amino acid frequencies near Cys residues in disulfide-bonded proteins
Total Cys
A
Disulfide Cys
B
C
No disulfide Cys

### Slide 49
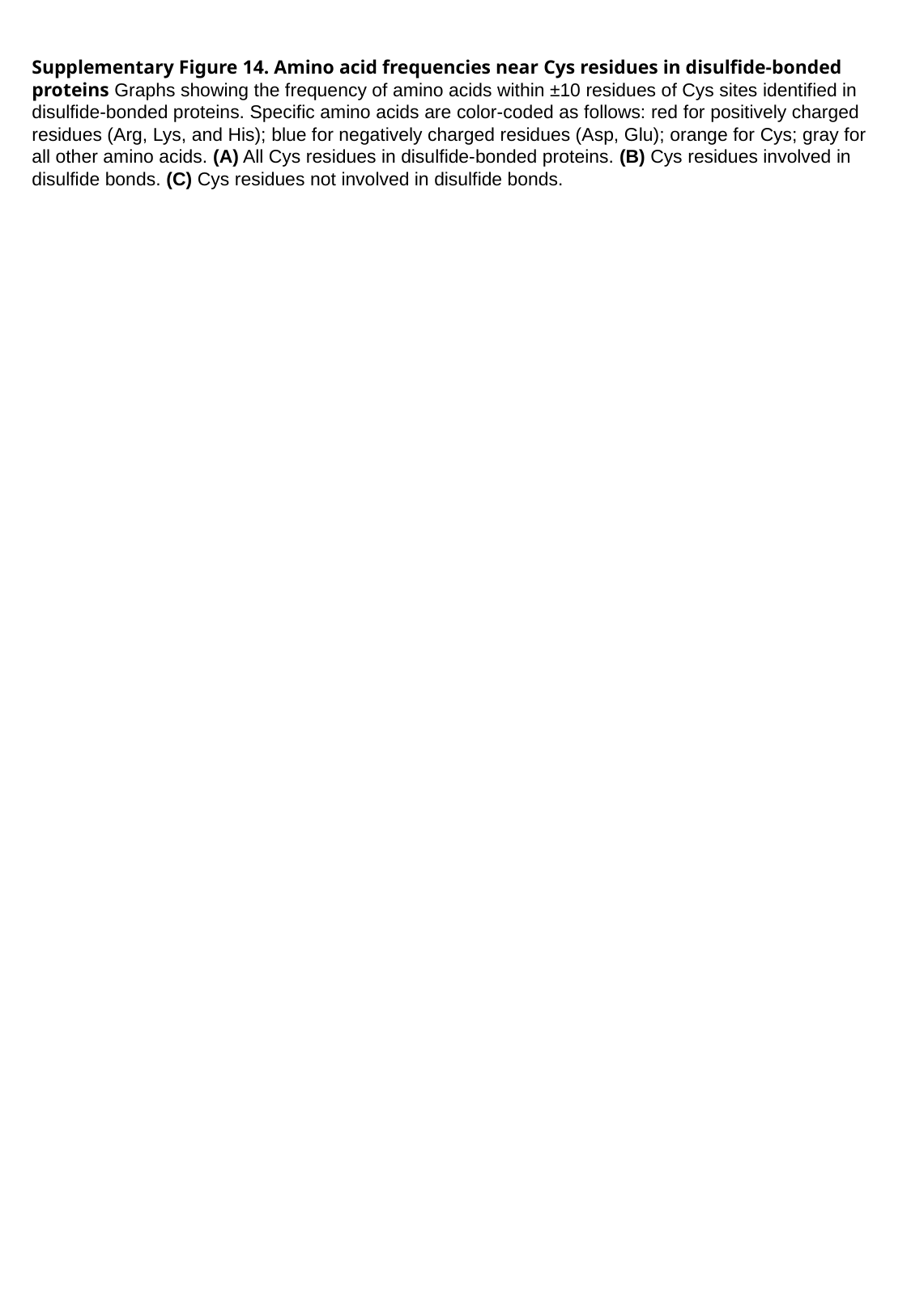

Supplementary Figure 14. Amino acid frequencies near Cys residues in disulfide-bonded proteins Graphs showing the frequency of amino acids within ±10 residues of Cys sites identified in disulfide-bonded proteins. Specific amino acids are color-coded as follows: red for positively charged residues (Arg, Lys, and His); blue for negatively charged residues (Asp, Glu); orange for Cys; gray for all other amino acids. (A) All Cys residues in disulfide-bonded proteins. (B) Cys residues involved in disulfide bonds. (C) Cys residues not involved in disulfide bonds.
